# Ffp1, an ancestral *Porphyromonas* spp. fimbrillin

**DOI:** 10.1101/2023.12.08.570808

**Authors:** Luis Acuña-Amador, Frédérique Barloy-Hubler

## Abstract

**Background:** Little is known about fimbriae in the *Porphyromonas* genus. Besides *fim* and *mfa*, a third *Porphyromonas gingivalis* adhesin called Ffp1 has been described, and seems to be capital for outer membrane vesicle (OMV) production.

**Objective:** We aimed to investigate the distribution and diversity of type V fibrillin, particularly Ffp1, in the *Porphyromonas* genus.

**Methods:** A bioinformatic phylogenomic analysis was conducted using all accessible *Porphyromonas* genomes in order to generate a domain search for fimbriae, using HMM profiles.

**Results:** Ffp1 was found as the sole fimbrillin in all the analyzed genomes. After manual biocuration and 3D modeling, this protein was determined to be a type V fimbrillin, with a closer structural resemblance to a *Bacteroides ovatus* fimbrillin than to FimA or Mfa1 from *P. gingivalis*.

**Conclusion:** It appears that Ffp1 represents ancestral fimbriae present in all *Porphyromonas* species. Additional investigations are necessary to elucidate the biogenesis of Ffp1 fimbriae and his potential role in OMV production and niche adaptation.

## Introduction

Fimbriae (fibrillae or pili) are adhesins consisting of protein polymers forming filamentous appendages that protrude from the bacterial cell surface. Unlike motility flagella, fimbriae have adhesive properties to attach to surfaces. In Gram-negative bacteria, fimbriae are classified according to their assembly pathways, including the chaperone-usher (CU) pilus system, the type IV pilus, and the conjugative type IV secretion pilus (1,2). In 2016, a new prevalent type V pilus was discovered within the human gut microbiome (3) and was described as a new donor strand-mediated system restricted to the Bacteroidia class (2). This system resembles the CU type, but requires the lipoprotein sorting pathway, and outer membrane proteinases (4).

Type V fimbriae have been mainly studied in *P. gingivalis* which classically produces two distinct adhesins, termed FimA (described in 1984 (5)), and Mfa1 (described in 1996 (6)), according to the names of stalk subunits (7). Both stalk proteins must be processed and matured. They possess long leader peptides (8) that facilitate their transport to the periplasm via the Sec system. Subsequently, they undergo lipid modification and are cleaved by type II signal peptidase (9), followed by a proteolytic maturation achieved by RgpA, RgpB and Kgp proteinases called gingipains (10). Finally, mature fibrillin monomers polymerize (11). The genetic loci for both fimbriae are distinct but organized into two clusters: *fimA-E* and *mfa1-5* (12).

In 2017, a third *P. gingivalis* adhesin was described (PGN_1808 in the ATCC 33277 strain or PG1881 in the W83 strain) and termed Ffp1 for filament-forming protein 1 (13). It corresponds to filaments 200 to 400 nm in length and 2 to 3 nm in diameter, that can be degraded, unlike FimA or Mfa1, by detergents and temperature into 50 kDa monomers (14). Ffp1 is among the exclusive repertoire of proteins within the order Bacteroidales and is conserved across *Porphyromonas* and *Bacteroides* (15,16). This protein was identified among the outer membrane proteins and especially the O-glycoproteome of *P. gingivali*s (17) and was described as essential in the production of outer membrane vesicles (OMVs), as the Ffp1 null-mutants exhibited a 30% reduction in OMVs production compared to the wild-type strain (13). Moreover, a recent study indicates a connection between Ffp1 and the production of sphingolipids (SL). In the absence of SL, *P. gingivalis* generates OMVs without Ffp1, whereas OMVs containing SLs exhibit an enrichment of Ffp1. Interestingly, these SL-containing OMVs limit host inflammation (18).

Ffp1 C-terminal region is homologous to type IV fimbriae from *Bacillus* spp. (15) and its sequence bears a significant similarity to the adhesion protein BACOVA_01548 (PDB ID: 4rfj) from *Bacteroides ovatus* (3). Its structural modeling suggests a donor strand-mediated assembly mechanism (14), which would classify Ffp1 as a new type V pilin (13). However, unlike FimA or Mfa1, no accessory component has yet been identified for Ffp1 despite its apparent co-expression as an operon with three upstream genes, annotated as a Cys-RNAt ligase, a patatin (lipase) and a glycosyl transferase. This co-transcription suggests the involvement of these four proteins in the same biochemical pathway or utilization of the same substrates/transporters, albeit without physical interaction (14).

To date, Ffp1 has been the subject of few works limited to *P. gingivalis*, only on two reference strains ATCC33277 and W83, and no information is available for the other 21 *Porphyromonas* species. At the genus level, knowledge for *non-P. gingivalis* Ffp1 or other fimbriae is scarce, with the exception of description of FimA-like and Mfa1-like fimbriae in *P. gulae*, a closely related species to *P. gingivalis* (19,20), and reports indicating fimbriation in *P. circumdentaria*, *P. macacae* and *P. asaccharolytica* (21–23), without further characterizations.

In this context, the aim of this study is to complete this knowledge gap and to investigate the distribution and diversity of type V fibrillin, particularly Ffp1, in the *Porphyromonas* genus. To do so, we performed an *in silico* analysis of type V fimbrillin locus in all 144 available genomes of *Porphyromonas*, investigating their presence/absence and then focus on Ffp1 diversity, and 3D predicted structure.

## Material and Methods

### Porphyromonas taxogenomics

All 144 *Porphyromonas* genomes (Table S1) were automatically downloaded from the NCBI RefSeq database using the ncbi-genome-download script^*^. Unannotated Metagenome-Assembled Genomes (MAGs) with inconsistent taxonomic labels were not considered. To categorize all genomes into reliable groups, genomic data-driven taxonomic confirmation and/or assignment were performed. To confirm the assignment of genomes with a species name, we conducted a comparison of three metrics : i) the 16S rRNA gene percentage identity (when annotated), evaluated using a threshold of 98.65% (24); ii) the digital DNA-DNA hybridization distance (DDH) using the GGDC v2.1 (25) and ggdc-robot script^*^, with the default threshold of 70% using formula 2 (25–27); and iii) the whole genome Average Nucleotide Identity (gANI), calculated using FastANI with a threshold of 96% for species demarcation (28). In case of a disagreement between these three metrics, we combined alignment fraction values (AF) with gANI using 60% and 96.5 % as threshold values respectively, to assign a genome pair to the same species (29). Additionally, when needed, we also used OrthoANI^‡^ to measure and visualize the overall similarity between some *Porphyromonas* species.

For the genomes without a specified species name (*Porphyromonas* sp.), as the majority of them originated from environmental samples (human- or animal-associated habitats) and are often highly fragmented, it was crucial to ensure that they were not contaminated and do not correspond to genome assemblies containing a mixture of different species. This genomic homogeneity was evaluated with Kraken2 (30) using the non-redundant nucleic database (updated April 22). Only assemblies that consisted of over 80% of *Porphyromonas* content and/or larger than 80% of the expected average genome size (2.5 Mb) were retained for our analysis. Their affiliation to the *Porphyromonas* genus was first confirmed using fIDBAC server^§^ (31) and their position within the *Porphyromonas* taxonomy was validated using an Orthofinder rooted species tree (32). This tree was construted using all *Porphyromonas* sp. (*P.* sp.) and one reference genome per *Porphyromonas* species (refer to Table S1) and was visualized using FigTree^**^. For each branch, one or several *P. sp* were associated to a *Porphyromonas* species through ANI and DDH, employing the same thresholds as previously described.

### *Porphyromonas* fimbriae identification and classification

#### 1. Dataset construction

Sequences from type V fimbriae (FimABCDE, Mfa12345 and Ffp1) were manually extracted from the 59 *P. gingivalis* genomes, and were used as queries to identify homologous sequences all in the genomes of other *Porphyromonas* spp. using BlastP (identity ≥ 30%; query coverage ≥ 60%; e-value < 10e^-5^). All sequences were grouped as dataset 1.

#### 2. Functional domain-based screening

Dataset 1 was subjected to analysis using InterProScan to identify all protein domains associated with those sequences. The resulting domains were searched in the complete orfeomes of *Porphyromonas* downloaded from PATRIC 3.6.6 (33), using HMMsearch from HMMER v3.3.1 (34) and the hidden Markov models (HMMs) from Pfam 33.1 (May 2020) database (35). Sequences harboring the targeted domains with an e-value < 10e^-06^ were retained and grouped into dataset 2.

#### 3. Protein clustering, biocuration and HMM profile construction

Dataset 2 was clustered with MMseqs2 (36) via the easy-cluster command. Each cluster obtained underwent manual biocuration after multiple alignment using Clustal Omega (37) and any missing genes were annotated. Subsequently, for each cluster, the multiple alignments were converted from FASTA format to Stockholm format with ‘sreformat’ command and HMM profiles were generated using the ‘hmmbuild’ command with default settings. Clustering and HMM profiles creation was first performed on raw data and then refined on biocurated data.

#### 4. Final classification

The obtained HMM profiles (refer to Supplementary material) were used to identify and classify all fimbrillins within the *Porphyromonas* orfeomes, downloaded from PATRIC 3.6.6 database, using ‘hmmsearch’ command from HMMER package.

#### 5. *In silico* analysis of *Porphyromonas* fimbrillins

Geneious Prime (38) was used to visualize the genomic context of each identified fimbrillin. Biocuration for start codon were proposed, based on sequence homology, to optimize the prediction of SPII signal peptide and the cleavage site positions. N-terminal region was identified using *charge* (window size=3) from EMBOSS 6.6.0 (39), the H hydrophobic region was characterized with Kyte-Doolittle hydropathy plot made with ProtScale (40) (window size=3), and the cleavage site was confirmed by SignalP 6.0 (41) and LipoP (42). Palmitoylation in the lipobox cysteine residue was verified using CSS-Palm (43). Protein sizes were represented using violin plots (geom_violin) and/or boxplot (geom_box), both functions from the ggplot2 package (44).

For each fimbrillin family, a multiple alignment was performed using MAFFT (L-INS-I algorithm and BLOSUM62 matrix; gap open penalty and offset value by default) (45). This alignment was visualized in two dimensions using Alignmentviewer v1.1^††^ which employs the UMAP algorithm (46) and Hamming distance to cluster aligned sequences. Phylogenetic trees were calculated using FastTree 2.1.11 (47), PhyML 3.3 (48), and RaxML (49) with default parameters.

The taxonomic distribution of fimbrillin genes was analyzed across a phylogenetic tree constructed using OrthoFinder based on the pangenomes of all confirmed *Porphyromonas* species groups (32) and visualize using FigTree. The phylogenetic reconstruction was performed both using native and mature proteins (i.e excluding their signal peptides) using RaxML (evolution model GAMMA LG and 100 bootstrap). Robinson-Foulds, Nye Similarity and Jaccard Robinson Foulds distances between the phylogenetic trees were calculated using TreeDist^‡‡^ R library and tanglegrams were created with the R package phytools^§§^ (scripts TREE.R and Tanglegram.R).

#### 6. 3D modeling

Secondary protein structure was predicted with PSIPRED in Phyre2 (50).

3D structures of Ffp1 mature proteins were modeled, based on homology modeling, using Robetta (51) and the RoseTTAFold method, as well as Phyre2. The quality of all five 3D models generated by Robetta for each Ffp1 protein was assessed and validated using two quality calculation tools: ERRAT (52) and Verify3D (53). The most accurate predicted structure was chosen and superposed to the best model target, found by VAST+ (54), Phyre2 and iPBA (55). The RMSD value (56) as well as the number and percentage of aligned residues were retrieved and compared to Phyre2 results. RMSD < 3Å were considered significant between Ffp1 predicted structure and 3D models (57).

## Results

### 1. *Porphyromonas* taxogenomic assignment

The 144 *Porphyromonas* genomes studied in this work (Table S1) were predominantly in draft form (85% of the genomes), with only 6 out of the 17 analyzed species possessing at least one complete genome.

The taxogenomic assignment for the genomes classified into the 17 *Porphyromonas* species was verified (Table S1). Species *P. loveana* and *P. pasteri* have only one representative genome and therefore cannot be verified intra-specifically. For the other species, intra-specific analysis combining ANI, 16s rRNA and DDH comparison (Figure 1A) showed no anomalies for taxonomic placements, except for *P. uenonis*, *P. somerae*, and *P. canoris*.

**Figure 1.**
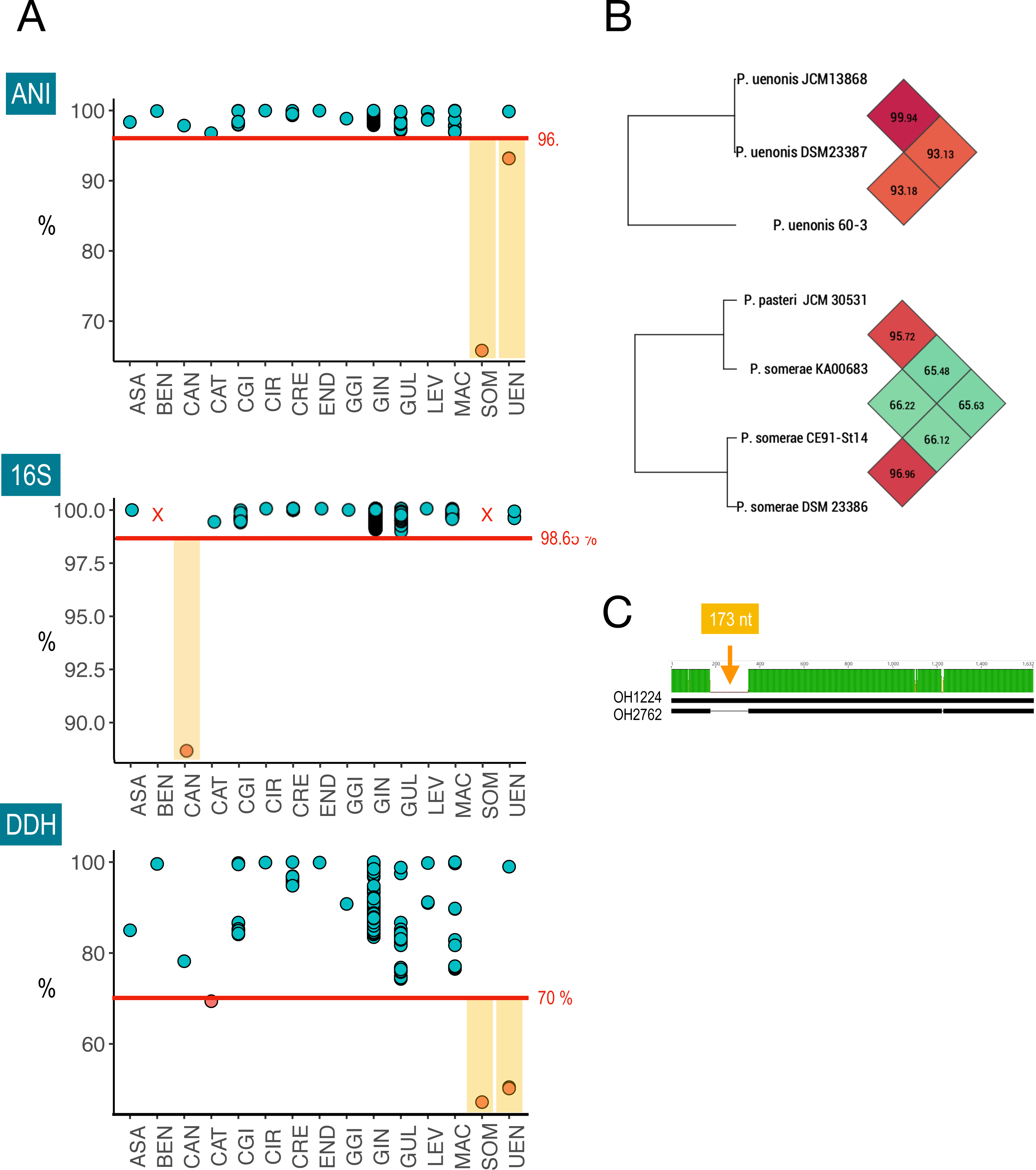
Validation of the taxogenomic assignment of *Porphyromonas* genomes. **A.** Intra-species homogeneity was checked by calculating intra-species distances using gANI, rRNA 16S identity and DDH. **B.** Checking *P. uenonis* and *P. somerae* genomes homogeneity using OrthoANI. **C.** Difference in 16S rRNA sequences of two strains of *P. canoris* with 173 nt insertion in strain OH1224. 3-letter code acronyms correspond to ASA: *P. asaccharolytica*; BEN: *P. bennonis*; CAN: *P. canoris*; CAT: *P. catoniae*; CGI: *P. cangingivalis*; CIR: *P. circumdentaria*; CRE: *P. crevioricanis*; END: *P. endodontalis*; GGI: *P. gingivicanis*; GIN: *P. gingivalis*; GUL: *P. gulae*; LEV: *P. levii*; LOV: *P. loveana*; MAC: *P. macacae*; PAS: *P. pasteri*; SOM: *P. somerae*; and UEN: *P. uenonis*.

Firstly, for *P. uenonis*, the differences in metrics reflect a significant distance between strain 60-3 and the two other strains (Figure 1A and 1B). In fact, strain 60-3 rRNA operon is found within a single contig (6019 nt) that does not contain any other genes. Given that *P. uenonis* strain 60-3 was isolated from a human metagenome (vagina) and exhibits a highly fragmented genome, its 250 contigs underwent analysis using Kraken2 (Figure S1). Of the 238 contigs classified by Kraken2 (95.2% of total contigs), 78% corresponds to *Porphyromonas*, totaling 2.1 Mb, which is approximately 85% of the expected average size. It was concluded that *P. uenonis* 60-3 belongs to the *Porphyromonas* genus but its classification within the *P. uenonis* species appears to be incorrect based on ANI/DDH analysis. This genome has been retained for the study but as an unclassified *Porphyromonas*, denoted as PSP_60-3 (Table S1).

Secondly, in the case of *P. somerae* KA00683, all indicators suggest a taxonomic assignment inaccuracy. BlastN analysis of the 16S gene fragment (852 nt) reveals a 100% identity with *P. pasteri*, in accordance with ANI (95.7) and DDH (64.53) values, even though the latter two values are slightly below the established threshold (Figure 1B). However, the Kraken2 analysis indicates a genomic mixture, with only 32.5% of the reads being attributed to *Porphyromonas*. Consequently, we have opted not to include *P. somerae* KA00683 in our study, leaving only two strains within this species (Table S1).

Finally, regarding *P. canoris* (2 genomes), the difference in the 16S rDNA sequences was associated with an additional 173-nt fragment in strain OH1224, resulting in a longer gene (Figure 1C). However, it’s worth noting that these genomes are in draft and fragmented into 14 and 21 contigs. As a result, it is impossible to determine whether this difference represents genuine genomic diversity or a sequencing error. Nevertheless, since all other indicators (ANI, DDH and orthology) confirmed the uniformity of this species, we disregarded this 16S rRNA disparity and consider both genomes as belonging to *P. canoris*.

Furthermore, 28 *Porphyromonas* genomes lacked a species label, necessitating a multi-stage analysis. Initially, the genomic contents of these strains were examined using Kraken2, and genomes with less than 80% of *Porphyromonas* reads and/or that reconstructed less than 80% of *Porphyromonas* average genome size (2.5 Mb) were excluded from the study (Figure S1 and Table S1). Consequently, 17 strains were omitted from this study (Table S1). Among the 11 remaining *Porphyromonas* sp., their placement in the Orthofinder species tree based on ANI/DDH metrics (Figure 2) allowed to assign *P.* sp. OH4946 to species *P. gulae*; *P.* sp. MGYG-HGUT-04267 to species *P. asaccharolytica*; *P.* sp. UMGS1452 to species *P. uenonis* and *P.* sp. OH1349 and OH2963 to species *P. canoris* (Table S1). Finally, there were 6 *P.* sp. genomes that could not be assigned to any specific group and were individually examined (unassigned, Table S1). This examination further supported the reclassification of *P*. *uenonis* 60-3 as PSP_60-3 (Figure 2 and Table S1).

**Figure 2.**
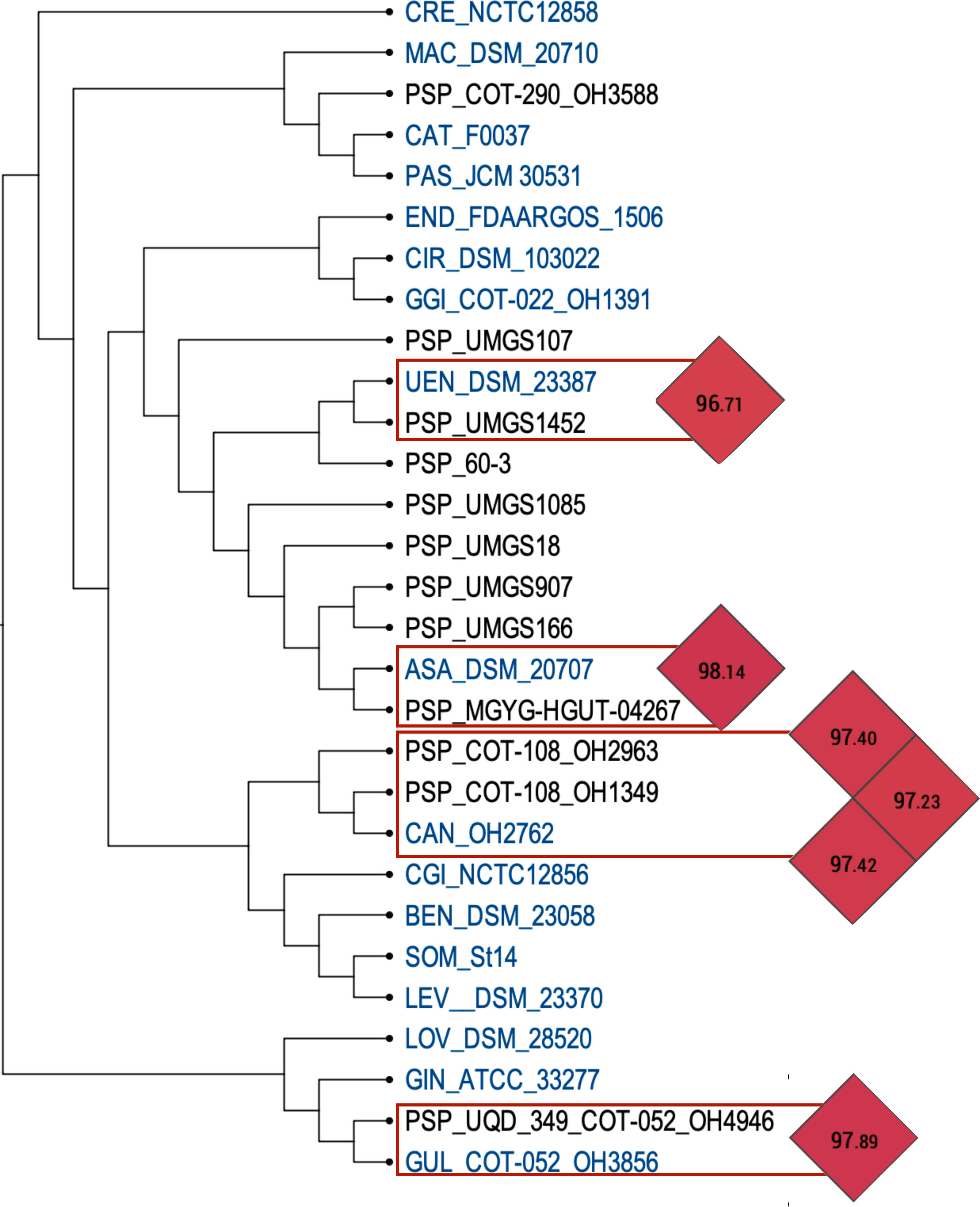
Phylogenetic species tree derived from OrthoFinder analysis. This tree was used to place some *Porphyromonas* spp. into UEN (UMGS1452), ASA (MGYG-HGUT-0467), CAN (OH2963 and OH1349) and GUL (OH4946) genus, after confirmation via OrthoANI.

After completing this taxogenomic biocurated analysis, our study retains a total of 126 *Porphyromonas* genomes clustered into 24 groups (comprising 17 species and 7 *P.* sp. singletons), unequally distributed between the genus, ranging from 59 genomes for *P. gingivalis* (almost half of all available genomes in the genus) to just one genome for *P. loveana*, *P. pasteri* and each *Porphyromonas* sp. (PSP).

### 2. FFp1 is the only fimbrillin common to all *Porphyromonas*

Screening and clustering fimbrillin genes from *Porphyromonas* genomes resulted in the definition of 12 HHM profiles, one for each gene in either FimABCDE or Mfa12345, and two for Ffp1. Searching for sequence similarity in each *Porphyromonas* orfeome, using each of the 12 HHM profiles, enabled the identification and classification of these three fimbriae systems in all *Porphyromonas* genomes (Figure 3).

**Figure 3.**
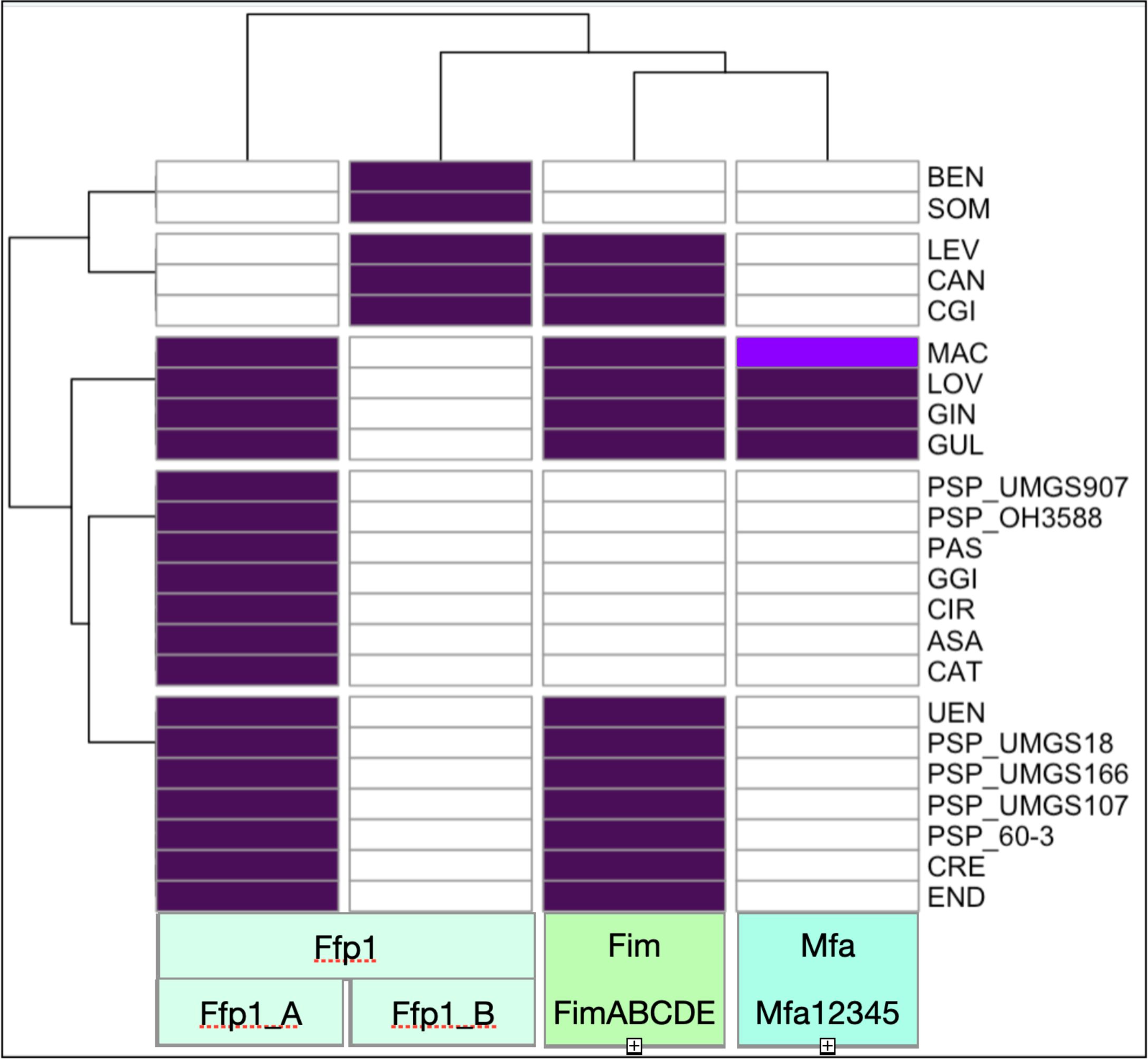
Heatmap depicting the presence/absence of fimbrillins. The heatmap scale color indicates whether fimbriae systems (FimABCDE, Mfa12345 or Ffp1_A or B) were detected: white (absence), dark purple (presence as one *locus*) and light purple (presence as two *loci*).

#### 2.1. *fimABCDE* locus

For the FimABCDE proteins (Figure S2A), an expect value (E-value) calibration was performed and set to a minimum threshold of e^-100^ for each of the 5 profiles. Using this threshold, the detection of the locus *fimABCDE* exhibited both sensitivity and specificity, perfectly correlating with presence/absence of each gene.

In each genome, these genes are colocalized and organized into operons, with an average size of 7.3 kb. Out of all the genomes, two stand out as outliers: *P. gingivalis* A7436 due to an IS5 family transposase ISPg8 insertion in *fimC*, and *P. uenonis* UMGS1452 for which the locus remains incomplete because located at the end of a contig.

It is noteworthy that all *P. macacae* strains possess two complete *fimABCDE* loci, a unique feature in *Porphyromonas*. This duplication raises questions about the redundancy or functional complementarity of both *loci*, especially as *P. macacae* JCM15984 has a pseudogenized *fimE* in locus 1 and a pseudogenized *fimD* in locus 2.

The utilization of HMM profiles in our search strategy allows for the rapid and unambiguous identification and classification of fimbrial genes, even in cases with low mean amino acid percentage identities: 52.3% (FimA), 63.7 % (FimB), 56.7% (FimC), 48.2% (FimD) and 49.8% (FimE). Additionally, the annotations of FimABCDE proteins are inconsistent, with the majority being labeled as hypothetical proteins or simply categorized as fimbrial proteins without any additional characterization (Figure S2B). As such ontology searches are almost impossible.

Moreover, the establishment an E-value threshold facilitates pinpointing abnormalities.

For instance, in *P. gingivalis*, for the FimB HMM profile, the E-value is greater than the established threshold due to a nonsense mutation in *fimB* for the ATCC33277 strain (58), this gene is annotated as two genes (PGN_0181, e-value = 2.8e^-63^ and PGN_0182, e-value = 1.4e^-55^). The same case occurs in *P. uenonis*, for the FimE HMM profile, due to the incompleteness of this gene (at the end of contig) for the UMGS1452 strain.

In every analyzed *Porphyromonas* genome, the *fimABCDE* locus is consistently present, with only nine groups lacking this operon: *P. asaccharolytica*, *P. bennonis*, *P. catoniae*, *P. circumdentaria*, *P. gingivicanis*, *P. pasteri*, *P. somerae*, *P.* sp. OH3588 and *P.* sp. UMGS907.

#### 2.2. *mfa12345* locus

Significant E-values ranging from e^-200^ and e^-100^ were observed for each of the 5 Mfa12345 profiles (Figure S2C). Specifically, regarding the Mfa1 HMM profile, three distinct situations were evident: i. Mfa1 was recovered, with low E-values, in four species (*P. gingivalis*, *P. gulae*, *P. loveana* and *P. macacae*); ii. in 14 groups, Mfa1 was identified with higher E-values; and iii. in six species (*P. bennonis*, *P. canoris*, *P. cationae, P. cangingivalis* and *P. pasteri*, as well as PSP_OH3588), no Mfa1 was detected. The Mfa2 HMM profile produces identical results, yielding the same three groups.

The Mfa3 HMM profile successfully identified this protein in the same four species (*P. gingivalis*, *P. gulae*, *P. loveana* and *P. macacae*) and additionally in *P. endodontalis* that contains an Mfa3-like protein. Finally, both the Mfa4 and Mfa5 HMM profiles exclusively detected these proteins in *P. gingivalis*, *P. gulae*, and *P. loveana* and in three of the six strains of *P. macacae*: JCM15984 and NCTC11632 (isolated from the oral cavity of cats) and OH2859 (isolated from a canine oral cavity). In OH2859, the *mfa12345* operon locus is intact, while in the cases of JCM15984 and NCTC11632, we observed two distinct loci: the first one contains genes encoding Mfa123 proteins, followed by two genes encoding proteins similar to FimD and FimE (referred to as *mfa123_fimDE*) and the second comprises genes encoding Mfa2345 proteins preceded by a non-characterized fimbrilin gene that shares similarity with Ffp1, indicated by low E-values of 7.3e^-58^ for Ffp1 profile A and 7.8e^-41^ for Ffp1 profile B (referred to as *ffp1-like_mfa2345*). It is worth noting that three strains of *P. macacae*, specifically OH2631 (isolated from the canine oral cavity), as well as NCTC13100 and DSM20710/JCM13914 (isolated from the macaque oral cavity), exhibit two tandemly organized *mfa123_fimDE loci*. Remarkably, these loci are not identical, displaying an average sequence identity of 53%. In the case of OH2631, these *loci* are separated by less than 2 kb, while in NCTC13100 and DSM20710, they are separated by a 3 kb region that includes an IS4 pseudogene. None of these three strains harbor the *ffp1-like_mfa2345* locus.

*P. endodontalis* features an additional alternative locus comprising six genes, including Mfa1-like, Mfa2, Mfa3-like, followed by two genes encoding lipoproteins and one gene encoding a von Willebrand factor type A (VWA) domain-containing protein. Interestingly, several other species, such as *P. asaccharolytica*, *P. circumdentaria*, *P. crevioricanis*, *P. gingivicanis* and *P. uenonis*, also exhibit alternative *loci*, which likely correspond to novel fimbrilin systems. These systems require in-depth dedicated future studies for thorough characterization.

In conclusion, when considering only the complete *mfa12345* locus as a reference, we identified its presence in four species: *P. gingivalis*, *P. gulae*, *P. loveana*, and *P. macacae* strain OH2859. We also illustrate the effectiveness of HMM profiles in distinguish true *mfa loci* from alternative *loci*. As for FimABCDE, the descriptions found in the annotations of Mfa12345 proteins are uninformative, often annotated as hypothetical or fimbria. This labeling makes it nearly impossible to conduct meaningful ontology searches (Figure S2D).

#### 2.3. ffp1

MMseqs2 clustering reveals the separation of Ffp1 orthologs in two distinct groups which resulted in two distinct HMM profiles termed Ffp1_A and Ffp1_B (Figure S2E). Ffp1_A mature amino acid sequences, excluding the signal peptide, share a 57.4% identity, while Ffp1_B sequences exhibit only a 37% identity, primarily due to divergence in *P. bennonis.* The identity between the two groups decreases to 24%.

Ffp1_A HMM profile retrieves genes from all *Porphyromonas* species except for *P. bennonis*, *P. canoris*, *P. cangingivalis*, *P. levii*, and *P. somerae*, which are recovered with Ffp1_B HMM profile. So, remarkably, fimbrillin Ffp1 is indeed present in all *Porphyromonas* spp., contrary to FimABCDE and Mfa12345 (except for *P*. sp. UMGS1085 where a 186 nt fragment of a gene (at the start of a contig) is identified by Ffp1_A HMM profile with an E-value at 6.7 e-22 (Figure S2E). This higher E-value is the result of being obtained for only 61 amino acids instead of about 500 for a Ffp1_A protein.

As shown in figure Figure S2F, approximately 70% of the identified Ffp1 proteins are annotated as hypothetical or uncharacterized, 22% as fimbrillin/fimbriae (with half linked to the PGN_1808 protein, described as Ffp1 in the *P. gingivalis* ATCC33277 reference strain) and 8% are described as lipoproteins.

Using HMMsearch with both Ffp1_A and Ffp1_B profiles, using an E-value threshold at e^-100^, in Ensembl Genome Bacteria (taxid:2) database, only *Porphyromonas* proteins are retrieved. As a conclusion, Ffp1 fimbrillins are the sole fimbriae proteins conserved across all *Porphyromonas* species, making them unique to the genus.

### 3. Characterization of *Porphyromonas* FFp1 fimbriae

Ffp1 exhibits variable pre-cleavage sizes among *Porphyromonas* species, in both subclasses. For the Ffp1_A group, protein sizes range from 439 aa (*P*. *circumdentaria* DSM 103022) to 553 aa (*P. asaccharolytica* PR426713P-I), and for the Ffp2_B, from 483 aa (*P*. *somerae* DSM 23387) to 527 aa (*P*. *canoris*) (Figure 4A). Size is well conserved within *Porphyromonas* species with the exception of the *P*. *asaccharolytica*, *P*. *circumdentaria*, *P*. *macacae* and *P*. *uenonis* for Ffp1_A, and *P*. *bennonis* for Ffp1_B (Figure 4A).

**Figure 4.**
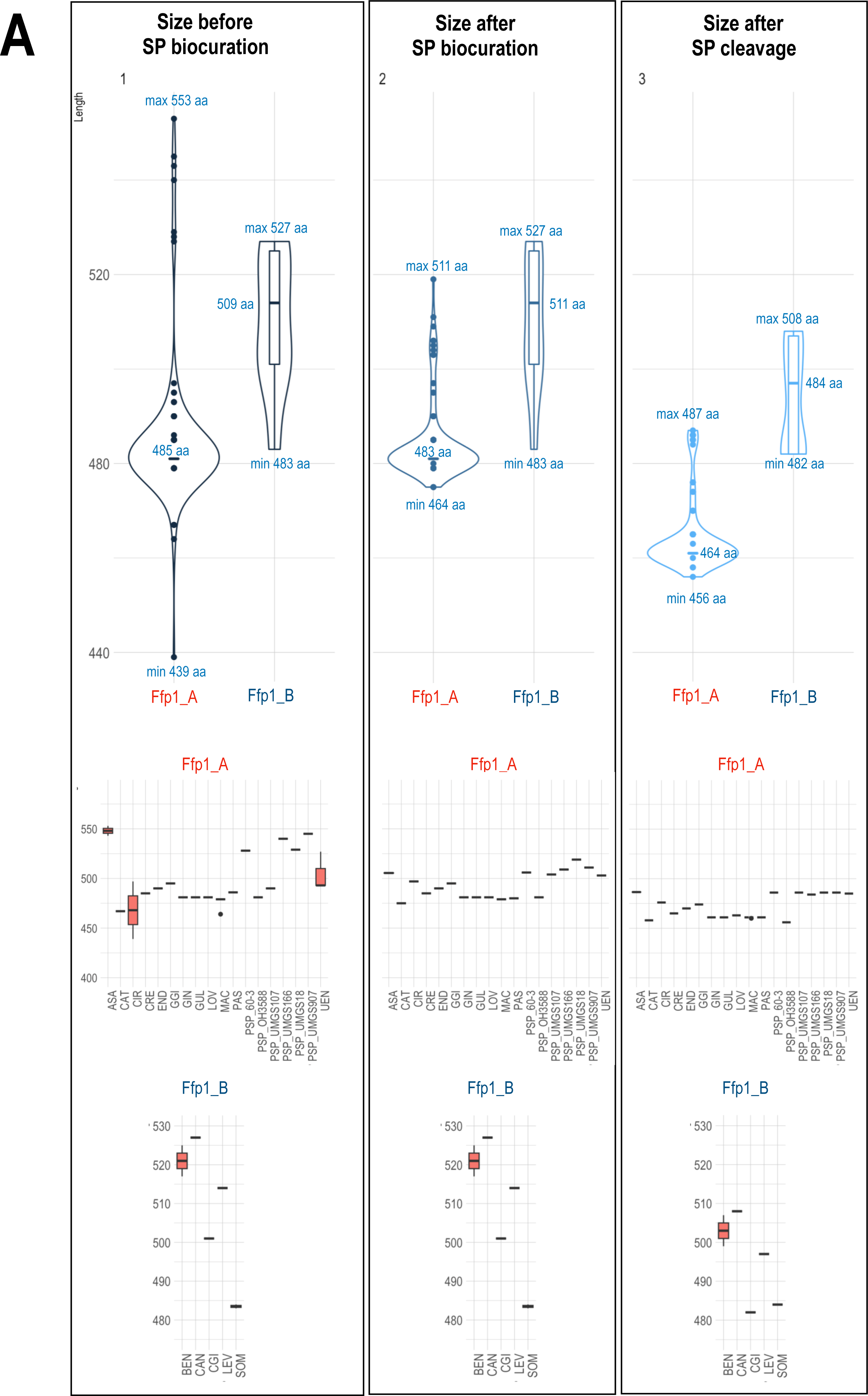
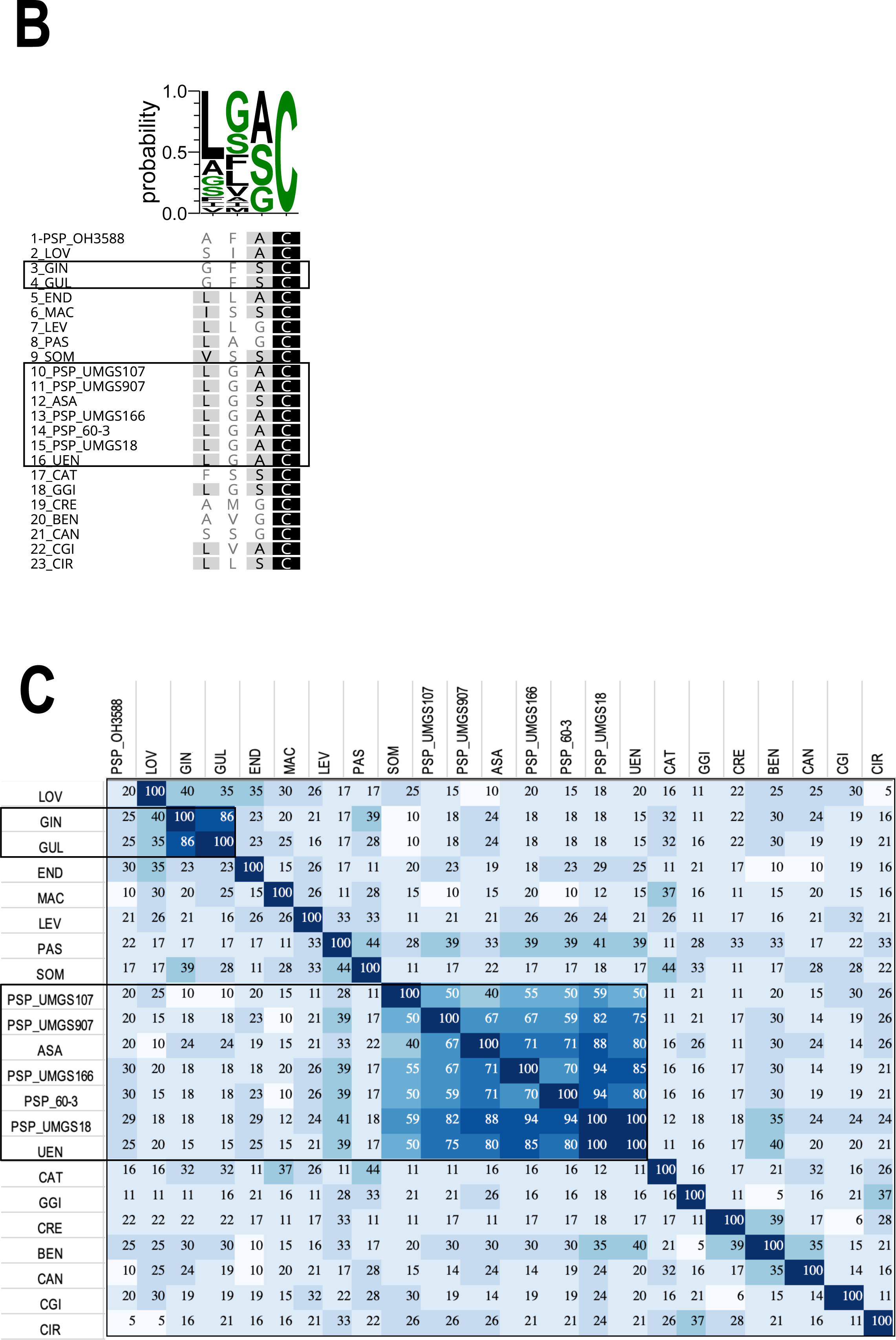
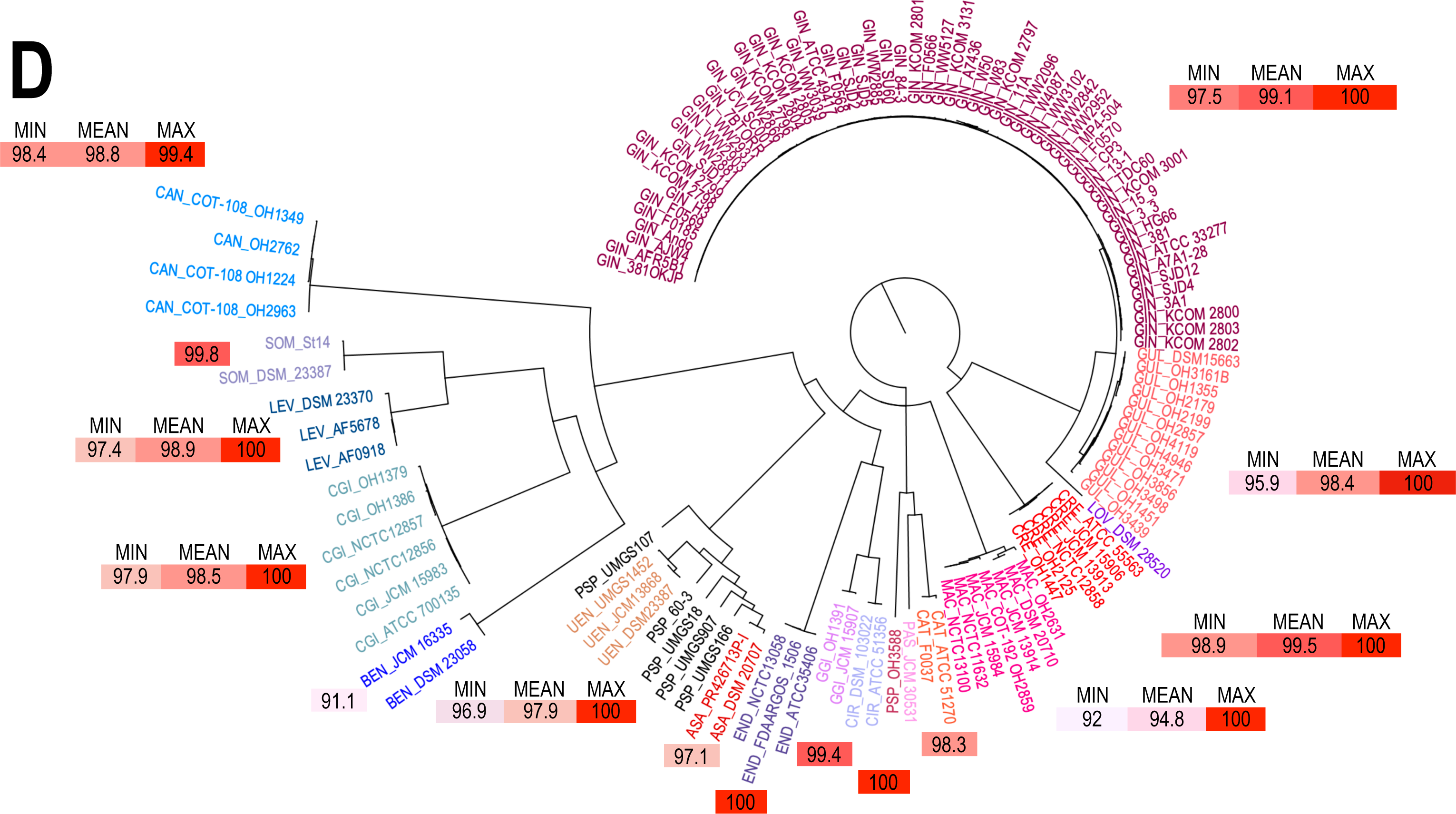
A. Violin plots of Ffp1_A and Ffp1_B amino acid lengths. From left to right: sizes as initially annotated in Genbank files (no curation), sizes after signal peptide (SP) biocuration prior to cleavage, and sizes after SP cleavage by signal peptidase II (SPII). In the box plot associated with each violin plot, the middle line represents the median and the whiskers indicate the interquartile range. **B. Multiple sequence alignment and sequence logo of Ffp1 lipobox**. Boxes represent groups of identical sequences. **C. Heat map illustrating the percent nucleotide identity** of Ffp1 signal peptides. **D. Circular phylogram of *Porphyromonas* Ffp1 proteins distance tree.** The Ffp1_A proteins are depicted in warm colors, while Ffp1_B are shown in various shades of blue. The boxes indicate the minimum, average, and maximum intraspecific identity values. If only one value is displayed, it represents the average identity percentage.

The observed differences for *P*. *asaccharolytica* are due to the presence of 33 additional nucleotides in strain PR426713P-I (at position 88-120), absent in strain DSM 20707. For *P*. *circumdentaria*, it is a 175 nt shorter annotation in strain DSM 103022 (compared to strain ATCC 51356). For *P*. *macacae* these are due to the gene encoding Ffp1_A being at the end of the contig and truncated at the 5’ end, in strain *P*. *macacae* JCM 15984. For *P*. *uenonis*, it is also the choice of an alternative start codon for the UMGS1452 strain, 34 amino acids upstream of those chosen for the DSM23387 and JCM13868 strains. Finally, for *P*. *bennonis*, at position 1410 in the DSM 23058 strain, a C base, absent from the JCM 16335 strain, leads to a frameshift. This frameshift leads to a shorter C-terminal sequence compared to DSM 23058 strain. Note that for *P*. *somerae*, the sizes are similar but the annotated sequences are “shifted” and proteins different on the N-terminal (20 aa longer in DSM_23387 compared to St14) and C-terminal (21 aa shorter in DSM_23387 due to a partial CDS at the end of the contig).

Accurate annotation of the N-terminus of proteins, which predicts their cellular localization, is crucial and deserves the attention of annotators. For this purpose, we re-annotated the start codons of Ffp1, when needed, to optimize both the SPII cleavage prediction score and the presence of charged residues at the N-terminus, followed by hydrophobic amino acids. The resulting re-annotations and their implications for cell localization predictions are listed in Table S2.

In the absence of comprehensive biocuration, a substantial part of Ffp1 proteins are predicted as cytoplasmic (*P*. *asaccharolytica*, *P*. *catoniae*, *P*. *circumdentaria* DSM 103022, *P*. *somerae* St14) or having localization predictions classified as indeterminate (PSP UMGS107, PSP UMGS166, PSP UMGS907, *P*. *uenonis* DSM 23387, *P*. *uenonis* JCM 13868). Some proteins are predicted to be cleaved by SPII, but biocuration enhances both the signal peptide prediction score and the likelihood of cleavage by SPII. As a result of this reannotation work, all Ffp1 proteins are predicted as lipoproteins, with a signal peptide of about 20 amino acids (15 to 25 aa), consistent with the requirements cited previously: 2 to 4 positively charged amino acids followed by a hydrophobic region of 10 to 15 aa (Figure S3) and a lipobox [ASG]↓C positions -1 to 1 (Figure 4B). *In silico* predictions also confirm the predicted palmitoylation (addition of acyl chains) of the cysteine residue.

These biocurated peptide signals exhibit a high degree of intra-species conservation, while demonstrating significant inter-species variability, with only a 25% pairwise identity when considering all species collectively (min. 5% - max. 100%, Figure 4C). However, two groups characterized by similar signal peptide sequences can be discerned: a first one formed by *P*. *gingivalis* and *P*. *gulae* (ca. 86% identity) and a second more consistent, composed of species *P*. *asaccharolytica*, *P*. *uenonis*, PSP_60-3, PSP_UMGS907, PSP_UMGS18 and PSP_UMGS166 (66.7 to 100% identity, Figure 4C). The same groups were observed when examining the lipobox motif.

As shown in Figure 4A (second panel), Ffp1 signal peptides biocuration not only results in more consistent predictions of their cellular localization, but also leads to a homogenization of their size, both within and across species, with the exception of *P*. *bennonis* (since the frameshift occurs in the 3’ region of the gene). This size homogenization becomes even more pronounced following signal peptide cleavage (Figure 4A, third frame). It can be seen that mature Ffp1s in group B are larger than those in group A by about 20 aa.

As shown in Figure 4D, the average intra-specific identity of the Ffp1_A subclass is very high and ranges from 100% to 94.8% depending on the species. The most divergent species are *P*. *macacae*, *P*. *gulae*, and *P*. *uenonis*. In the first two cases, this divergence can be attributed to the coexistence of two distinct homology groups within the same species. However, regrettably, the available metadata does not provide sufficient information to elucidate the underlying reasons for these discrepancies. For *P*. *uenonis*, the strain UMGS1452 that derived from a metagenome is different from the two other strains. As previously noted, the conservation of interspecific Ffp1_A sequences is small (57.5%) with only 4.5% of identical sites between all of them. When examining the Ffp1_B group, it is worth noting that the average intra-specific identity is elevated, oscillating between 98.8 and 91.1% (Figure 4D). *P*. *bennonis* is the most divergent because the two strains have proteins with the last 75 aa that differ. It is noteworthy that Ffp1_B is less homogeneous than Ffp1_A with only an average inter-specific identity of 36.4% and 4.3% identical sites. The number of conserved sites decreases to 0.7% if we compare both groups, Ffp1_A and Ffp1_B.

### 4. 3D structures confirm that *Porphyromonas Ffp1* are fimbrillins

As the signal peptide is absent in the mature protein, it was excised prior to structure prediction for all Ffp1 proteins. PSIPred predict 30% to 44% residues as strand (mean = 36.6, SD = 3.4) and 2% to 7% residues as helix (mean = 5.4, SD = 1.4) for Ffp1_A group. For group Ffp1_B, predictions concern 24% to 42% amino acids in strand (mean = 29.6, SD = 6.5) and 4% to 9% in helix (mean = 7, SD = 2.1).

The optimal structures for all *Porphyromonas* Ffp1 representatives, as predicted by Robetta and assessed by ERRAT and Verify3D, are depicted Figure 5. These structures were subjected to comparison with existing models and, the best hit, obtained either via VAST+ or Phyre2, correspond to *Bacteroides ovatus* cell adhesion protein (BACOVA_01548, 4JRF.pdb) for all *Porphyromonas* Ffp1 proteins, irrespective of species or Ffp1_class.

**Figure 5.**
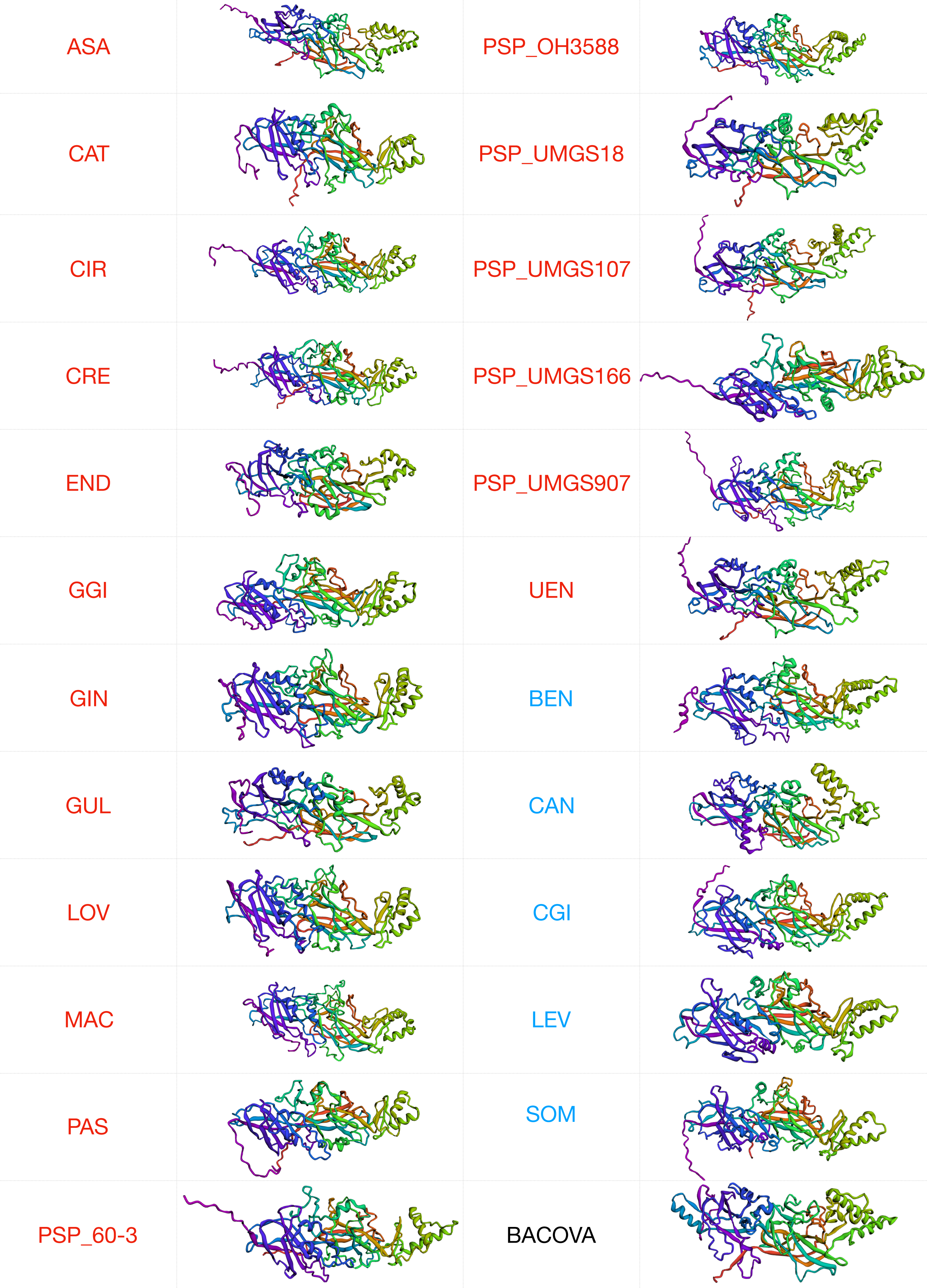
Predicted tertiary structure for mature proteins of *Porphyromonas* reference strains (one per genus). These structures correspond to predictions made by Robetta and evaluated by ERRAT and Verify3D. Only the best prediction is represented. Ffp1_A proteins are in red, Ffp1_B in blue. BACOVA_01548 was also predicted using Robetta.

According to Phyre2 and iBPA results (Figure S4), more than 82% of Ffp1 sequences were modeled with 100.0% confidence against 4JRF.pdb. Superposition of *Porphyromonas* Ffp1 and BACOVA_01548 3D structures were performed by iBPA and all evaluation values (RMSD, GDT_TS), reflect good overall similarity. For all overlapping morphologies, the aligned fraction is about 50% of the protein sequence, with mean reported RMSDs of 2.26 Å [range 2.09 to 2.53 Å] and mean GDT-TS distance scores of 32 [range 32 -37.3] (Figure S4). For *Porphyromonas gingivalis*, the structures of FimA (4Q98.pdb) and Mfa1(5NF2.pdb) are available and comparisons by superposition between Ffp1 and these two other fimbrillins (Figure S5) confirm that Ffp1 is indeed a new distinct *Porphyromonas* fimbrillin family.

### 5. *Porphyromonas* Ffp1 are ancestral orthologs but not synthelogs

In *P. gingivalis*, *ffp1* is the fourth gene in an operon-like structure comprising a gene encoding a cysteinyl-tRNA synthetase, a second gene encoding a patatin-like protein, and a third gene encoding a group 2 glycosyltransferase. An identical *locus* is found in all *P. gulae* genomes, while is absent in any other *Porphyromonas* (Figure S6). The *P. asaccharolytica*, *P. uenonis*, *P*. sp. UMGS18 and *P*. sp. UMGS107 group, mentioned earlier in this article, show a syntenic pattern upstream of *ffp1*, characterized by the presence of two conserved genes encoding dihydroorotate dehydrogenases, crucial enzymes involved in *de novo* pyrimidine biosynthesis in prokaryotic cells. However, there is no direct association with the fimbrillin function. Furthermore, the intergenic space spanning approximately 300 nucleotides is sufficiently substantial to preclude any functional linkage between these genes. The second group, comprising *P*. sp. UMGS166 and *P*. sp. UMGS907, also previously mentioned, shows synteny downstream of *ffp1* with a gene encoding a nitronate monooxygenase (degradation of propionate-3-nitronate) and another encoding a 4-hydroxy-tetrahydrodipicolinate synthase (involved in lysine biosynthesis). Similar to the previous group, no discernible functional relationship appears to exist, and the intergenic space of about 200 nt seems to confirm this hypothesis. For the other *Porphyromonas*, each species exhibits a distinct gene organization arrangement surrounding *ffp1* (Figure S6).

In conclusion, with the exception of phylogenetically closely related species, we find no preserved synteny in the *ffp1* locus, which would reflect the absence of co-localization constraints for co-functional genes. Nevertheless, as demonstrated by the tanglegram juxtaposing the orfeome tree and the Ffp1 tree (Figure 6), the remarkable congruence between these two trees provides compelling evidence that Ffp1 is an ancestral protein of *Porphyromonas,* and its evolution would have closely paralleled the evolutionary trajectory of the entire genus. This observation also holds true for the differentiation between the two Ffp1 classes (Figure 6). The absence of genes conservation in close chromosomal proximity to *ffp1,* along with the presence of a significant 5’ intergenic space (Figure S6), not only signifies the absence of selection pressure around this gene but also strongly suggests that *ffp1* functions as an independent transcriptional unit.

**Figure 6.**
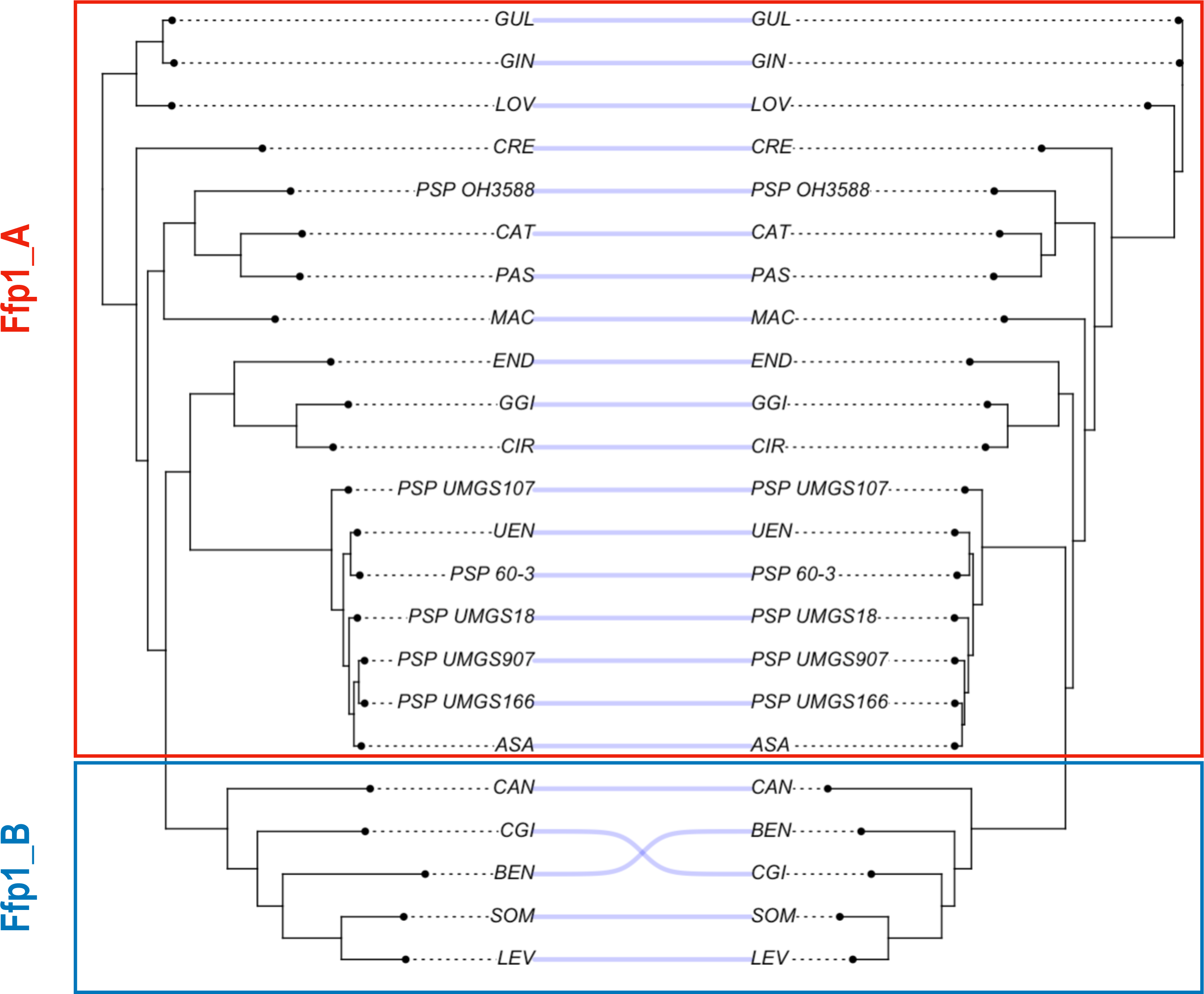
Tanglegram comparing the tree constructed from the primary sequences of the Ffp1 proteins in representative strains of *Porphyromonas* (on the left) with the species tree based on the orthology of the orfeomes.

## Discussion

Our analysis of fimbrillin *loci* within the *Porphyromonas* genus initiated with the retrieval of genomes from the NCBI RefSeq database. The first step encompassed the validation of genus-level assignment for each genome retrieved, followed, when feasible, by species-level confirmation. The Overall Genome Relatedness Indices (OGRI), namely: digital DNA-DNA hybridization distance (DDH) and genome Average Nucleotide Identity (gANI) were used to classify genomes into monophyletic groups. These OGRIs are increasingly used in taxogenomic studies and serve as a valuable tool for validating the taxonomic classification of isolates of interest (59). Likewise, in accordance with prior research, we employed more conventional methodologies for species-level genomes grouping, such as evaluating the percentage identity of the gene encoding 16S rRNA (when annotated) (60).Our study underscores the critical necessity of rigorously confirming the taxonomic classifications of genomes before embarking on any comparative genomics analysis to ensure their accuracy. Moreover, this checking step enables the possibility of taxonomic reassignment when warranted, aligning with findings from previous studies (61–63). In this investigation, we have identified genomes erroneously labeled as *Porphyromonas* (i.e strain 31_2, which is a *Parabacteroides*), misassignment of *Porphyromonas* to species (i.e strain 60.3, which does not belong to *P. uenonis*) as well as metagenomic mixture such as strain KA000683 imperfectly assigned to *P. somerae*.

Our study also raises questions about genomes assigned to *Porphyromonas* without any species assignment (28 out of 144 genomes, i.e., 19.5%). They all correspond to incomplete draft genomes which introduces bias into studies that rely on them (64). We can specifically mention the presence of gaps, local assembly errors, chimeras and contamination by fragments from other genomes (65,66). This contamination, defined as the presence of foreign sequences within a genome, can lead to incorrect functional inferences such as higher rates of horizontal gene transfer (HGT) and errors in phylogenomic studies. Such errors can be propagated throughout the scientific community and have been documented to exist in databases (67). To mitigate these types of errors, several studies, including the present one, advocate the practice of data biocuration throughout the study. To identify potential contamination in draft genomes, we employed Kraken2 software and assessed the cumulative contig size of incomplete genomes. By applying specific inclusion criteria, we were able to disqualify 17 draft genomes, corresponding to metagenomic mixtures and inaccurately labeled as *Porphyromonas*. Furthermore, among the 11 remaining draft genomes, our taxogenomic approach led to the reclassification of 5 genomes into four previously described species (*P. gulae*, *P. asaccharolytica*, *P. uenonis* and 2 genomes in *P. canoris*). The remaining 6 genomes that cannot be assigned to already described species may potentially represent novel, yet undescribed species, akin to hypotheses proposed in other bacterial genera (63,67). This suggests that the genus *Porphyromonas* may encompass a greater degree of species diversity than previously recognized.

Thus, in this study, we retained 126 *Porphyromonas* genomes (24 clades comprising 17 species and 7 singletons) to describe fimbriae loci. To accomplish our research objectives, distant homology between proteins must be detected and is fundamental for enabling comparative and evolutionary investigations, shedding light on protein families, and providing insights into their molecular structures and functions (68).

Current orthology detection methods include Position Specific Scoring Matrix (PSSM) techniques, like PSI-BLAST (Position-specific iterated BLAST, (69), which generate substitution score profiles by accounting for residue variability within homologous sequence families (70). An even more effective approach involves Hidden Markov Model (HMM) profiles, which incorporate emission and transition state probabilities at each protein sequence position, making them a superior choice for identifying distant homology (70,71).

Using ontology as a protein search strategy search is ineffective, as most fimbrillin genes are poorly annotated or annotated as “hypothetical protein” (between 21.1% and 88.6% of annotated genes). Specifically, stem and anchor proteins (FimAB or Mfa12) are better annotated with deficient annotation rates ranging from 21.1% to 50.7%. In contrast, accessory proteins (FimCDE or Mfa345) suffer from particularly poor annotations with error percentages ranging from 58.9% to 88.6%. These annotation errors are present within the databases and, without biocuration and correction, are likely to persist, potentially perpetuating inconsistencies, inaccuracies, and errors in subsequent genome annotations (72). For example, for a gene family, nearly 20% of sequences may exhibit significant errors such as inaccuracies in gene names, partial sequences or initiation codon misassignments (73). In the context of less extensively researched bacterial species, as is the case in this study, the prevalence of erroneous or uninformative annotations are much higher, reaching 77.1% of sequences identified as Ffp1 where the annotation was “hypothetical protein” or “lipoprotein”.

In this study, we leveraged 12 HMM profiles developed from *P. gingivalis* genomes, which were further refined through a strategy involving functional domain screening, clustering and biocuration. This approach enabled a comprehensive exploration of the *Porphyromonas* orfeomes, revealing variations in the three fimbriae loci across all species within this genus.

The *fimABCDE* locus is present in 9 (of 24 groups, or 37.5% of *Porphyromonas* species) with two distinct *fim loci* present in all *P. macacae* genomes. The *mfa12345 locus* is present only in three closely related species (*P. gingivalis*, *P. gulae* and *P. loveana*). For this *locus*, hybrid *fim/mfa* or *ffp1/mfa loci* are present in two species (*P. endodontalis* and *P. macacae*): *mfa123_fimDE* and *ffp1-like_mfa2345* in *P. macacae*; and a distinctive six-gene *locus* in *P. endodontalis*. This *locus* encompasses genes encoding Mfa1-like, Mfa2, and Mfa3-like proteins, along with two genes responsible for lipoproteins and a gene encoding a protein featuring a von Willebrand factor type A (VWA) domain. Interestingly, for the gene encoding Mfa5, the prevailing description is rather nondescript, simply stating it as a “protein containing a VWA domain”. This description, however, falls short in conveying the functional significance of this gene. It’s worth emphasizing that proteins featuring VWA domains play pivotal roles in diverse biological processes, including but not limited to cell adhesion and defense mechanisms. Thus, a more detailed annotation is warranted to better appreciate the functional implications of Mfa5 (74).

Finally, other species (i.e. *P. asaccharolytica*, *P. circumdentaria*, *P. crevioricanis*, *P. gingivicanis* and *P. uenonis*) have fimbrilin genes identified through HMM profiles that remain uncharacterized. These two *loci*, *fimABDCE* and *mfa12345*, have been described in other closely related species, for example an Mfa system (with only *mfa1* and *mfa2*) in *Bacteroides thetaiotaomicron* (75), and a cluster with *fimABCDE*-like genes and genes similar to either *mfa1*/*mfa2* or *mfa4*/*mfa2* with either *mfa1* or *mfa4* encoding the fimbriae stem and *mfa2* as an anchor in *Parabacteroides distasonis* (76). The *fim* and *mfa loci* in *Porphyromonas* spp. will be the main subject of an ulterior publication.

Concerning Ffp1 fimbriae (77.1% of all *ffp1* genes were deficiently annotated), this protein was most recently described in *P. gingivalis* (13,14). The encoding gene has two variants, denoted as A and B in our study. Ffp1_A is the predominant variant found in 19 *Porphyromonas* species/groups, whereas Ffp1_B is restricted to only 5 species (*P. bennonis*, *P. canoris*, *P. cangingivalis*, *P. levii*, and *P. somerae*). Furthermore, this study demonstrates that the utilization of HMM profiles reveals that *ffp1* is confined to the *Porphyromonas* genus and is absent in closely related genera like *Bacteroides* or *Prevotella*. This finding contrasts with approaches employing BLASTp and psiBLAST (16).

The presence of multiple fimbriae loci within genomes is a common phenomenon observed in other bacterial models. These loci are often associated with general niche colonization abilities or the adhesion to more specific substrates (77,78). Further investigations are needed on species more closely related to *Porphyromonas* and within this bacterial genus. These studies can shed light on aspects such as host specificity and their association with species-related pathologies (79).

Given that the majority of *in silico* coding sequence (CDS) annotators tend to prioritize the prediction of the longest possible Open Reading Frame (ORF) by favoring the initiation codon (ATG) over alternative codons (TTG and GTG) (80,81), and considering the variable size of proteins across *Porphyromonas* species, we conducted a thorough examination of the annotated initiation codons for each predicted Ffp1. Given that fimbrillins are lipoproteins (9), their N-terminal region is expected to feature a signal peptide starting with positively charged amino acids, followed by hydrophobic amino acids, and concluding with a cysteine-terminated lipobox, which serves as the cleavage site for signal peptidase II. The biocuration of start codons led to a more consistent protein size post-signal peptide cleavage. Additionally, the extracellular prediction of mature lipoproteins was confirmed, characterized by the presence of charged and hydrophobic residues, the lipobox, and a palmitoylation site. These features align with the ancestral nature of FFp1.

In addition, Ffp1 3D modeling of the mature protein was performed with several software packages, and the predictions were evaluated with classical metrics (82,83). In all cases, the generated models were compared with existing 3D structures, and the most significant match was found with the cell adhesion protein BACOVA_01548 from *Bacteroides ovatus* (3). This *B. ovatus* protein has not been extensively studied, but was classified by the authors as the stem of a type V pilus, sharing common features with type V fimbriae. These characteristics include export to the periplasm as a lipoprotein (prepilin), subsequent delivery to the outer membrane, translocation to the cell surface and cleavage by Rgp (Arg-gingipain) (4,84).

Moreover, this new fimbrillin, Ffp1 exhibits notable distinctions from both FimA and Mfa1, as evident from the obtained metrics when superimposing the 3D structures of these proteins available for *P. gingivalis*. Furthermore, the gene arrangement of *ffp1* is differs from the *fim* and *mfa* operons as the gene encoding Ffp1 does not appear to be in an operon structure.

## Concluding remarks

HMM profiles are potent tools for detecting distant homologies and facilitating phylogenetic studies. For conducing these investigations, meticulous manual biocuration is essential, as with any *in silico* research. In this article, these HMM profiles make it possible to discriminate, without ambiguity, three *Porphyromonas* fimbriae and to describe their distribution: *mfa12345*, limited to the three closely related species (*P. gingivalis*, *P. gulae* and *P. loveana*), *fimABCDE* present in nearly 40% of the *Porphyromonas* species and *ffp1*, present in all *Porphyromonas* but restricted to this bacterial genus. Our study predicts that Ffp1 is a new fimbrillin, distinct from FimA and Mfa1. It is closely related to another type V fimbrillin protein, BACOVA_01548, as evidenced by manual start codon curation and 3D modeling. Given the ancestral nature of Ffp1, as elucidated by our study, and its presence in all studied *Porphyromonas* genomes, in contrast to the fimbrillins Fim and Mfa, the question of its function becomes paramount. Especially in absence of co-localization of accessory genes ensuring its stability, assembly, and anchorage to the cell surface. What role does it play in the production and cargo of Outer Membrane Vesicles (OMVs), a phenomenon observed in numerous studies? Further wet-lab investigations are necessary to address these pending inquiries.

## Supporting information

HMMprofiles

## Legend to supplementary figures

**Figure S1. Sankey diagrams representing Kraken 2 report results for each *Porphyromonas* sp. genome.** Each diagram illustrates the percentage of the genome classified under the genus *Porphyromonas* and the cumulative size of the accurately assigned fragments for each strain.

**Figure S2. A-C-E:** Box plot representing hmmsearch E-value results for all *Porphyromonas* genus groups and all HMM profiles used in this study. The dotted lines represent the thresholds used. **B-D-F:** Frequencies of the terms employed to describe the genes of the 3 fimbriae systems of *Porphyromonas*, as originally annotated in the Genbank files.

**Figure S3. Ffp1 signal peptide prediction by the SignalP-6.0.** For each *Porphyromonas* group, a reference sequence was chosen (named at the top of the SignalP6 graphs). For each group, we present the intra-specific consensus sequence of the signal peptides (after biocuration when required) in the form of a logo. The blue stars represent the charged amino acids (predicted by EMBOSS charge prediction) and the orange curve represents the prediction of hydrophobicity (Protscale, Amino acid Hydropathicity using Kyte and Doolittle method).

**Figure S4. Evaluation of 3D models of Ffp1 proteins predicted *in silico***.

Assessments were performed using iBPA and Phyre2 based on template 4JRF (BACOVA_01548). Right column shows superposition of predicted structure (green) with model structure (in red) using iPBA.

**Figure S5. Comparison of Robetta structures prediction for *P. gingivalis* FimA, Mfa1 and Ffp1.** The boxes represent the results of the superposition in Ffp1 and either BACOVA_01548, GIN_FimA or GIN_Mfa1.

**Figure S6. Diagram of synteny in the vicinity of the *Ffp1* gene in the different species of *Porphyromonas*.** Ffp1 genes are shown in light blue and surrounding genes in dark blue using Geneious Prime. The yellow boxes correspond to contig extremities.

## Supplementary tables

**Table S1. List of *Porphyromonas* genomes used in this study.**

Genomes were grouped into clades following genomic data-driven taxonomic clustering. Complete genomes are in green, *Porphyromonas* sp. grouped in a clade are in blue, a mislabelled *P. somerae* is indicated in yellow, *P.* sp. genomes that could not be grouped with others are in dark grey and two mislabelled “*Porphyromonas*” genomes are in purple. All accession numbers are indicated as well as the strain.

**Table S2. List and details of all *ffp1* genes identified during this study.**

Information includes HMM group membership, old and new locus_tags, items related to start codon reannotation (when needed), cell localization predictions, peptide signal, and before and after cleavage sizes.

## Supplementary material

**HMM profiles.** This archive contains the 12 HMM profiles that were generated and employed in this study for the detection and the classification of *Porphyromonas* fimbriae.

## Footnotes

* https://github.com/kblin/ncbi-genome-download

† https://github.com/andrewfrank/ggdc-robot

‡ https://www.ezbiocloud.net/tools/orthoani

§ http://fbac.dmicrobe.cn/tools/

** http://tree.bio.ed.ac.uk/software/figtree/

†† https://github.com/sanderlab/alignmentviewer

‡‡ https://cran.rstudio.com/web/packages/TreeDist/index.html

§§ https://cran.r-project.org/web/packages/phytools/index.html

