## Supplementary material for "Ffp1, an ancestral *Porphyromonas* spp. fimbrillin": SupplFigures1to6 and SupplTables1to2

| Strain | % Classified | % *Porphyromonas* | Size (kb) | Conclusions | Kept in study |
|---|---|---|---|---|---|
| **31-2** | 100 | 0 | 0 | ***Parabacteroides distasonis*** | **NO** |

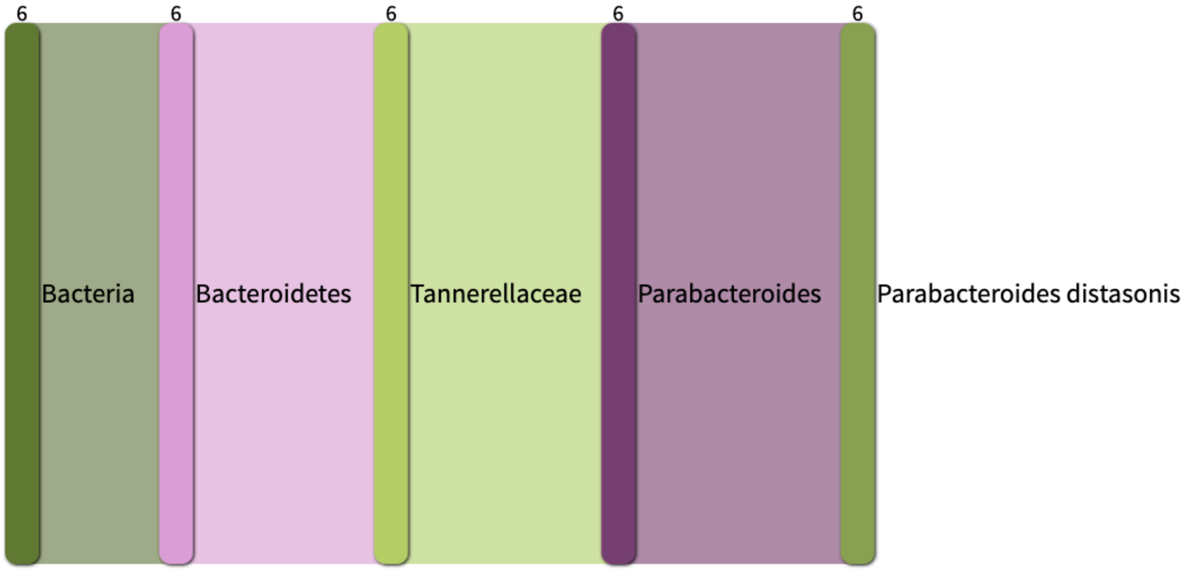

| Strain | % Classified | % *Porphyromonas* | Size (kb) | Conclusions | Kept in study |
|---|---|---|---|---|---|
| **bin_26** | 100 | 7 | 73 183 | **Genomes mixture** | **NO** |

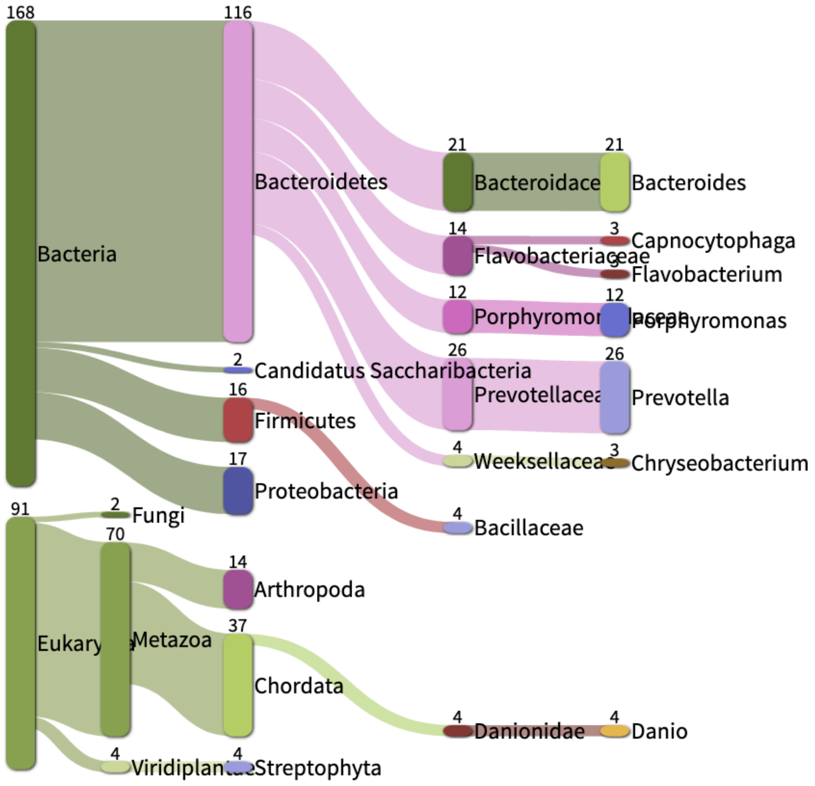

| Strain | % Classified | % *Porphyromonas* | Size (kb) | Conclusions | Kept in study |
|---|---|---|---|---|---|
| CAG_1061 | 95.64 | 17 | 326 475 | **Genomes mixture** | **NO** |

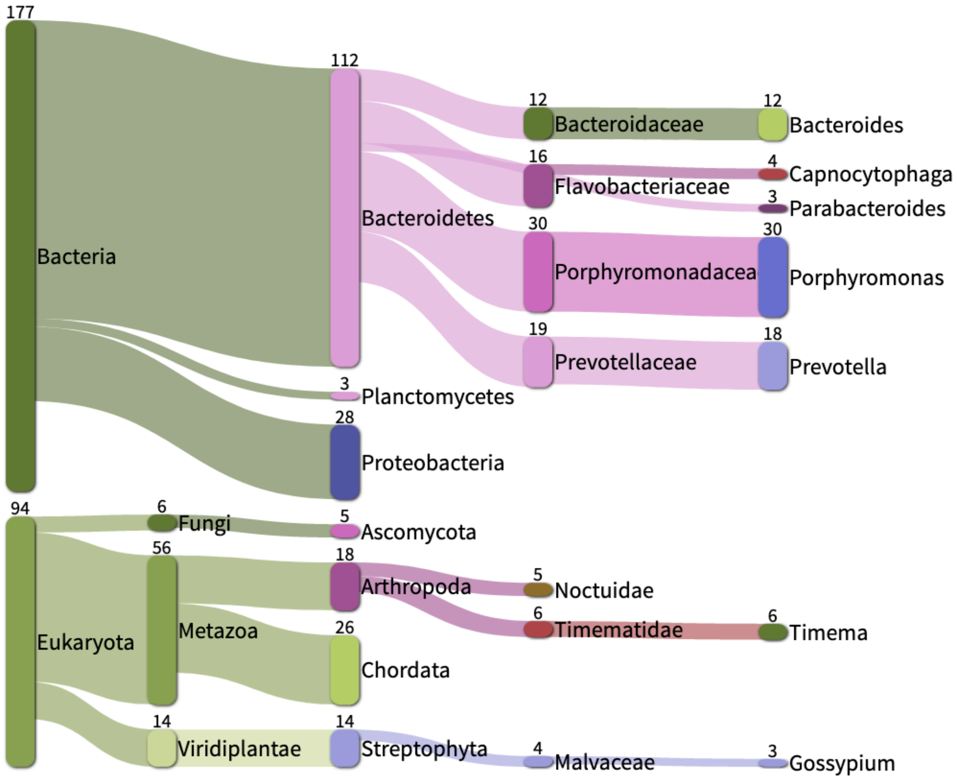

| Strain | % Classified | % *Porphyromonas* | Size (kb) | Conclusions | Kept in study |
|---|---|---|---|---|---|
| F450 | 100 | 43 | 1 363 365 | **Genomes mixture** | **NO** |

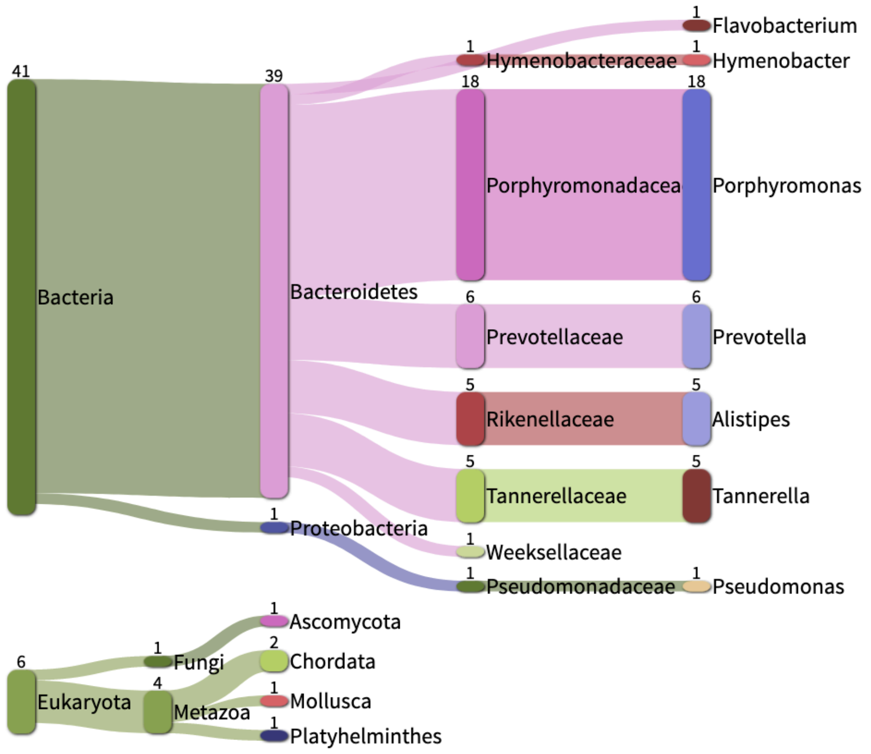

| Strain | % Classified | % *Porphyromonas* | Size (kb) | Conclusions | Kept in study |
|---|---|---|---|---|---|
| HMSC065F10 | 87.38 | 37 | 1 008 234 | **Genomes mixture** | **NO** |

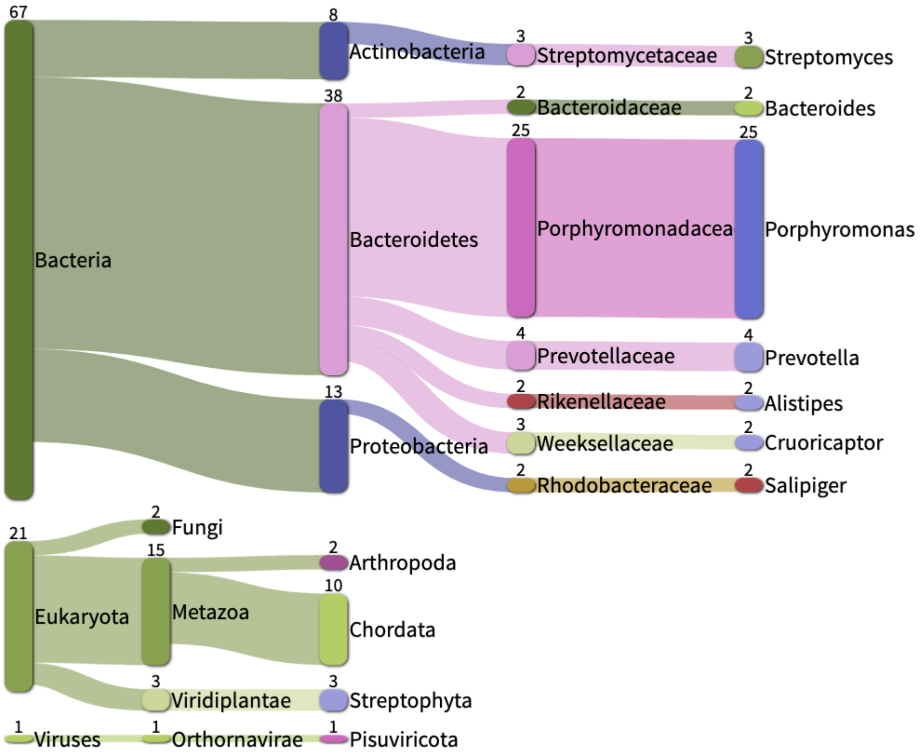

| Strain | % Classified | % *Porphyromonas* | Size (kb) | Conclusions | Kept in study |
|---|---|---|---|---|---|
| HMSC077F02 | 78.82 | 49 | 1 162 849 | **Genomes mixture** | **NO** |

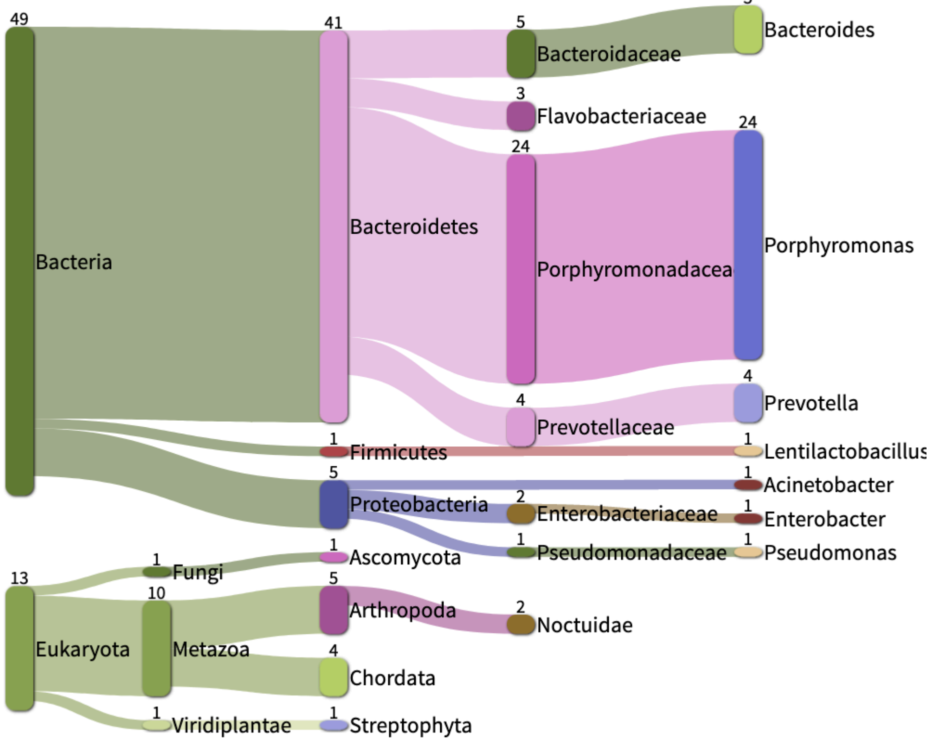

| Strain | % Classified | % *Porphyromonas* | Size (kb) | Conclusions | Kept in study |
|---|---|---|---|---|---|
| somerae KA00683 | 93.55 | 32.5 | 1 028 690 | **Genomes mixture** | **NO** |

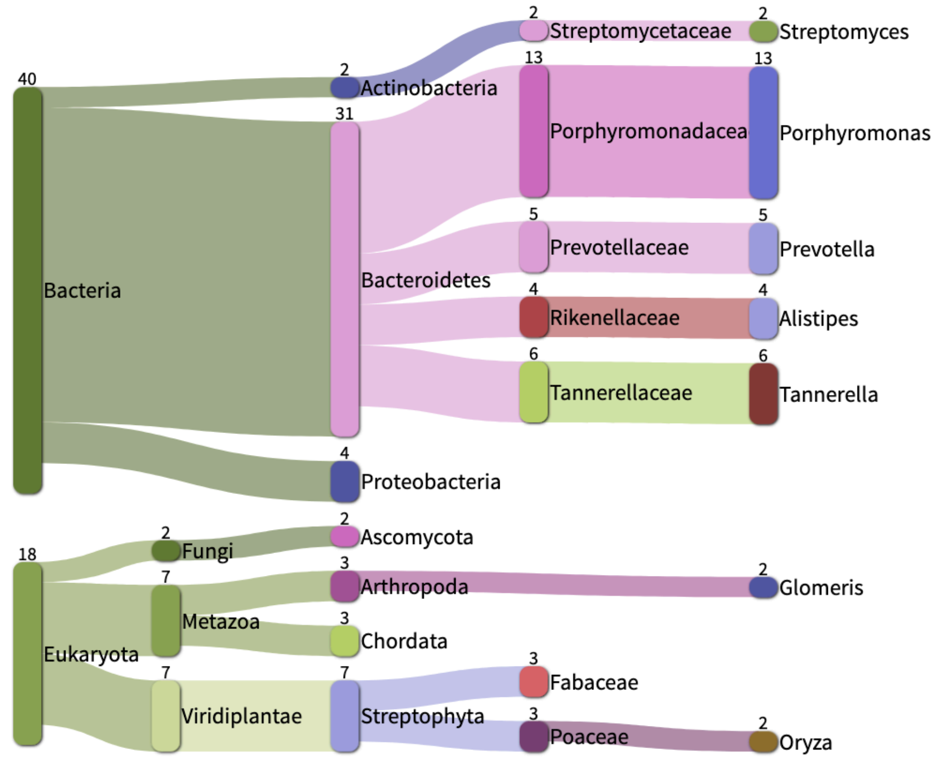

| Strain | % Classified | % *Porphyromonas* | Size (kb) | Conclusions | Kept in study |
|---|---|---|---|---|---|
| KLE_1280 | 100 | 50 | 1 726 267 | **Genomes mixture** | **NO** |

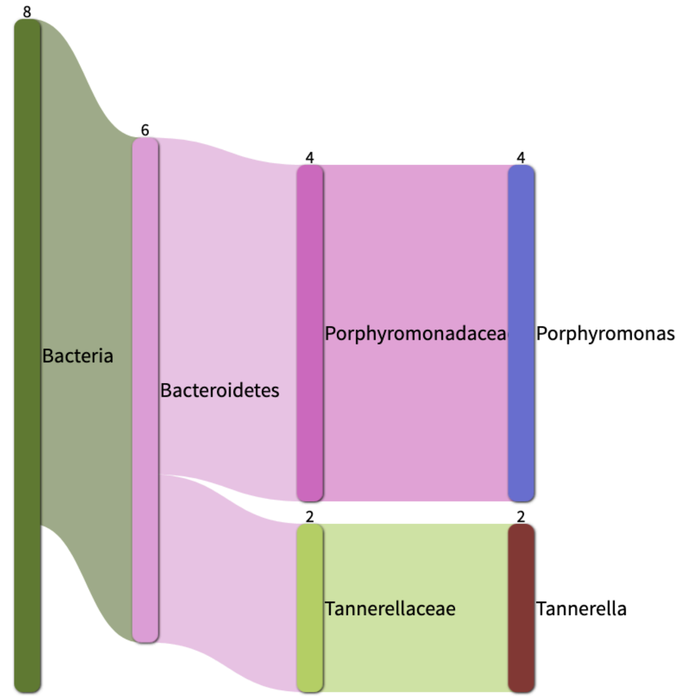

| Strain | % Classified | % *Porphyromonas* | Size (kb) | Conclusions | Kept in study |
|---|---|---|---|---|---|
| MGY-HGUT-04270 | 100 | 20 | 230 325 | **Genomes mixture** | **NO** |

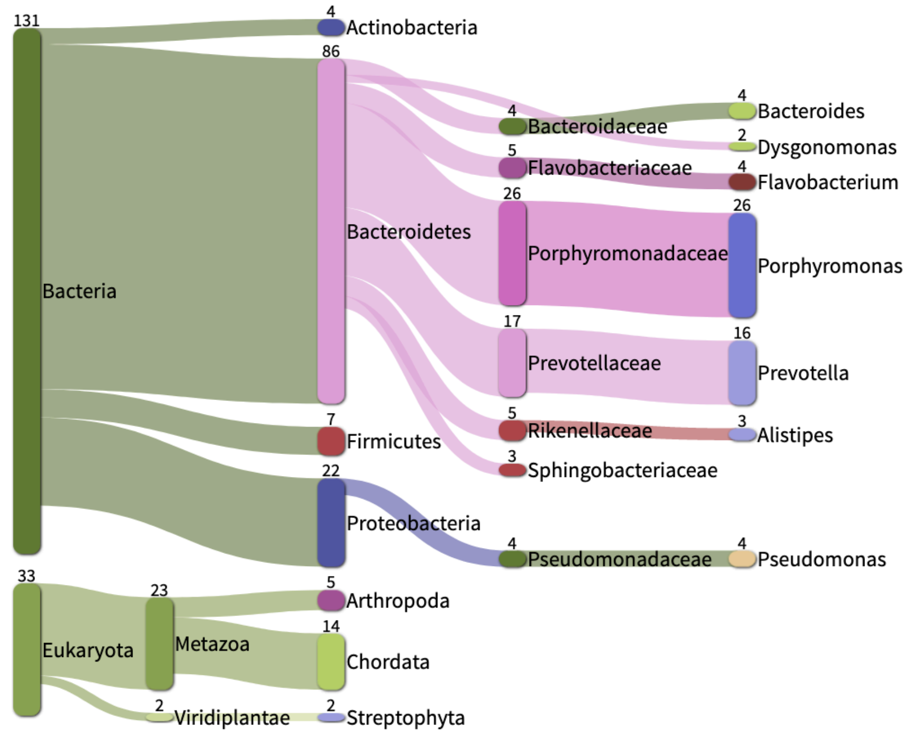

| Strain | % Classified | % *Porphyromonas* | Size (kb) | Conclusions | Kept in study |
|---|---|---|---|---|---|
| OH860 | 95.12 | 48 | 1 418 732 | **Genomes mixture** | **NO** |

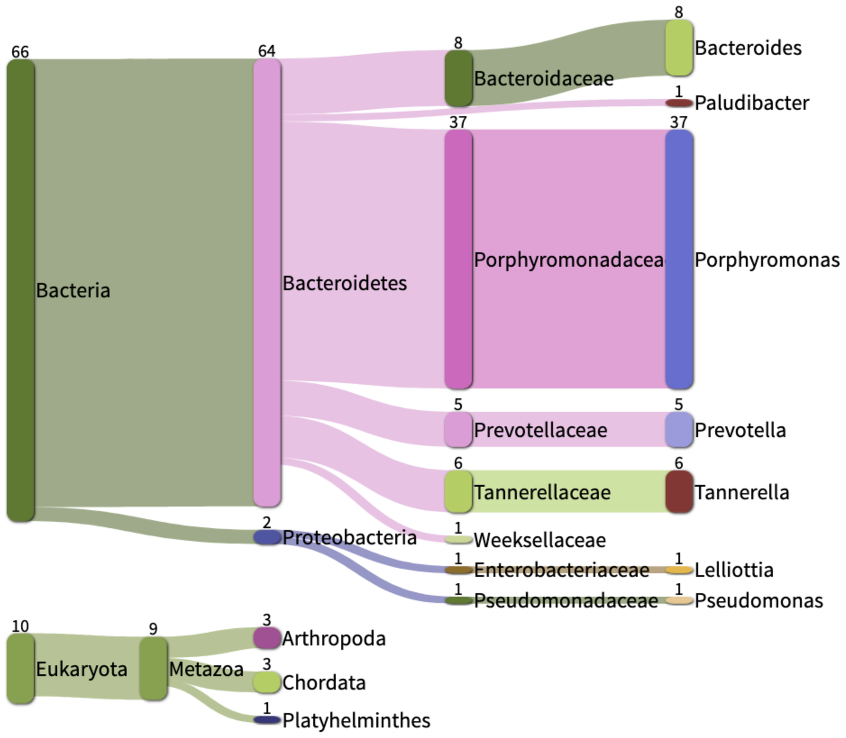

| Strain | % Classified | % *Porphyromonas* | Size (kb) | Conclusions | Kept in study |
|---|---|---|---|---|---|
| OH1446 | 94.5 | 45 | 1 393 092 | **Genomes mixture** | **NO** |

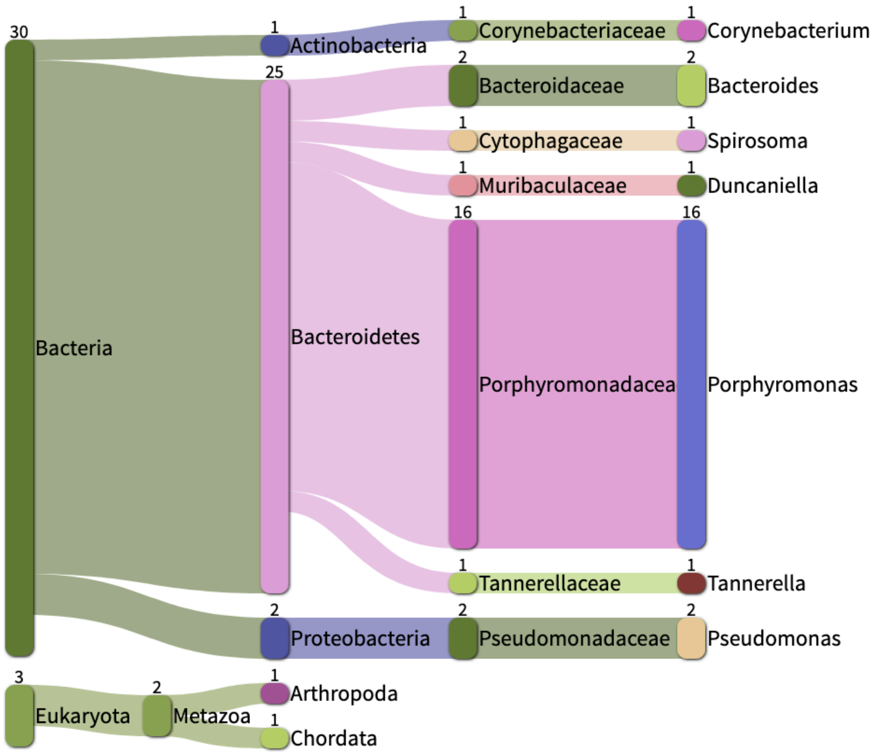

| Strain | % Classified | % *Porphyromonas* | Size (kb) | Conclusions | Kept in study |
|---|---|---|---|---|---|
| UMGS547 | 100 | 36 | 918 508 | **Genomes mixture** | **NO** |

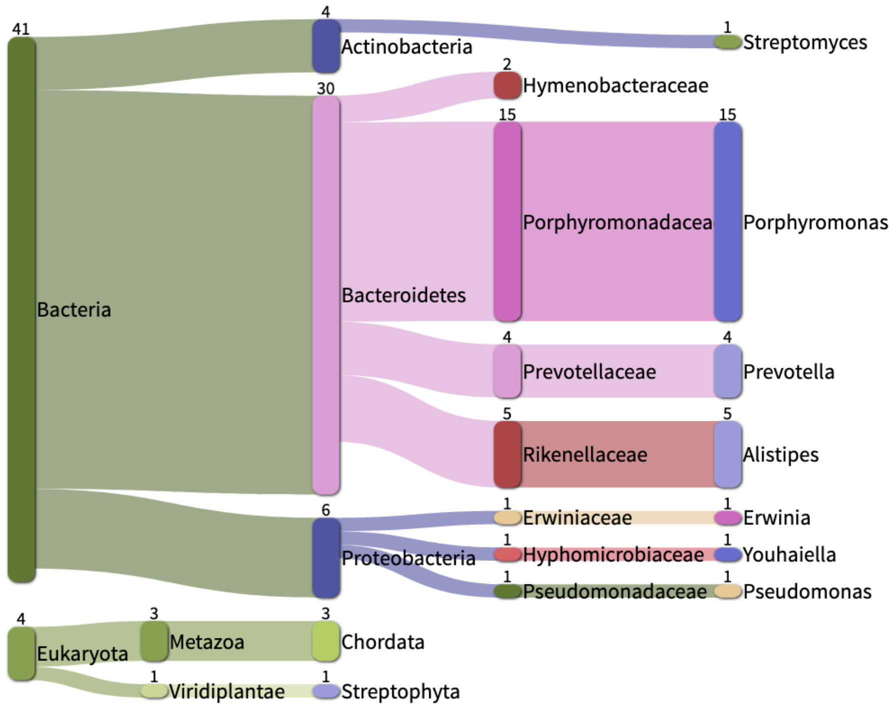

| Strain | % Classified | % *Porphyromonas* | Size (kb) | Conclusions | Kept in study |
|---|---|---|---|---|---|
| UMGS338 | 100 | 24 | 907 529 | **Genomes mixture** | **NO** |

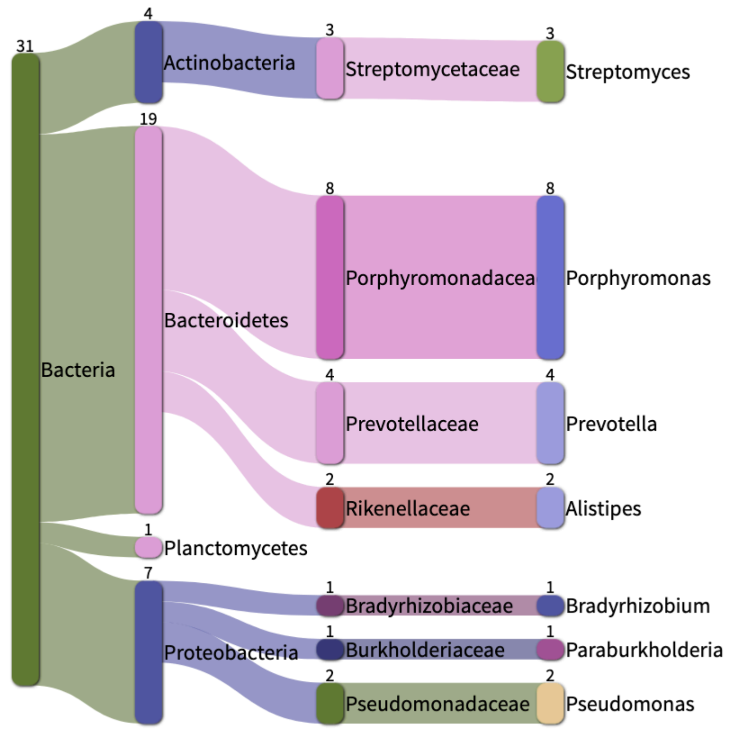

| Strain | % Classified | % *Porphyromonas* | Size (kb) | Conclusions | Kept in study |
|---|---|---|---|---|---|
| UMGS713 | 99.39 | 20 | 458 617 | **Genomes mixture** | **NO** |

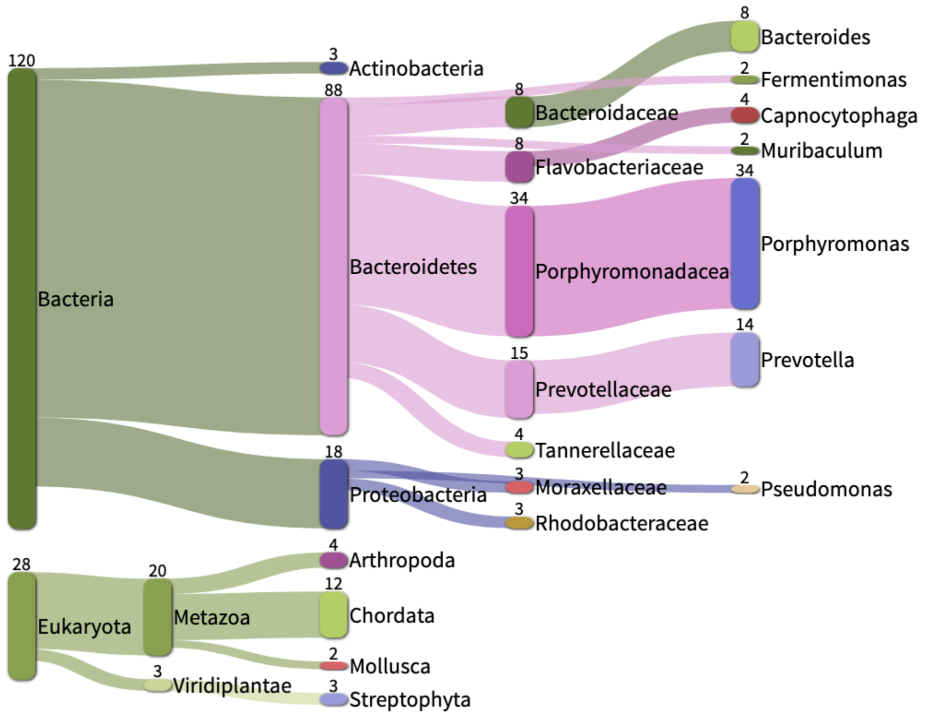

| Strain | % Classified | % *Porphyromonas* | Size (kb) | Conclusions | Kept in study |
|---|---|---|---|---|---|
| UMGS1769 | 100 | 10 | 199 747 | **Genomes mixture** | **NO** |

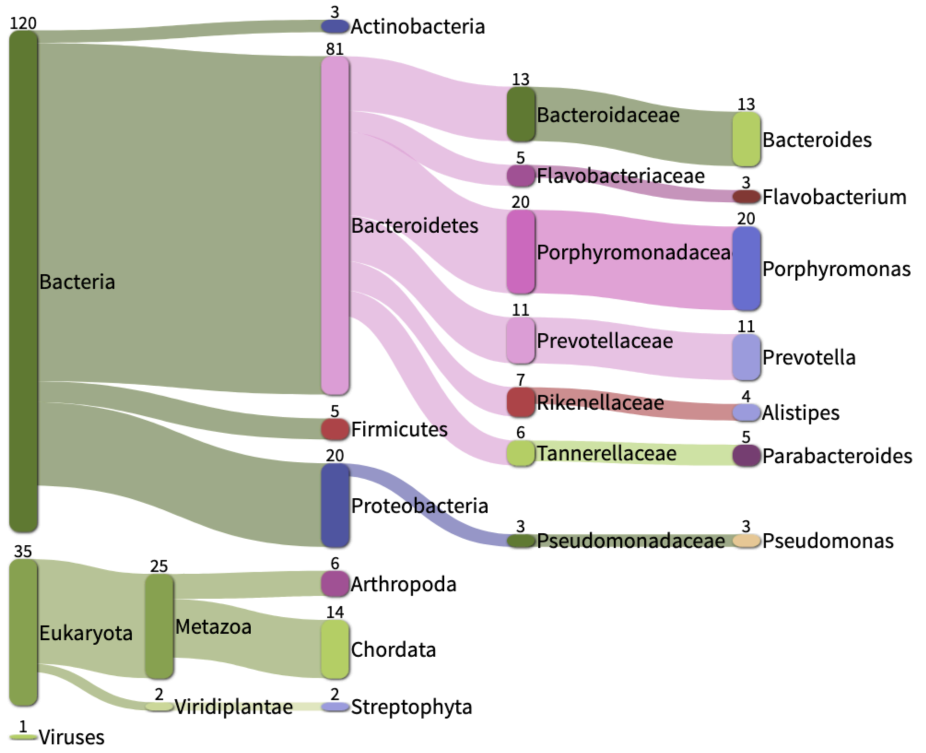

| Strain | % Classified | % *Porphyromonas* | Size (kb) | Conclusions | Kept in study |
|---|---|---|---|---|---|
| UMGS2020 | 99.64 | 4 | 46 937 | **Genomes mixture** | **NO** |

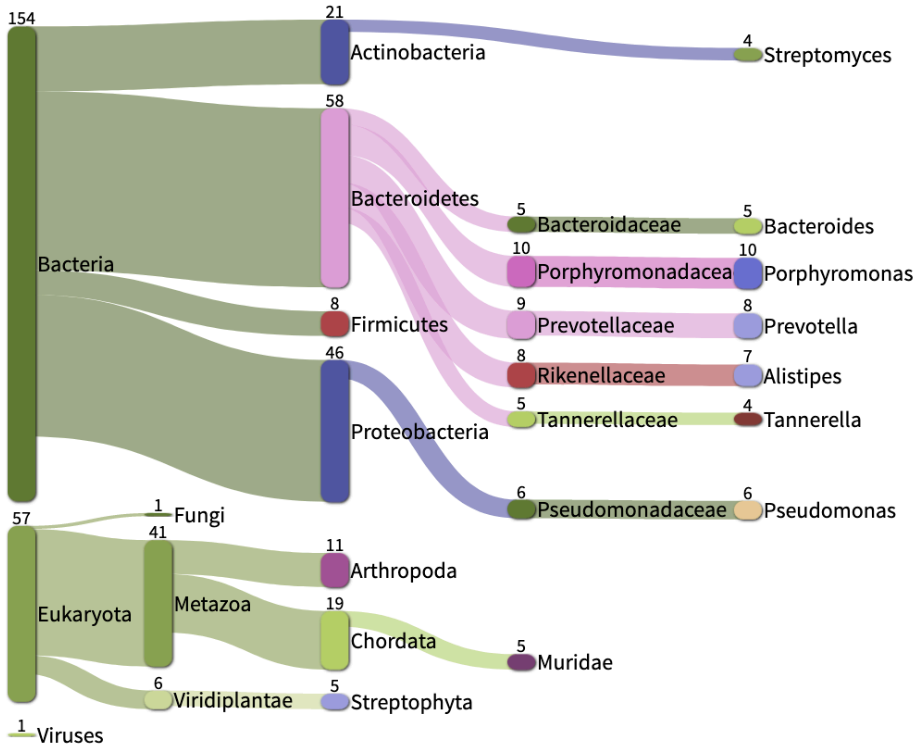

| Strain | % Classified | % *Porphyromonas* | Size (kb) | Conclusions | Kept in study |
|---|---|---|---|---|---|
| UMGS2040 | 100 | **100** | **768 256** | **Too fragmented, Too incomplete** | **NO** |

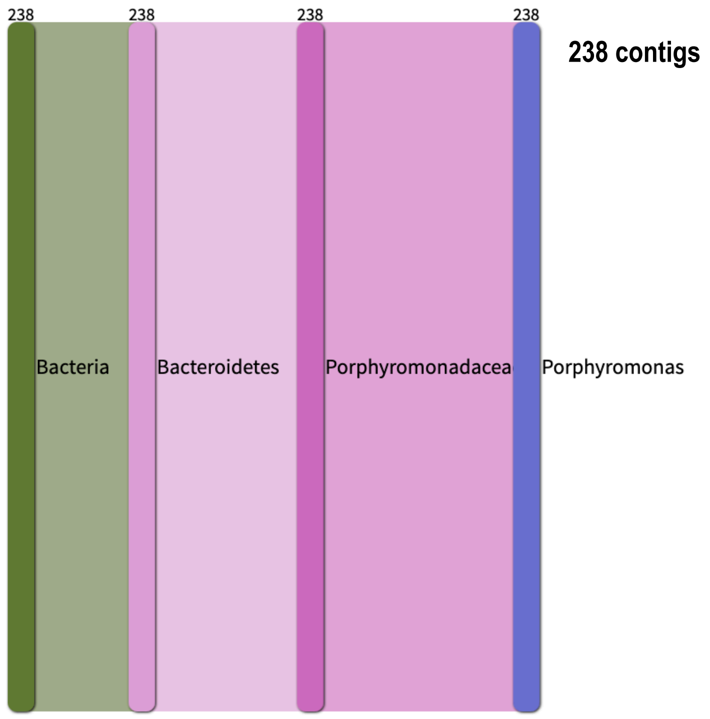

| Strain | % Classified | % *Porphyromonas* | Size (kb) | Conclusions | Kept in study |
|---|---|---|---|---|---|
| W7784 | 97.4 | 35 | 1 858 400 | **Genomes mixture** | **NO** |

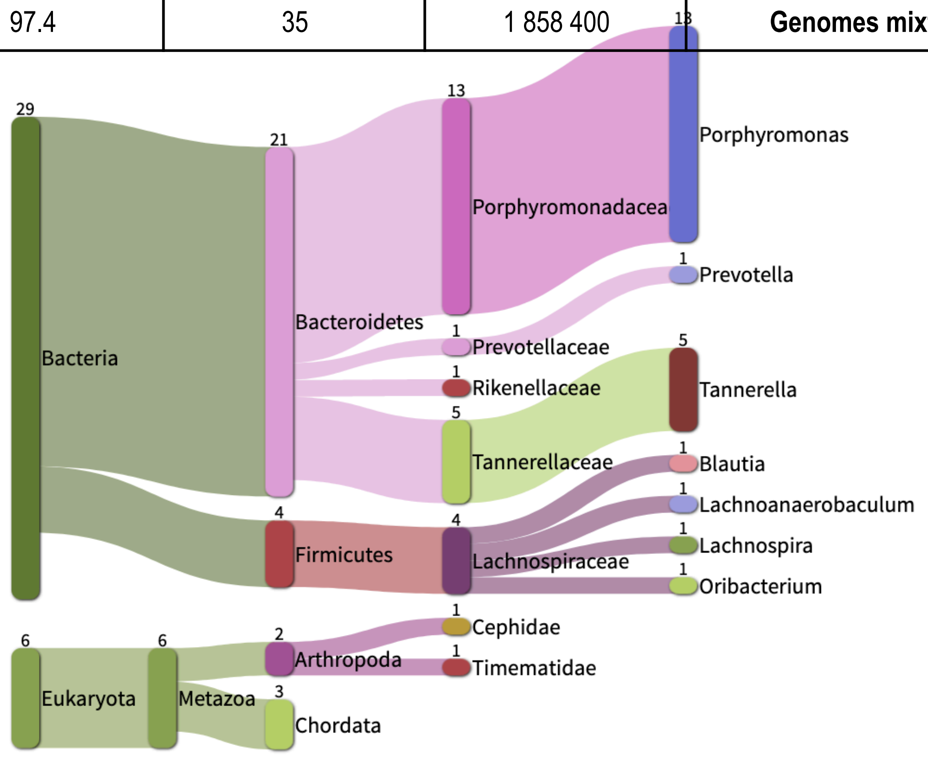

| Strain | % Classified | % *Porphyromonas* | Size (kb) | Conclusions | Kept in study |
|---|---|---|---|---|---|
| MGY-HGUT-04267 | 100 | **100** | 2 088 006 | ***P. asaccharolytica*** | **YES** |

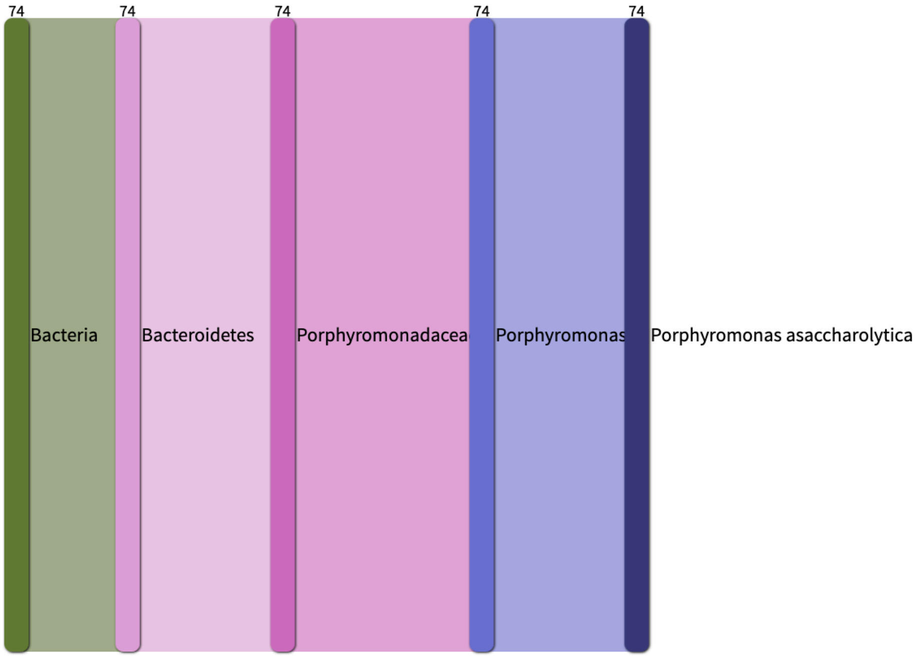

| Strain | % Classified | % *Porphyromonas* | Size (kb) | Conclusions | Kept in study |
|---|---|---|---|---|---|
| OH1349 | 93 | **80** | 2 255 484 | **Mainly *Porphyromonas*** | **YES** |

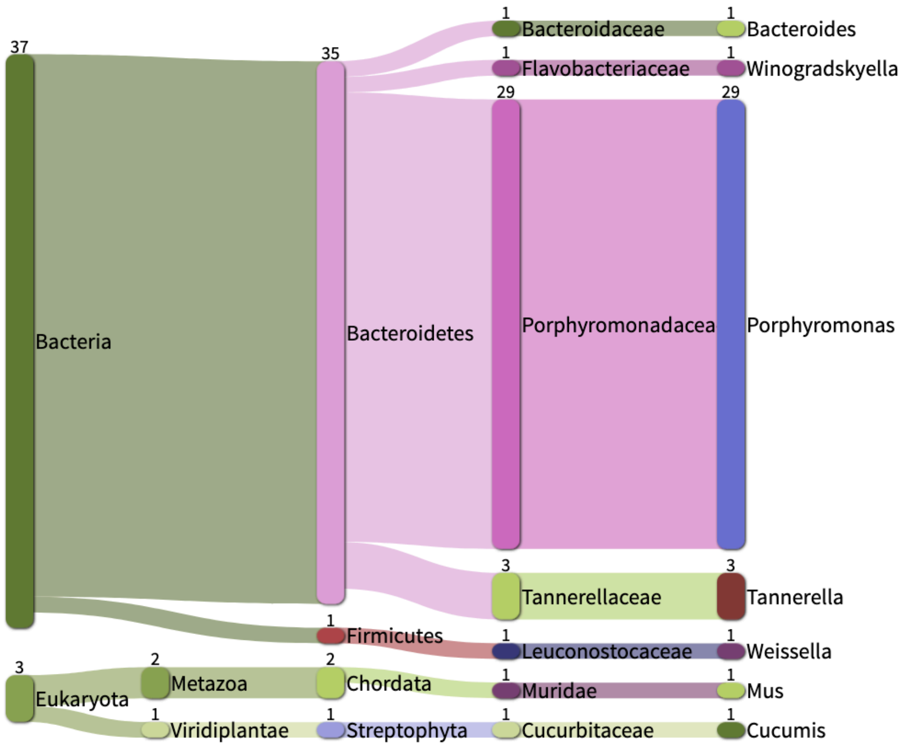

| Strain | % Classified | % *Porphyromonas* | Size (kb) | Conclusions | Kept in study |
|---|---|---|---|---|---|
| OH4946 | 97 | **88** | 2 117 249 | **Mainly *Porphyromonas*** | **YES** |

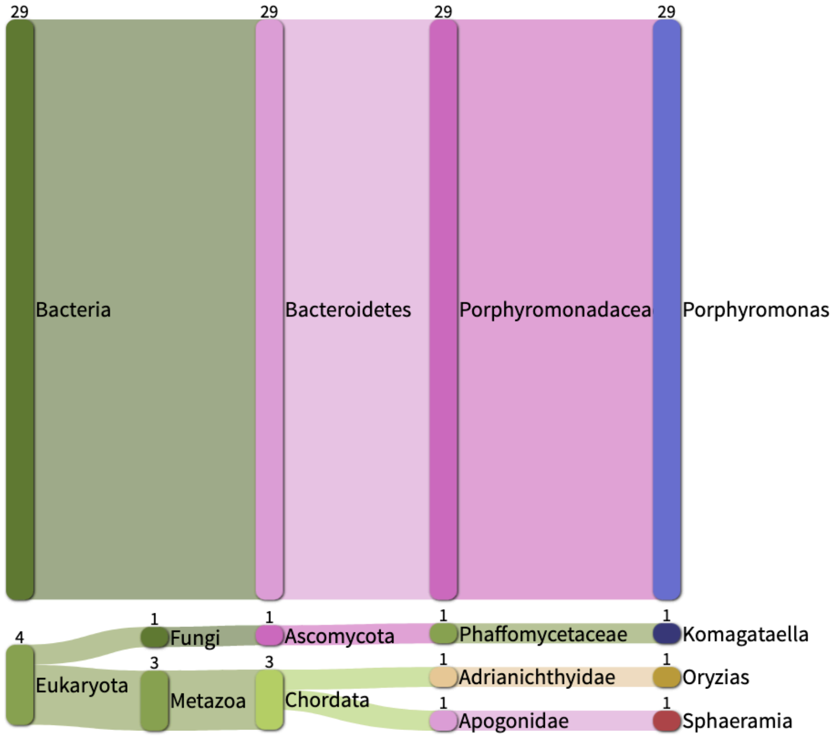

| Strain | % Classified | % *Porphyromonas* | Size (kb) | Conclusions | Kept in study |
|---|---|---|---|---|---|
| OH2963 | 100 | 65 | 2 116 475 | **Mainly *Porphyromonas*** | **YES** |

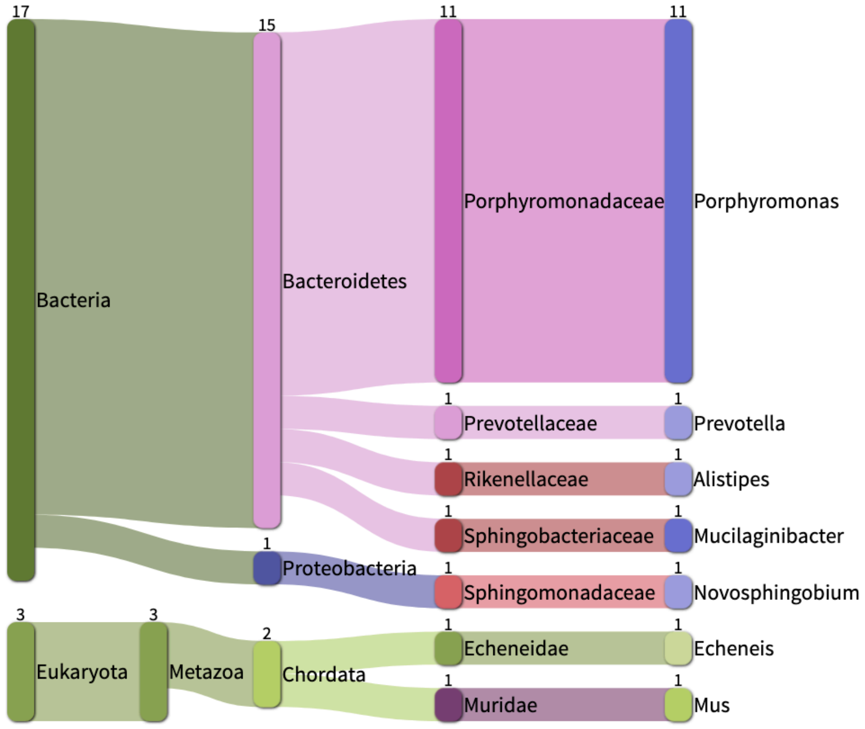

| Strain | % Classified | % *Porphyromonas* | Size (kb) | Conclusions | Kept in study |
|---|---|---|---|---|---|
| OH3588 | 100 | 67 | 2 117 249 | **Mainly *Porphyromonas*** | **YES** |

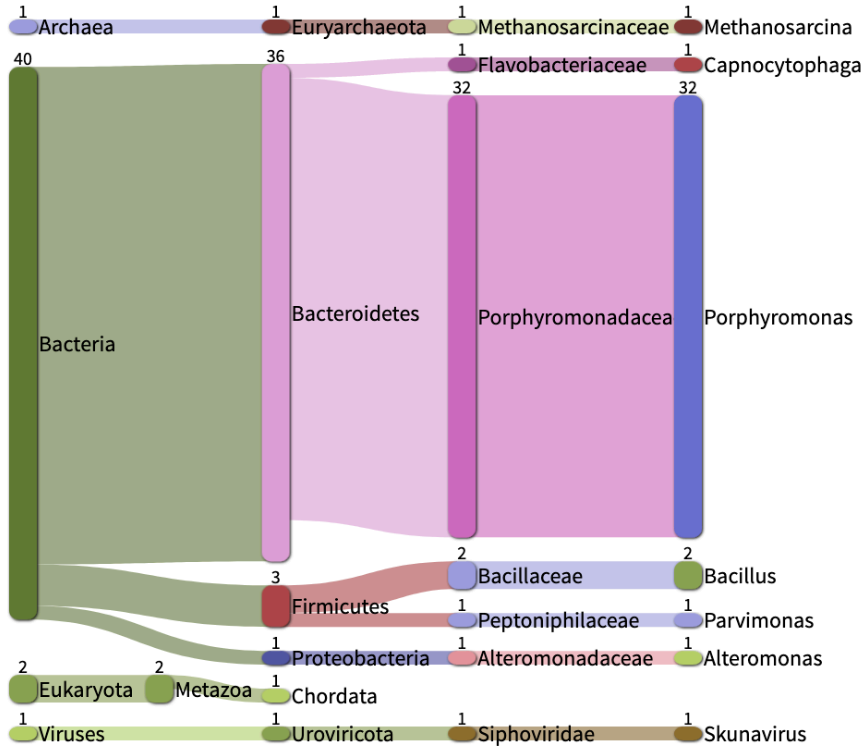

| Strain | % Classified | % *Porphyromonas* | Size (kb) | Conclusions | Kept in study |
|---|---|---|---|---|---|
| UEN 60-3 | 95.2 | 78 | 2 111 123 | **Mainly *Porphyromonas*** | **YES** |

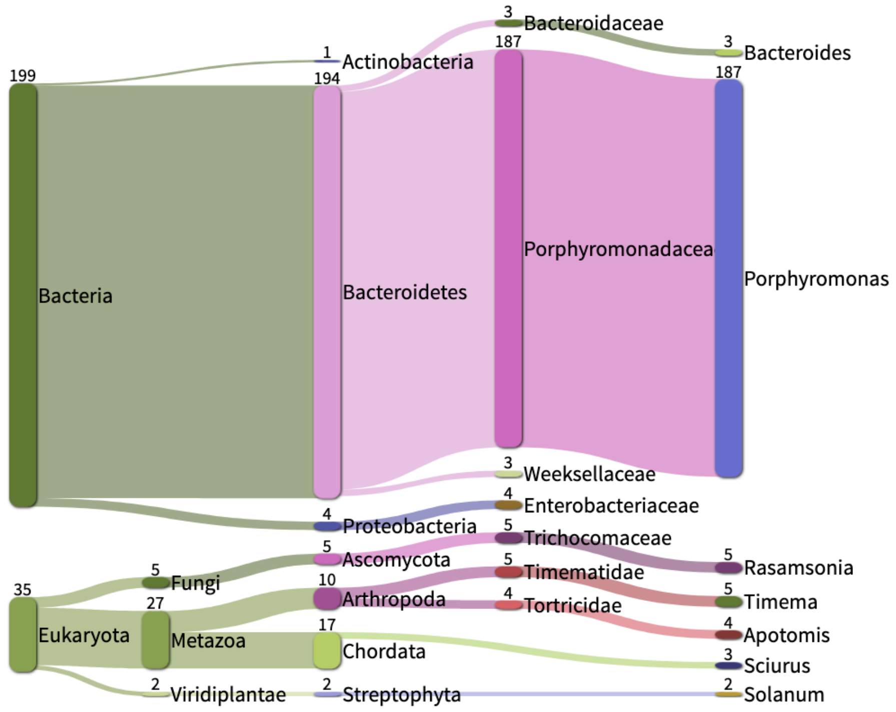

| Strain | % Classified | % *Porphyromonas* | Size (kb) | Conclusions | Kept in study |
|---|---|---|---|---|---|
| UMGS18 | 100 | **100** | 2 012 007 | ***Porphyromonas*** | **YES** |

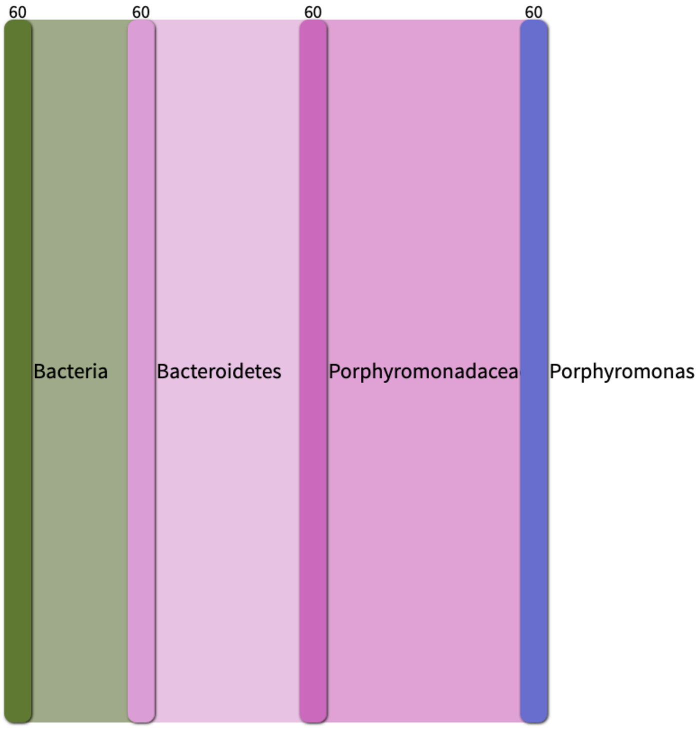

| Strain | % Classified | % *Porphyromonas* | Size (kb) | Conclusions | Kept in study |
|---|---|---|---|---|---|
| UMGS107 | 100 | **100** | 2 045 709 | ***Porphyromonas*** | **YES** |

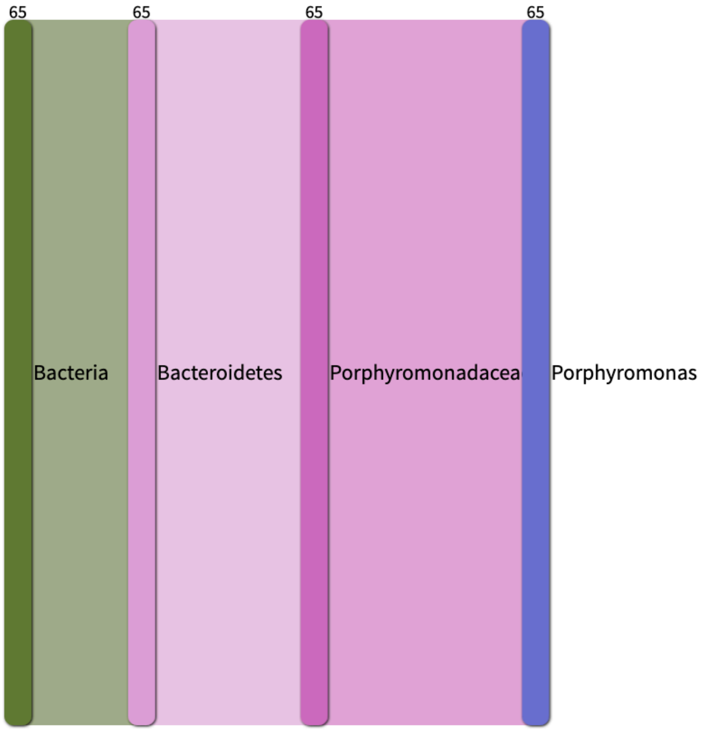

| Strain | % Classified | % *Porphyromonas* | Size (kb) | Conclusions | Kept in study |
|---|---|---|---|---|---|
| UMGS166 | 100 | **100** | 2 036 686 | ***Porphyromonas*** | **YES** |

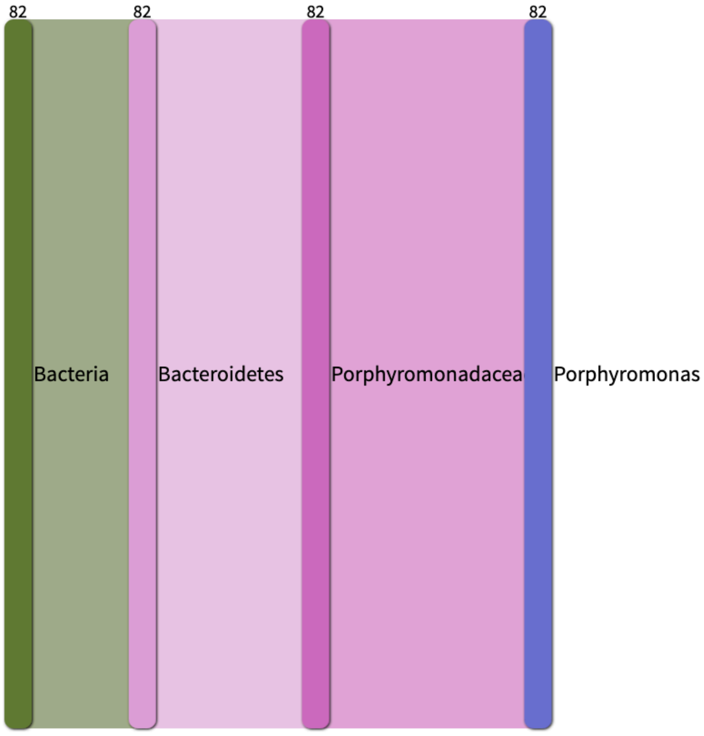

| Strain | % Classified | % *Porphyromonas* | Size (kb) | Conclusions | Kept in study |
|---|---|---|---|---|---|
| UMGS907 | 100 | **100** | 1 857 881 | ***Porphyromonas*** | **YES** |

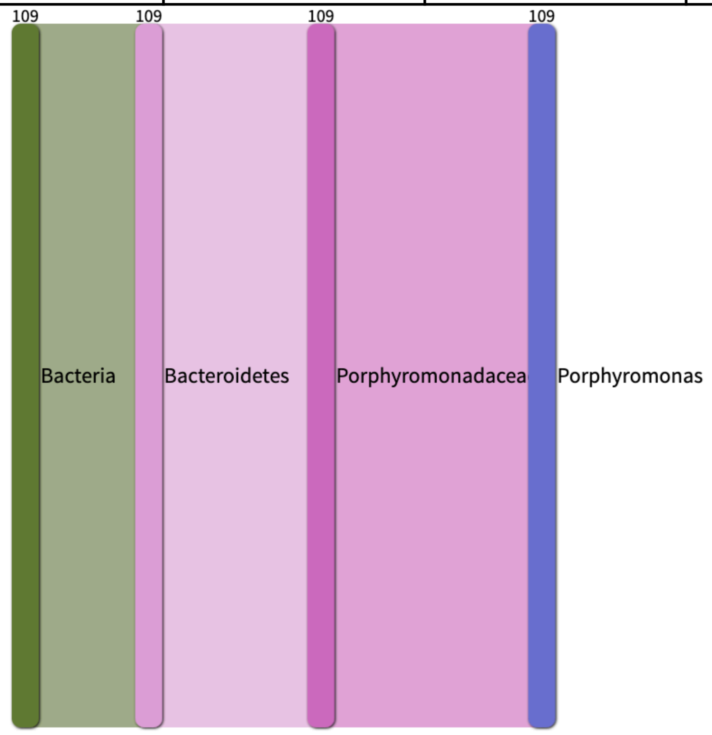

| Strain | % Classified | % *Porphyromonas* | Size (kb) | Conclusions | Kept in study |
|---|---|---|---|---|---|
| UMGS1085 | 100 | 97 | 1 452 709 | **Mainly *Porphyromonas*** | **YES** |

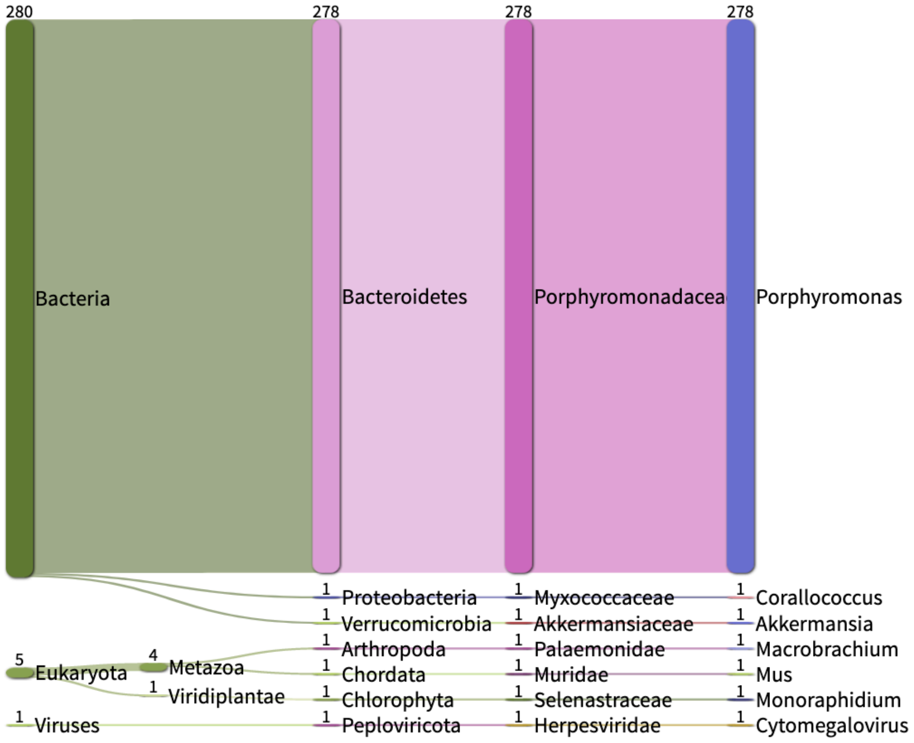

| Strain | % Classified | % *Porphyromonas* | Size (kb) | Conclusions | Kept in study |
|---|---|---|---|---|---|
| UMGS1452 | 100 | 97 | 1 616 451 | **Mainly *Porphyromonas*** | **YES** |

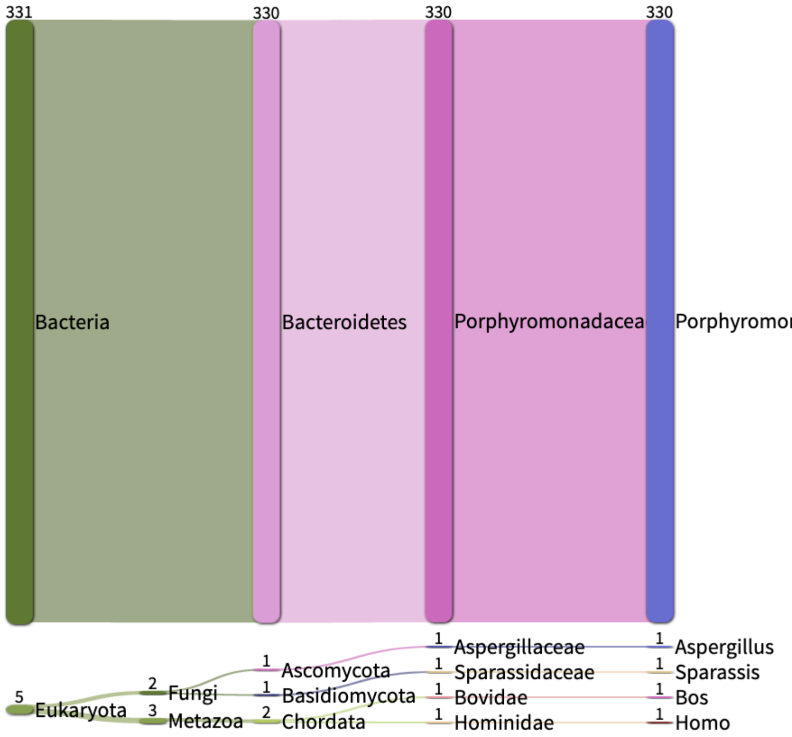

A

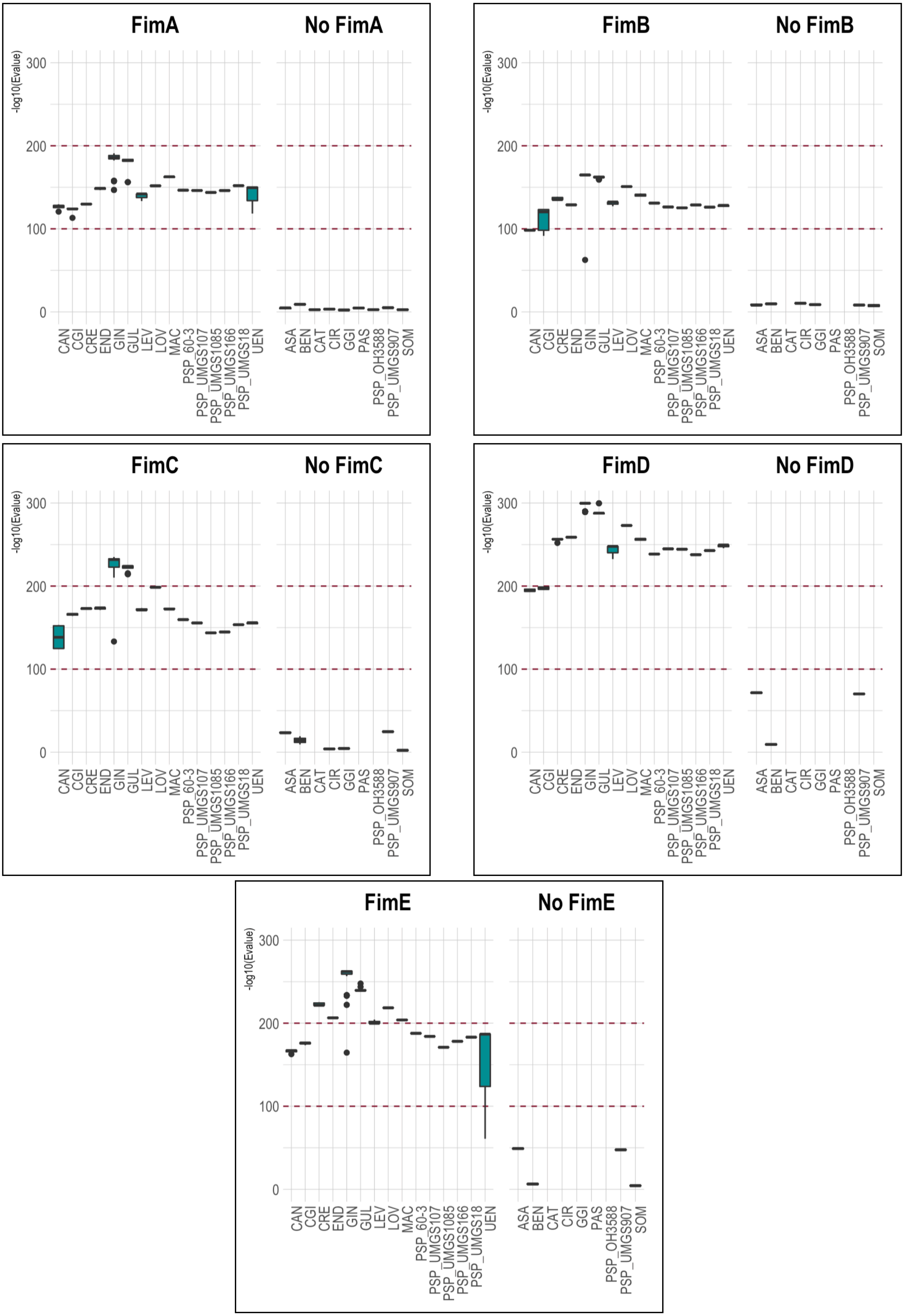

FimA
No FimA
FimB
No FimB
FimC
No FimC
FimD
No FimD
FimE
No FimE
-log10(Evalue)
300
200
100
0
CAN
CGI
CRE
END
GIN
GUL
LEV
LOV
MAC
PSP_60-3
PSP_UMGS107
PSP_UMGS1085
PSP_UMGS166
PSP_UMGS18
UEN
ASA
BEN
CAT
CIR
GGI
PAS
PSP_OH3588
PSP_UMGS907
SOM

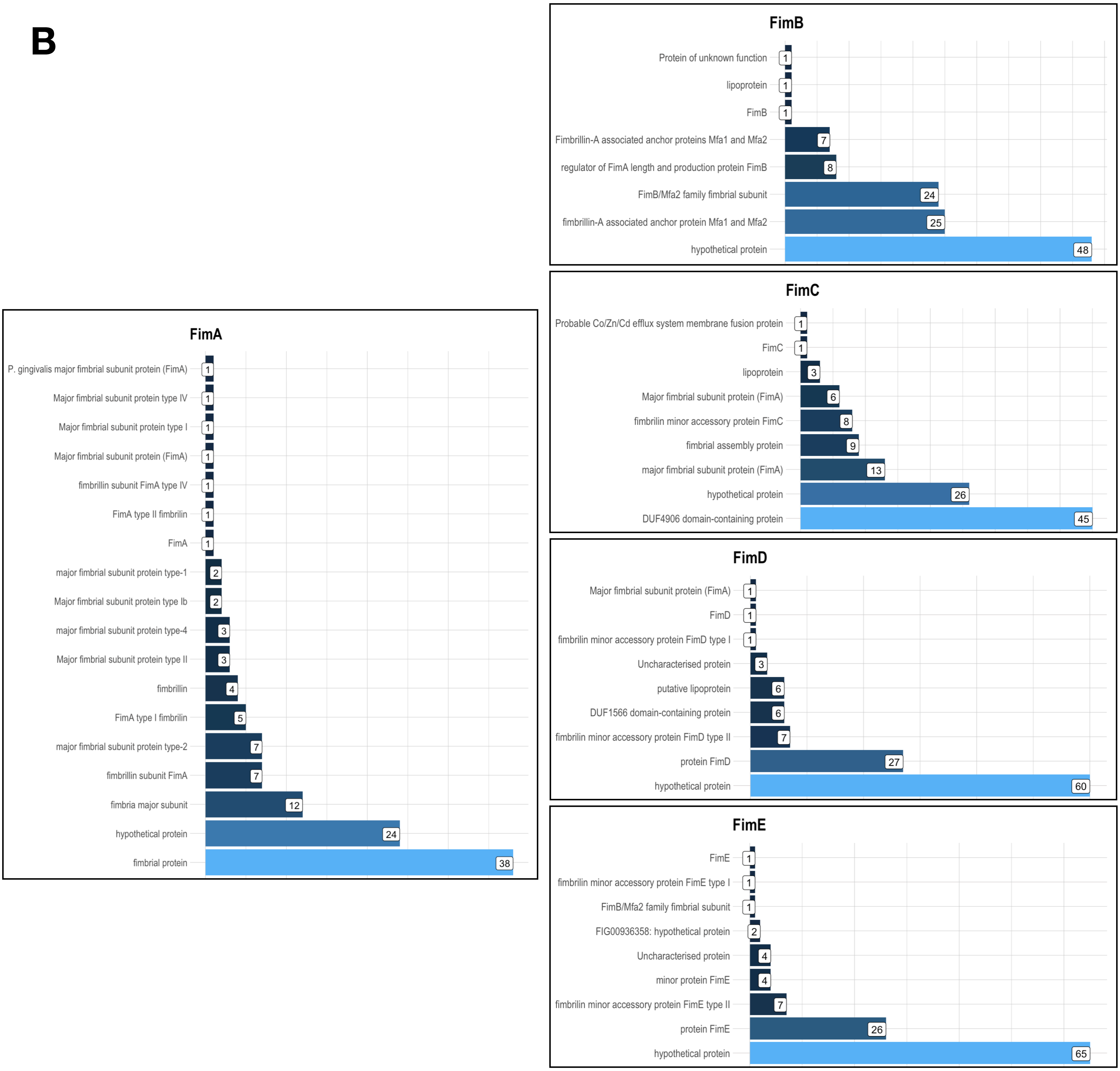
B
FimA
P. gingivalis major fimbrial subunit protein (FimA)
1
Major fimbrial subunit protein type IV
1
Major fimbrial subunit protein type I
1
Major fimbrial subunit protein (FimA)
1
fimbrillin subunit FimA type IV
1
FimA type II fimbrilin
1
FimA
1
major fimbrial subunit protein type-1
2
Major fimbrial subunit protein type Ib
2
major fimbrial subunit protein type-4
3
Major fimbrial subunit protein type II
3
fimbrillin
4
FimA type I fimbrilin
5
major fimbrial subunit protein type-2
7
fimbrillin subunit FimA
7
fimbria major subunit
12
hypothetical protein
24
fimbrial protein
38
FimB
Protein of unknown function
1
lipoprotein
1
FimB
1
Fimbrillin-A associated anchor proteins Mfa1 and Mfa2
7
regulator of FimA length and production protein FimB
8
FimB/Mfa2 family fimbrial subunit
24
fimbrillin-A associated anchor protein Mfa1 and Mfa2
25
hypothetical protein
48
FimC
Probable Co/Zn/Cd efflux system membrane fusion protein
1
FimC
1
lipoprotein
3
Major fimbrial subunit protein (FimA)
6
fimbrilin minor accessory protein FimC
8
fimbrial assembly protein
9
major fimbrial subunit protein (FimA)
13
hypothetical protein
26
DUF4906 domain-containing protein
45
FimD
Major fimbrial subunit protein (FimA)
1
FimD
1
fimbrilin minor accessory protein FimD type I
1
Uncharacterised protein
3
putative lipoprotein
6
DUF1566 domain-containing protein
6
fimbrilin minor accessory protein FimD type II
7
protein FimD
27
hypothetical protein
60
FimE
FimE
1
fimbrilin minor accessory protein FimE type I
1
FimB/Mfa2 family fimbrial subunit
1
FIG00936358: hypothetical protein
2
Uncharacterised protein
4
minor protein FimE
4
fimbrilin minor accessory protein FimE type II
7
protein FimE
26
hypothetical protein
65

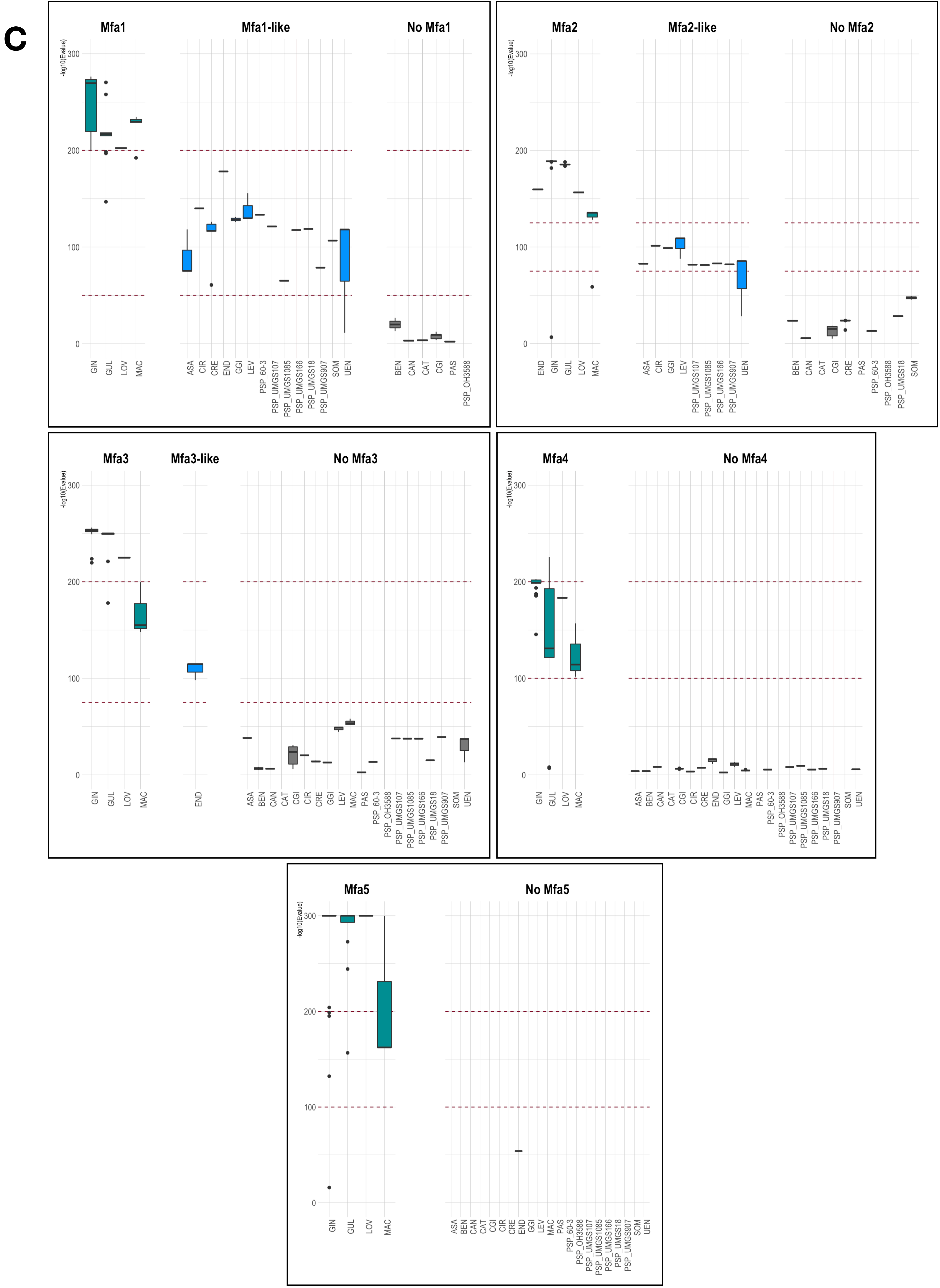
C
Mfa1
Mfa1-like
No Mfa1
Mfa2
Mfa2-like
No Mfa2
Mfa3
Mfa3-like
No Mfa3
Mfa4
No Mfa4
Mfa5
No Mfa5
-log10(Evalue)
0
100
200
300
GIN
GUL
LOV
MAC
ASA
CIR
CRE
END
GGI
LEV
PSP_60-3
PSP_UMGS107
PSP_UMGS1085
PSP_UMGS166
PSP_UMGS18
PSP_UMGS907
SOM
UEN
BEN
CAN
CAT
CGI
PAS
PSP_OH3588

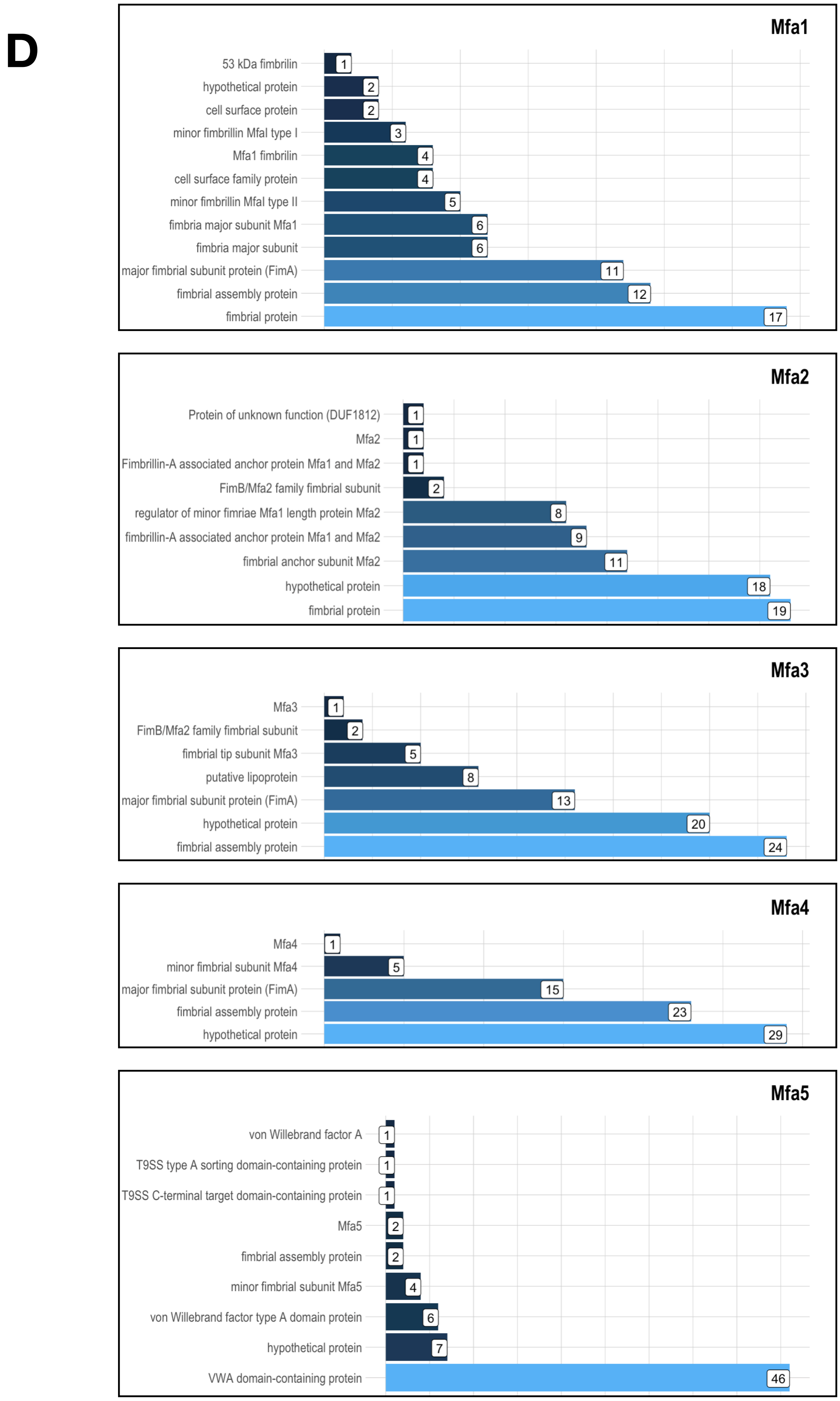
D
Mfa1
53 kDa fimbrilin
hypothetical protein
cell surface protein
minor fimbrillin Mfal type I
Mfa1 fimbrilin
cell surface family protein
minor fimbrillin Mfal type II
fimbria major subunit Mfa1
fimbria major subunit
major fimbrial subunit protein (FimA)
fimbrial assembly protein
fimbrial protein
1
2
2
3
4
4
5
6
6
11
12
17
Mfa2
Protein of unknown function (DUF1812)
Mfa2
Fimbrillin-A associated anchor protein Mfa1 and Mfa2
FimB/Mfa2 family fimbrial subunit
regulator of minor fimriae Mfa1 length protein Mfa2
fimbrillin-A associated anchor protein Mfa1 and Mfa2
fimbrial anchor subunit Mfa2
hypothetical protein
fimbrial protein
1
1
1
2
8
9
11
18
19
Mfa3
Mfa3
FimB/Mfa2 family fimbrial subunit
fimbrial tip subunit Mfa3
putative lipoprotein
major fimbrial subunit protein (FimA)
hypothetical protein
fimbrial assembly protein
1
2
5
8
13
20
24
Mfa4
Mfa4
minor fimbrial subunit Mfa4
major fimbrial subunit protein (FimA)
fimbrial assembly protein
hypothetical protein
1
5
15
23
29
Mfa5
von Willebrand factor A
T9SS type A sorting domain-containing protein
T9SS C-terminal target domain-containing protein
Mfa5
fimbrial assembly protein
minor fimbrial subunit Mfa5
von Willebrand factor type A domain protein
hypothetical protein
VWA domain-containing protein
1
1
1
2
2
4
6
7
46

E

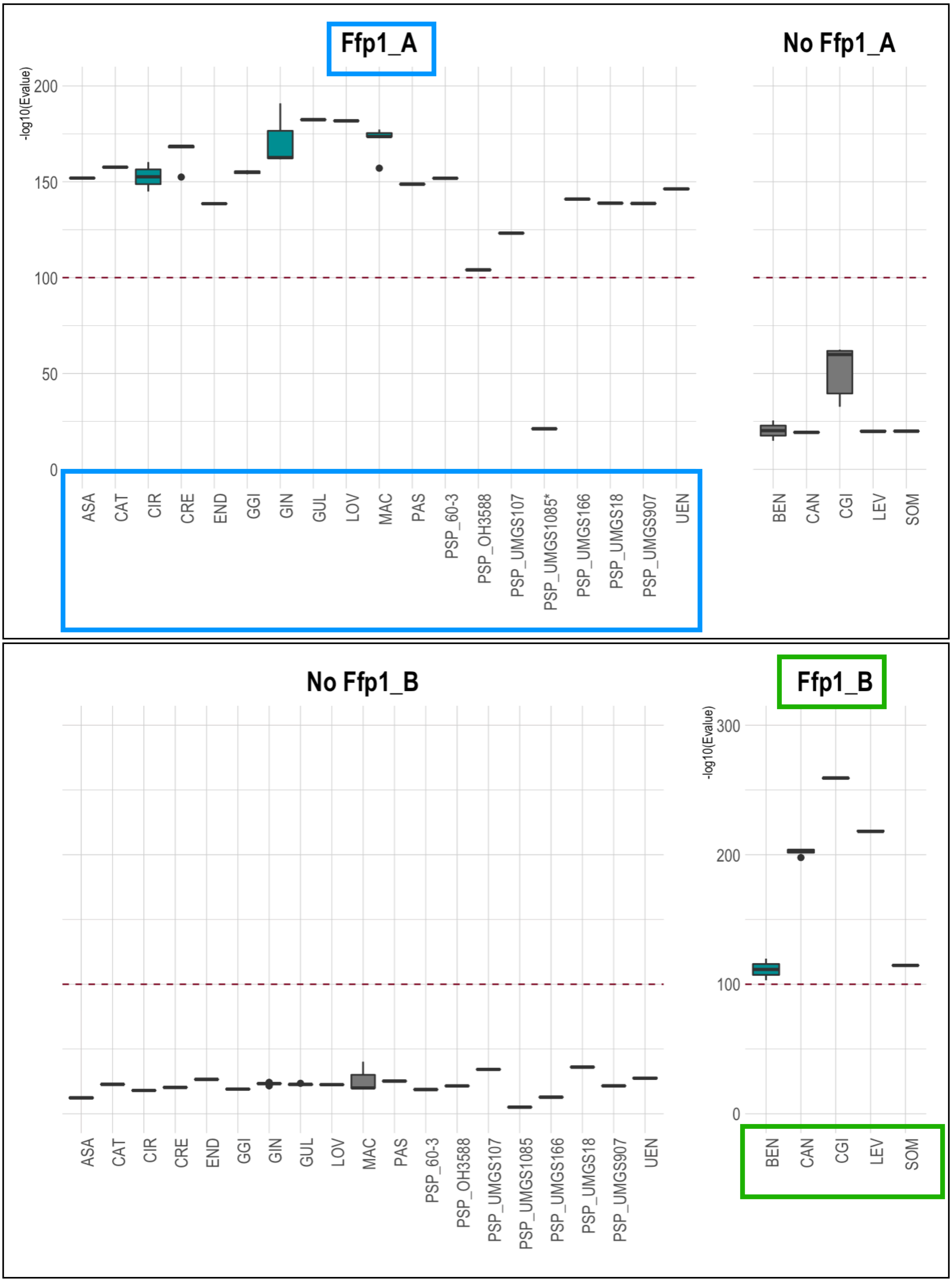
Ffp1_A
No Ffp1_A
-log10(Evalue)
200
150
100
50
0
ASA
CAT
CIR
CRE
END
GGI
GIN
GUL
LOV
MAC
PAS
PSP_60-3
PSP_OH3588
PSP_UMGS107
PSP_UMGS1085*
PSP_UMGS166
PSP_UMGS18
PSP_UMGS907
UEN
BEN
CAN
CGI
LEV
SOM
No Ffp1_B
Ffp1_B
300
200
100
0
ASA
CAT
CIR
CRE
END
GGI
GIN
GUL
LOV
MAC
PAS
PSP_60-3
PSP_OH3588
PSP_UMGS107
PSP_UMGS1085
PSP_UMGS166
PSP_UMGS18
PSP_UMGS907
UEN
BEN
CAN
CGI
LEV
SOM

F

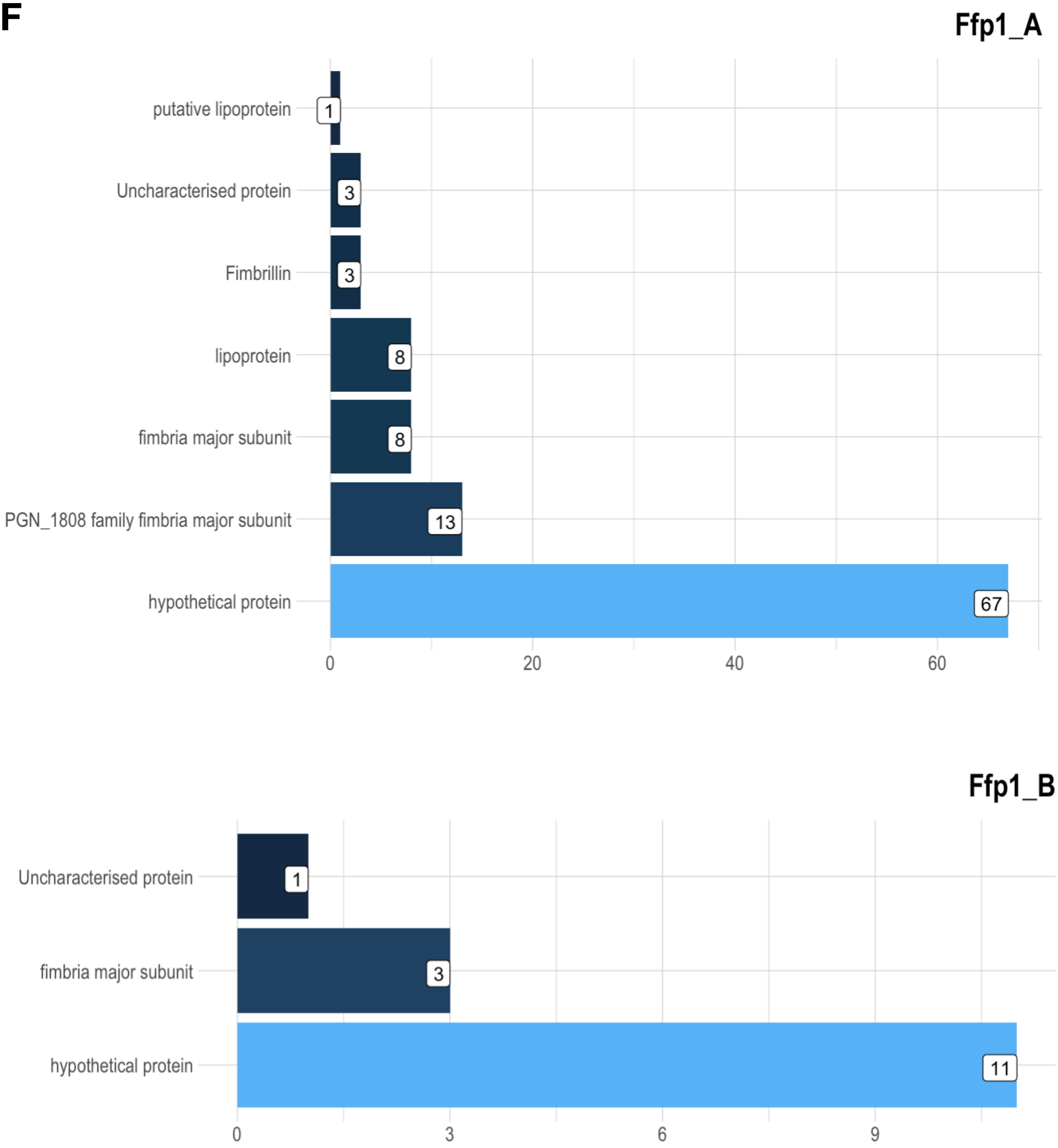
Ffp1_A
putative lipoprotein
Uncharacterised protein
Fimbrillin
lipoprotein
fimbria major subunit
PGN_1808 family fimbria major subunit
hypothetical protein
1
3
3
8
8
13
67
0
20
40
60
Ffp1_B
Uncharacterised protein
fimbria major subunit
hypothetical protein
1
3
11
0
3
6
9

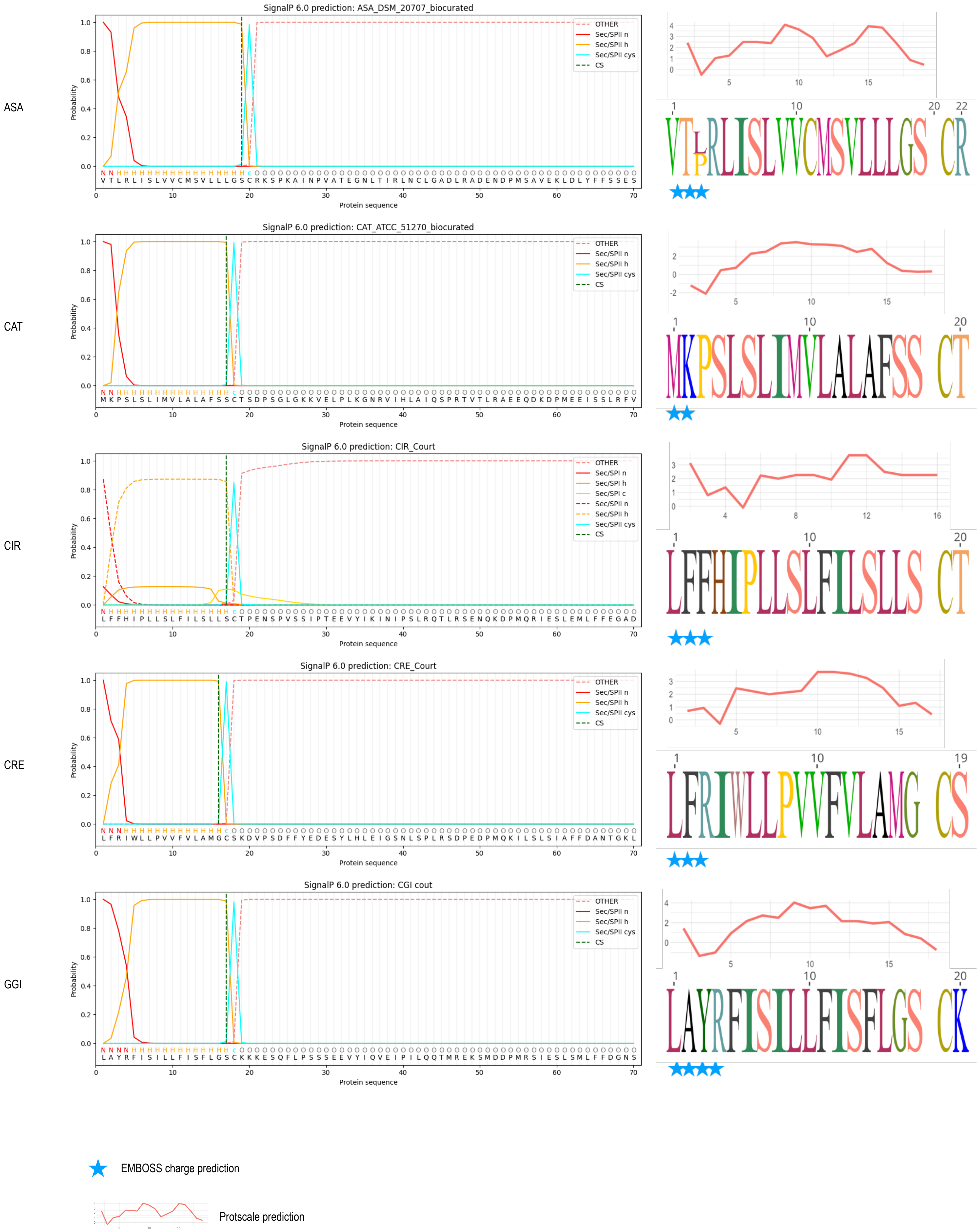

SignalP 6.0 prediction: ASA_DSM_20707_biocurated
ASA
SignalP 6.0 prediction: CAT_ATCC_51270_biocurated
CAT
SignalP 6.0 prediction: CIR_Court
CIR
SignalP 6.0 prediction: CRE_Court
CRE
SignalP 6.0 prediction: CGI cout
GGI
Probability
Protein sequence
OTHER
Sec/SPII n
Sec/SPII h
Sec/SPII cys
Sec/SPI c
CS
EMBOSS charge prediction
Protscale prediction

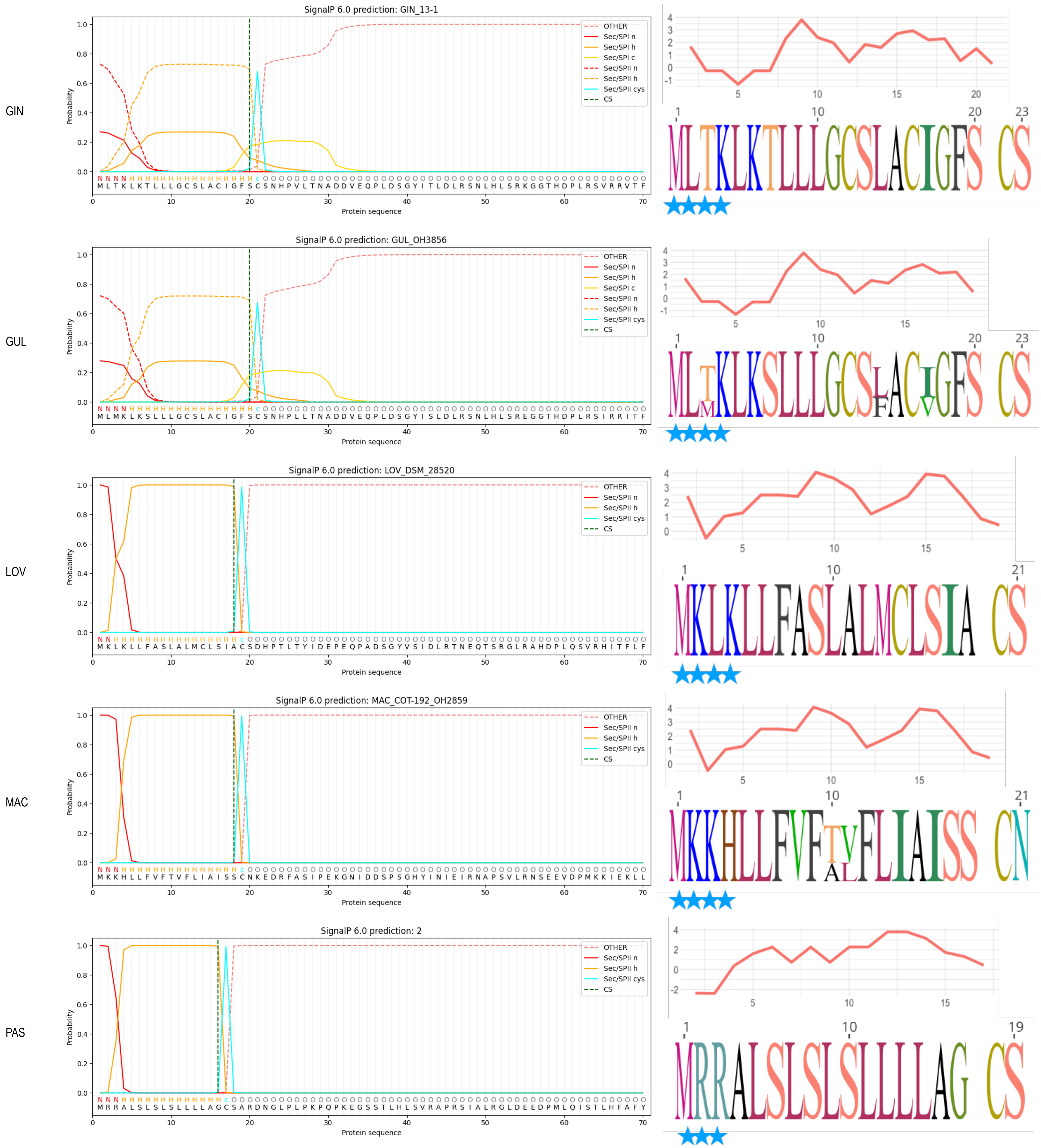

GIN
SignalP 6.0 prediction: GIN_13-1
OTHER
Sec/SPI n
Sec/SPI h
Sec/SPI c
Sec/SPII n
Sec/SPII h
Sec/SPII cys
CS
Probability
Protein sequence
M L T K L K T L L L G C S L A C I G F S C S N H P V L T N A D D V E Q P L D S G Y I T L D L R S N L H L S R K G G T H D P L R S V R R V T F
GUL
SignalP 6.0 prediction: GUL_OH3856
M L M K L K S L L L G C S L A C I G F S C S N H P L L T N A D D V E Q P L D S G Y I S L D L R S N L H L S R E G G T H D P L R S I R R I T F
LOV
SignalP 6.0 prediction: LOV_DSM_28520
M K L K L L F A S L A L M C L S I A C S D H P T L T Y I D E P E Q P A D S G Y V S I D L R T N E Q T S R G L R A H D P L Q S V R H I T F L F
MAC
SignalP 6.0 prediction: MAC_COT-192_OH2859
M K K H L L F V F T V F L I A I S S C N K E D R F A S I P E K G N I D D S P S G H Y I N I E I R N A P S V L R N S E E V D P M K K I E K L L
PAS
SignalP 6.0 prediction: 2
M R R A L S L S L S L L L L A G C S A R D N G L P L P K P Q P K E G S S T L H L S V R A P R S I A L R G L D E E D P M L Q I S T L H F A F Y

60-3

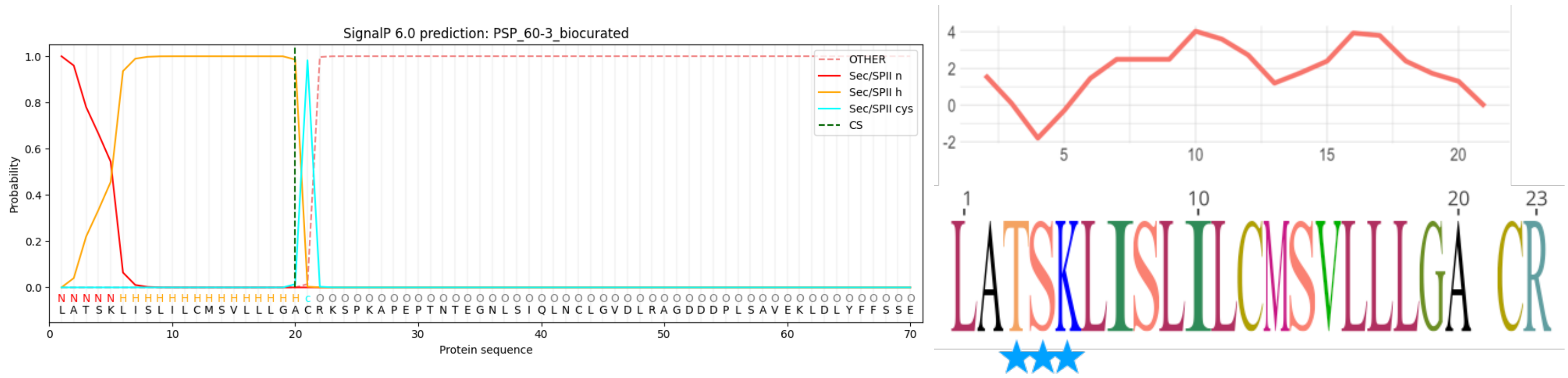

OH3588

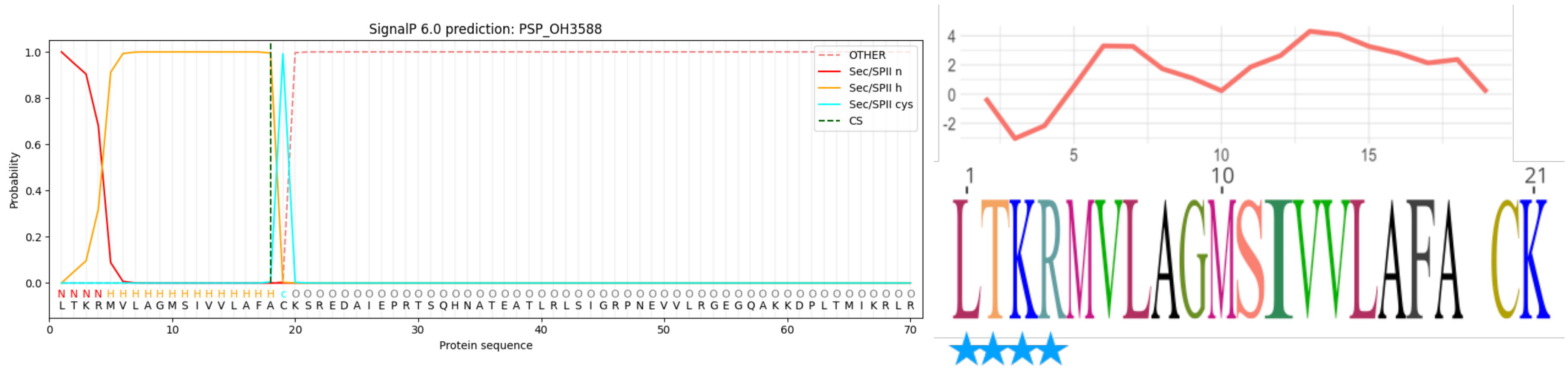

UMGS18

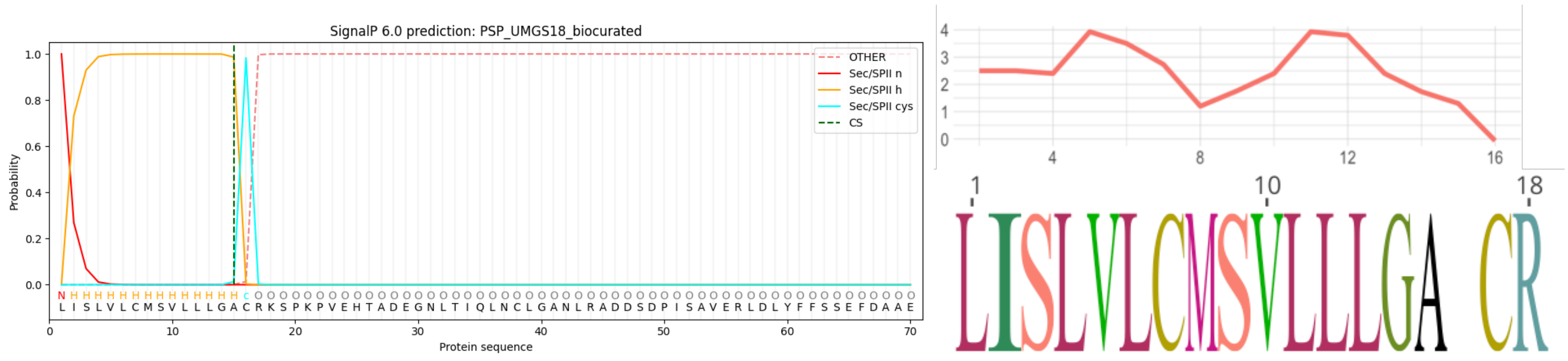

UMGS107

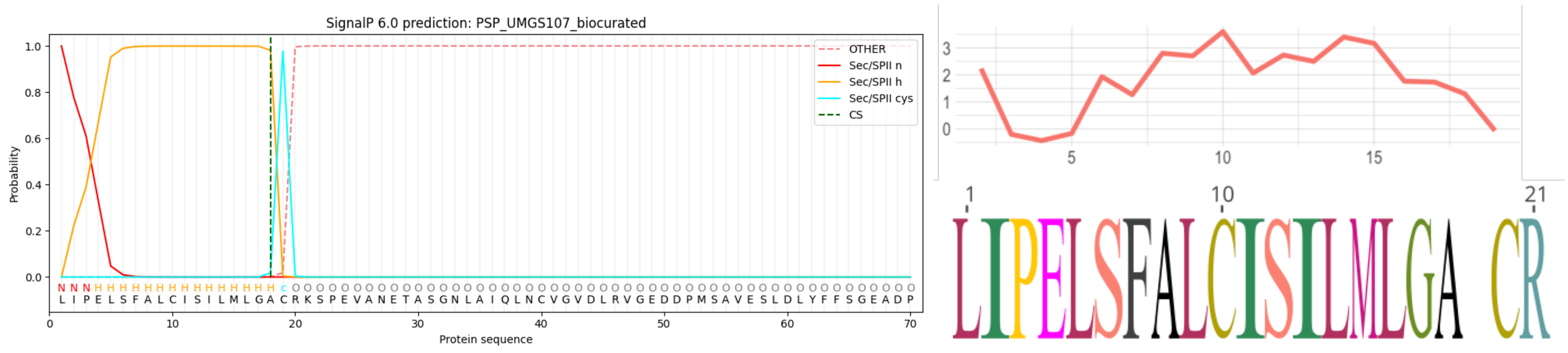

UMGS166

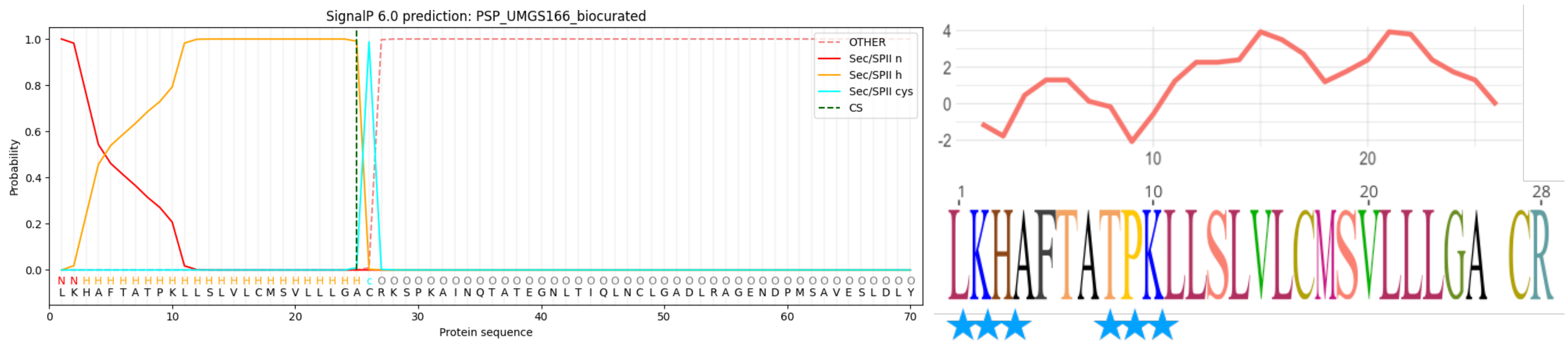

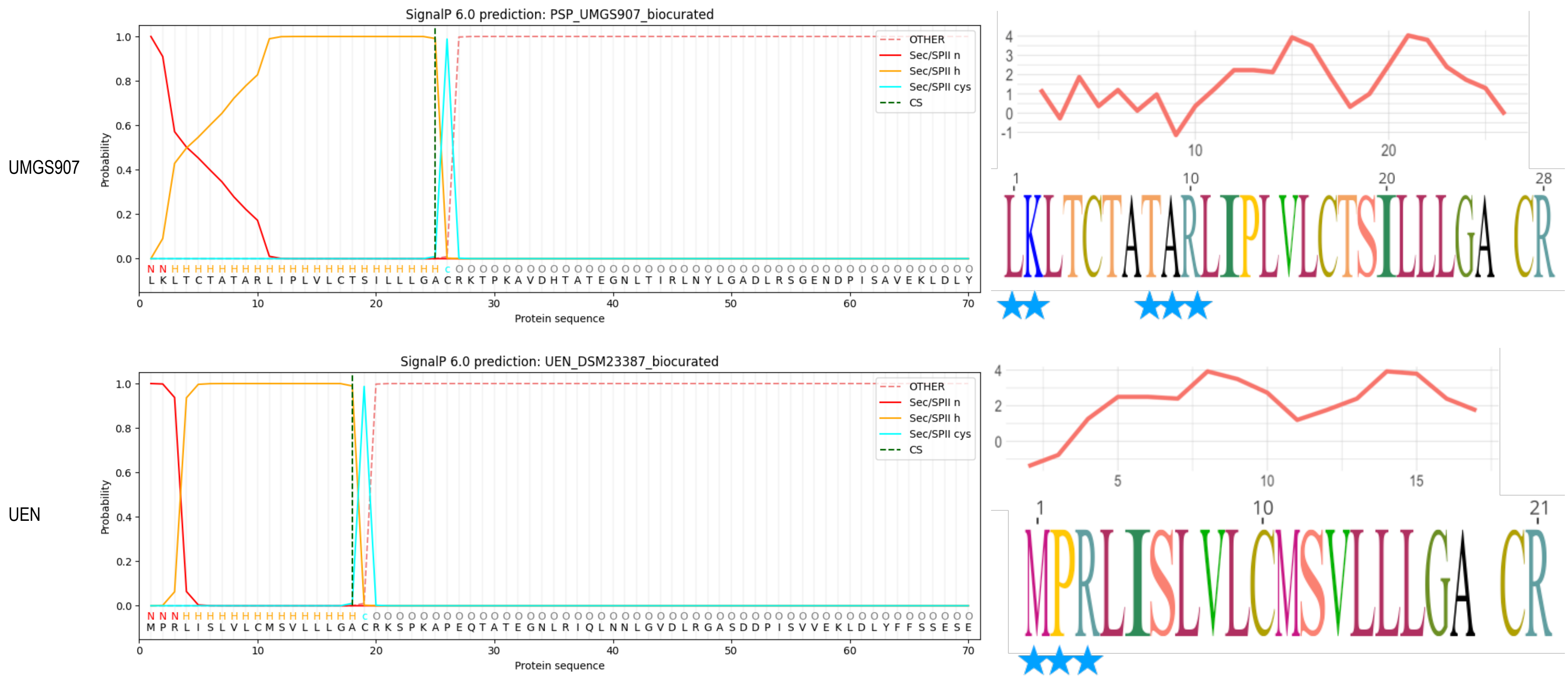

- **Sec/SPII**: lipoprotein signal peptides transported by the Sec translocon and cleaved by Signal Peptidase II (Lsp)

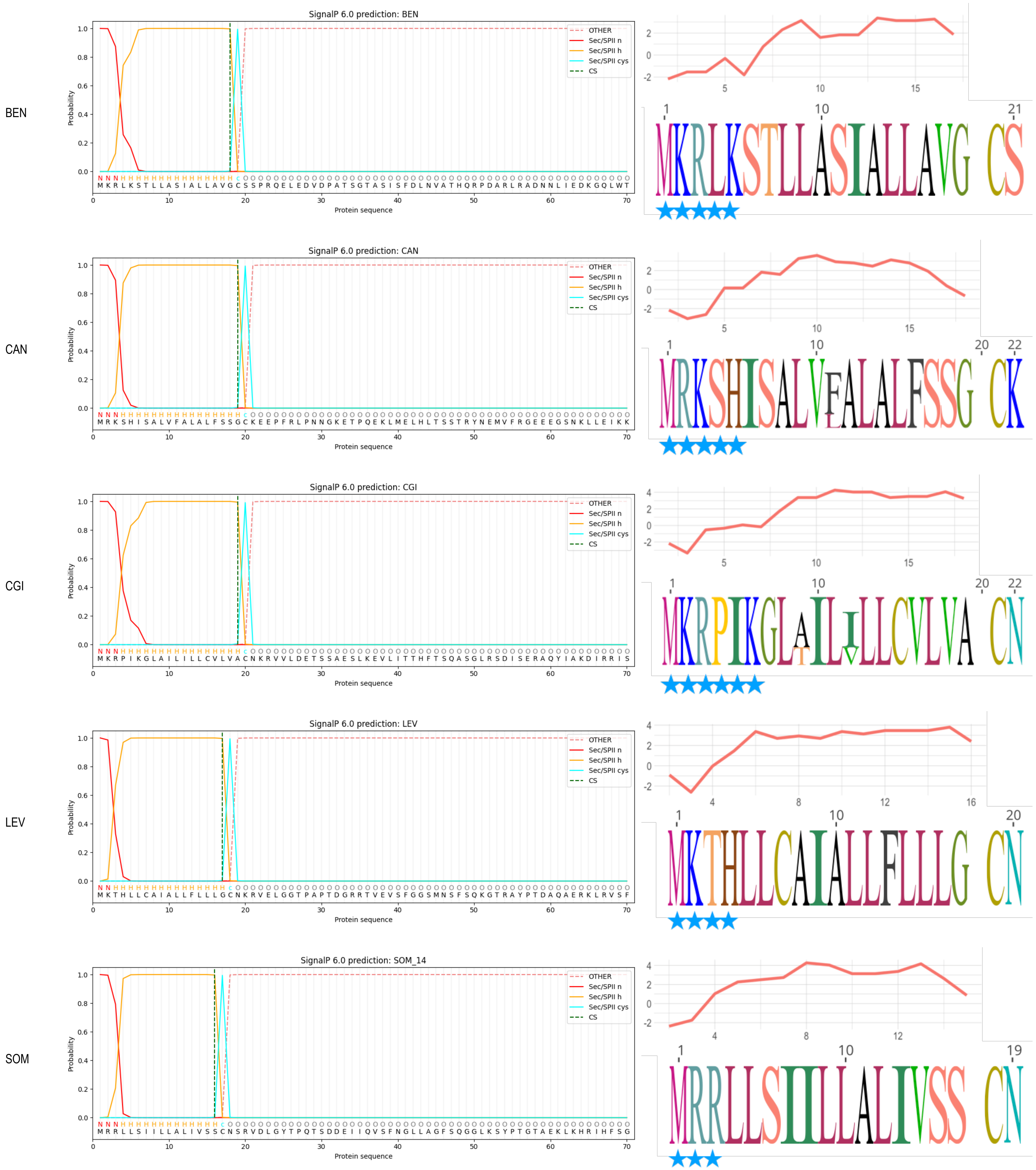

BEN
SignalP 6.0 prediction: BEN
M K R L K S T L L A S I A L L A V G C S S P R Q E L E D V D P A T S G T A S I S F D L N V A T H Q R P D A R L R A D N N L I E D K G Q L W T
MKRLKSTLLASIALLAVG CS
CAN
SignalP 6.0 prediction: CAN
M R K S H I S A L V F A L A L F S S G C K E E P F R L P N N G K E T P Q E K L M E L H L T S S T R Y N E M V F R G E E E G S N K L L E I K K
MRKSHISALVFALALFSSG CK
CGI
SignalP 6.0 prediction: CGI
M K R P I K G L A I L I L L C V L V A C N K R V V L D E T S S A E S L K E V L I T T H F T S Q A S G L R S D I S E R A Q Y I A K D I R R I S
MKRPIKGLAILILLCVLVA CN
LEV
SignalP 6.0 prediction: LEV
M K T H L L C A I A L L F L L L G C N K R V E L G G T P A P T D G R R T V E V S F G G S M N S F S Q K G T R A Y P T D A Q A E R K L R V S F
MKTHLLCAIALLFLLLG CN
SOM
SignalP 6.0 prediction: SOM_14
M R R L L S I I L L A L I V S S C N S R V D L G Y T P Q T S D D E I I Q V S F N G L L A G F S Q G G L K S Y P T G T A E K L K H R I H F S G
MRRLLSIILLALIVSS CN
OTHER
Sec/SPII n
Sec/SPII h
Sec/SPII cys
CS
Probability
Protein sequence

| Group | iPBA results | Phyre2 results | Superposition of predicted structure (green) with model structure (c4jrfA in red) using iPBA |
|---|---|---|---|
| ASA | Normalized score 90.90<br>RMSD 2.27<br>Alignment length 653<br>Aligned residues 339<br>Fraction aligned 51.91 %<br>GDT TS 33.25 | 430 residues (88% of sequence) have been modelled with 100.0% confidence | 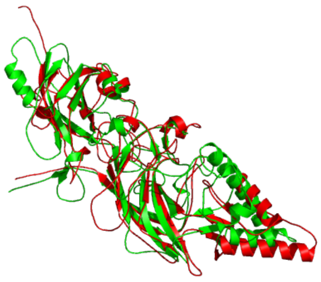 |
| CAT | Normalized score 122.53<br>RMSD 2.09<br>Alignment length 622<br>Aligned residues 341<br>Fraction aligned 54.82 %<br>GDT TS 36.58 | 402 residues (88% of sequence) have been modelled with 100.0% | 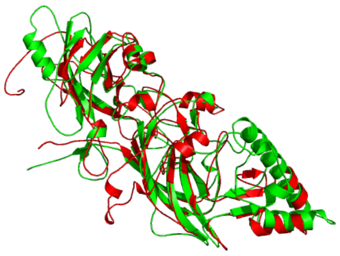 |
| CIR | Normalized score 58.01<br>RMSD 2.25<br>Alignment length 631<br>Aligned residues 350<br>Fraction aligned 55.47 %<br>GDT TS 35.97 | 424 residues (89% of sequence) have been modelled with 100.0% confidence | 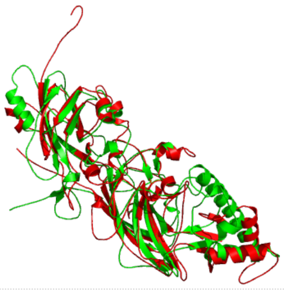 |
| CRE | Normalized score 50.92<br>RMSD 2.42<br>Alignment length 636<br>Aligned residues 334<br>Fraction aligned 52.52 %<br>GDT TS 31.56 | 418 residues (90% of sequence) have been modelled with 100.0% confidence | 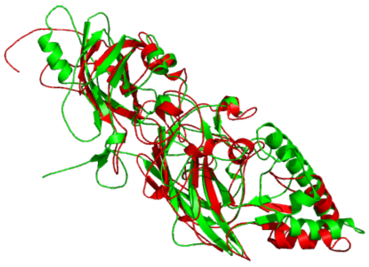 |
| END | Normalized score 57.54<br>RMSD 2.18<br>Alignment length 636<br>Aligned residues 339<br>Fraction aligned 53.30 %<br>GDT TS 35.29 | 424 residues (90% of sequence) have been modelled with 100.0% confidence | 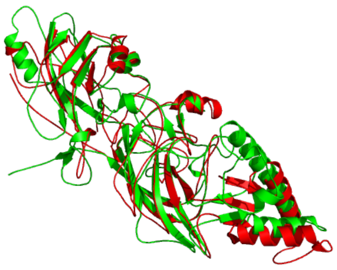 |
| GGI | Normalized score 131.46<br>RMSD 2.13<br>Alignment length 629<br>Aligned residues 350<br>Fraction aligned 55.64 %<br>GDT TS 37.30 | 427 residues (90% of sequence) have been modelled with 100.0% confidence | 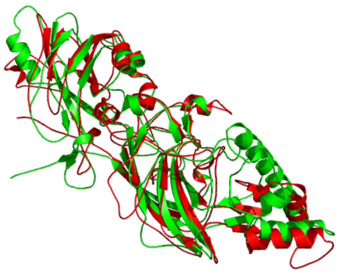 |
| GIN | Normalized score 57.00<br>RMSD 2.20<br>Alignment length 637<br>Aligned residues 329<br>Fraction aligned 51.65 %<br>GDT TS 33.69 | 413 residues (90% of sequence) have been modelled with 100.0% confidence | 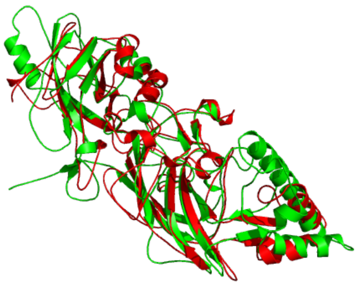 |
| GUL | Normalized score 53.22<br>RMSD 2.20<br>Alignment length 647<br>Aligned residues 319<br>Fraction aligned 49.30 %<br>GDT TS 32.46 | 412 residues (90% of sequence) have been modelled with 100.0% confidence | 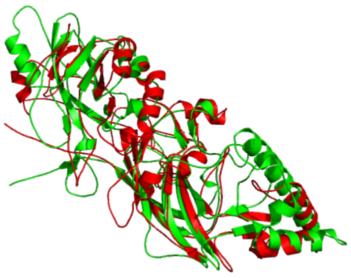 |

| Group | iPBA results | Phyre2 results | Superposition of predicted structure (green) with model structure (c4jrfA in red) using iPBA |
|---|---|---|---|
| LOV | Normalized score 65.21<br>RMSD 2.27<br>Alignment length 659<br>Aligned residues 309<br>Fraction aligned 46.89 %<br>GDT TS 29.68 | 413 residues (89% of sequence) have been modelled with 100.0% confidence | 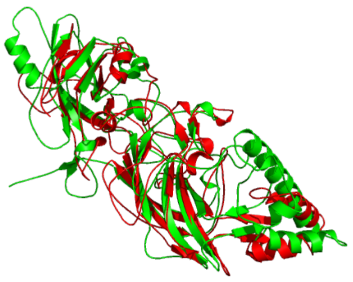 |
| MAC | Normalized score 27.70<br>RMSD 2.27<br>Alignment length 635<br>Aligned residues 331<br>Fraction aligned 52.13 %<br>GDT TS 33.33 | 411 residues (89% of sequence) have been modelled with 100.0% confidence | 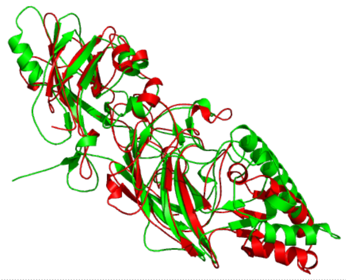 |
| PAS | Normalized score 113.17<br>RMSD 2.29<br>Alignment length 636<br>Aligned residues 329<br>Fraction aligned 51.73 %<br>GDT TS 32.58 | 402 residues (88% of sequence) have been modelled with 100.0% confidence | 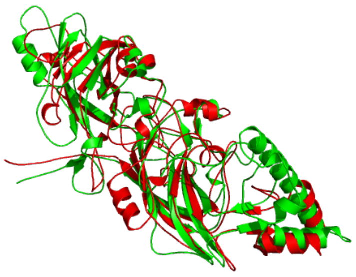 |
| PSP_60-3 | Normalized score 108.84<br>RMSD 2.24<br>Alignment length 657<br>Aligned residues 334<br>Fraction aligned 50.84 %<br>GDT TS 32.83 | 430 residues (89% of sequence) have been modelled with 100.0% confidence | 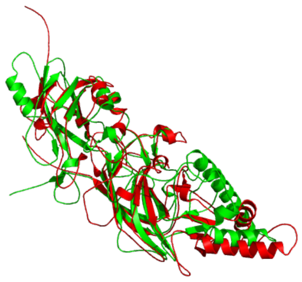 |
| PSP_OH3588 | Normalized score 57.31<br>RMSD 2.28<br>Alignment length 634<br>Aligned residues 327<br>Fraction aligned 51.58 %<br>GDT TS 32.72 | 403 residues (89% of sequence) have been modelled with 100.0% confidence | 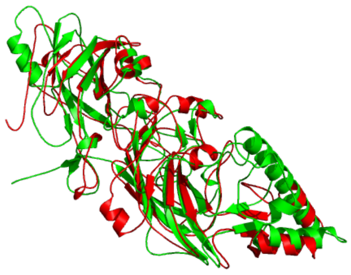 |
| PSP_UMGS18 | Normalized score 27.91<br>RMSD 2.22<br>Alignment length 663<br>Aligned residues 328<br>Fraction aligned 49.47 %<br>GDT TS 32.00 | 434 residues (89% of sequence) have been modelled with 100.0% confidence | 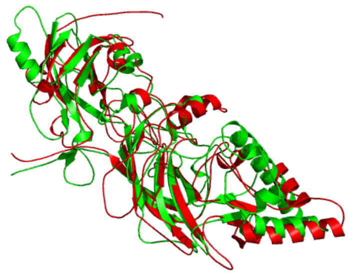 |
| PSP_UMGS107 | Normalized score 58.68<br>RMSD 2.22<br>Alignment length 654<br>Aligned residues 337<br>Fraction aligned 51.53 %<br>GDT TS 33.07 | 435 residues (90% of sequence) have been modelled with 100.0% confidence | 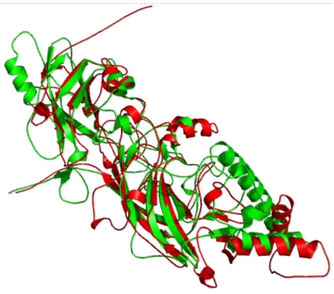 |
| PSP_UMGS166 | Normalized score 83.53<br>RMSD 2.28<br>Alignment length 657<br>Aligned residues 332<br>Fraction aligned 50.53 %<br>GDT TS 32.02 | 430 residues (89% of sequence) have been modelled with 100.0% confidence | 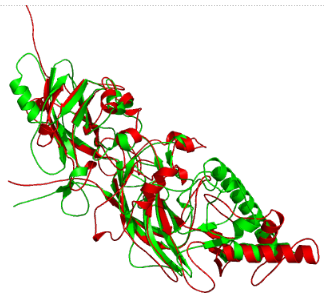 |

| Group | iPBA results | Phyre2 results | Superposition of predicted structure (green) with model structure (c4jrfA in red) using iPBA |
|---|---|---|---|
| PSP_UMGS907 | Normalized score 71.84<br>RMSD 2.19<br>Alignment length 664<br>Aligned residues 327<br>Fraction aligned 49.25 %<br>GDT TS 31.97 | 431 residues (89% of sequence) have been modelled with 100.0% confidence | 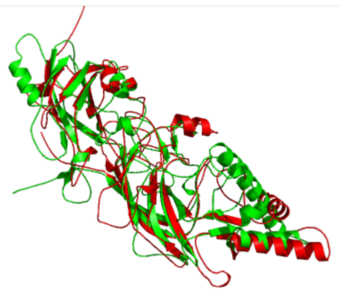 |
| UEN | Normalized score -109.10<br>RMSD 2.39<br>Alignment length 668<br>Aligned residues 322<br>Fraction aligned 48.20 %<br>GDT TS 29.76 | 429 residues (89% of sequence) have been modelled with 100.0% confidence | 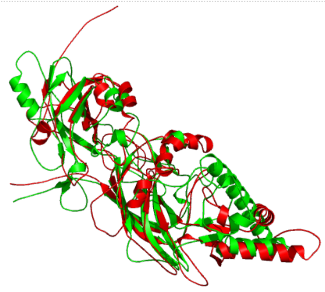 |
| BEN | Normalized score 66.46<br>RMSD 2.25<br>Alignment length 682<br>Aligned residues 331<br>Fraction aligned 48.53 %<br>GDT TS 31.24 | 438 residues (86% of sequence) have been modelled with 100.0% confidence | 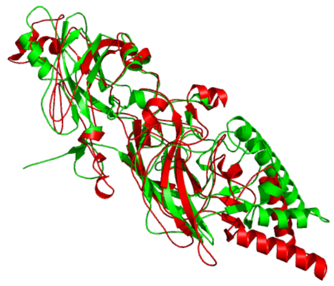 |
| CAN | Normalized score 31.90<br>RMSD 2.28<br>Alignment length 677<br>Aligned residues 336<br>Fraction aligned 49.63 %<br>GDT TS 31.45 | 392 residues (77% of sequence) have been modelled with 98.9% confidence | 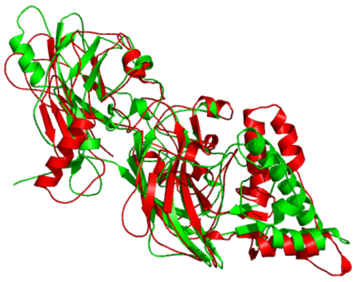 |
| CGI | Normalized score -3.48<br>RMSD 2.35<br>Alignment length 640<br>Aligned residues 347<br>Fraction aligned 54.22 %<br>GDT TS 33.63 | 421 residues ( 88% of your sequence) have been modelled with 100.0% confidence | 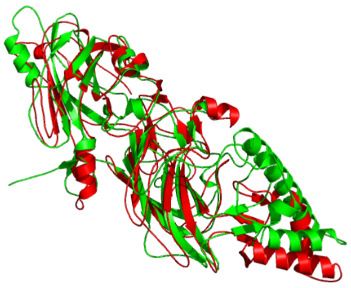 |
| LEV | Normalized score 60.67<br>RMSD 2.53<br>Alignment length 673<br>Aligned residues 329<br>Fraction aligned 48.89 %<br>GDT TS 28.77 | 409 residues (82% of sequence) have been modelled with 100.0% confidence | 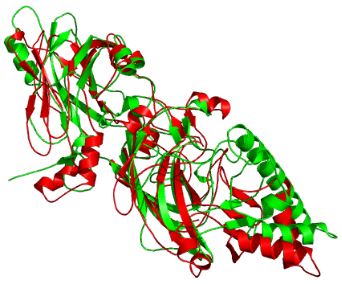 |
| SOM | Normalized score 59.26<br>RMSD 2.37<br>Alignment length 647<br>Aligned residues 347<br>Fraction aligned 53.63 %<br>GDT TS 33.14 | 186 residues (38% of sequence) have been modelled with 100.0% confidence | 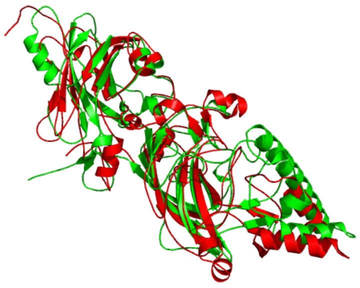 |

BACOVA

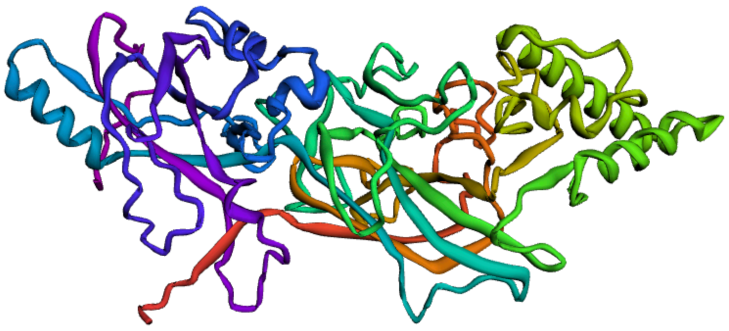

GIN_FIMA

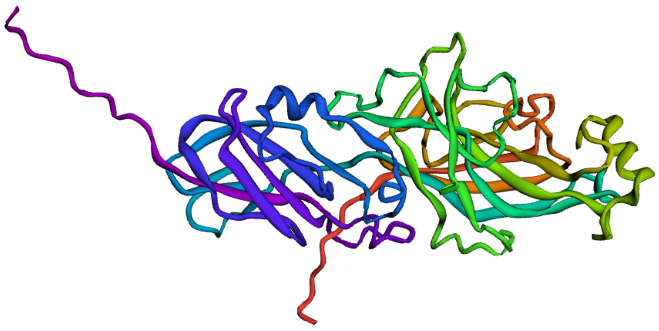

GIN_MFA1

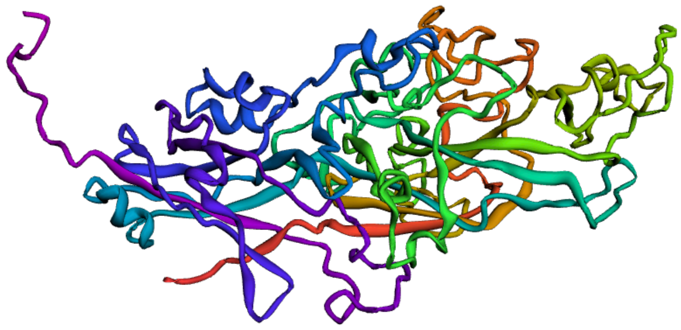

GIN_FFP1

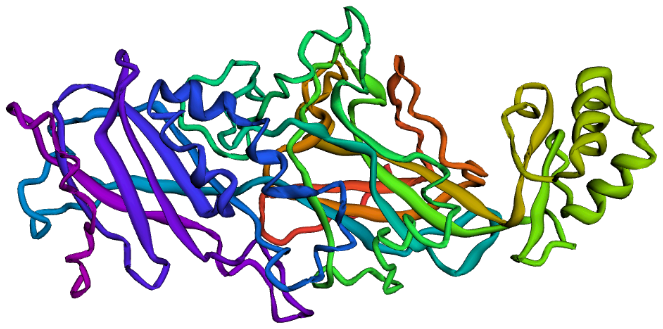

GIN_FFP1/BACOVA

| | |
|---|---|
| Normalized score | 57.00 |
| RMSD | 2.20 |
| Alignment length | 637 |
| Aligned residues | 329 |
| Fraction aligned | 51.65 % |
| GDT TS | 33.69 |

GIN_FFP1/FIMA

| | |
|---|---|
| Normalized score | -23.34 |
| RMSD | 2.48 |
| Alignment length | 588 |
| Aligned residues | 243 |
| Fraction aligned | 41.33 % |
| GDT TS | 24.56 |

GIN_FFP1/MFA1

| | |
|---|---|
| Normalized score | 18.18 |
| RMSD | 2.17 |
| Alignment length | 645 |
| Aligned residues | 321 |
| Fraction aligned | 49.77 % |
| GDT TS | 32.68 |

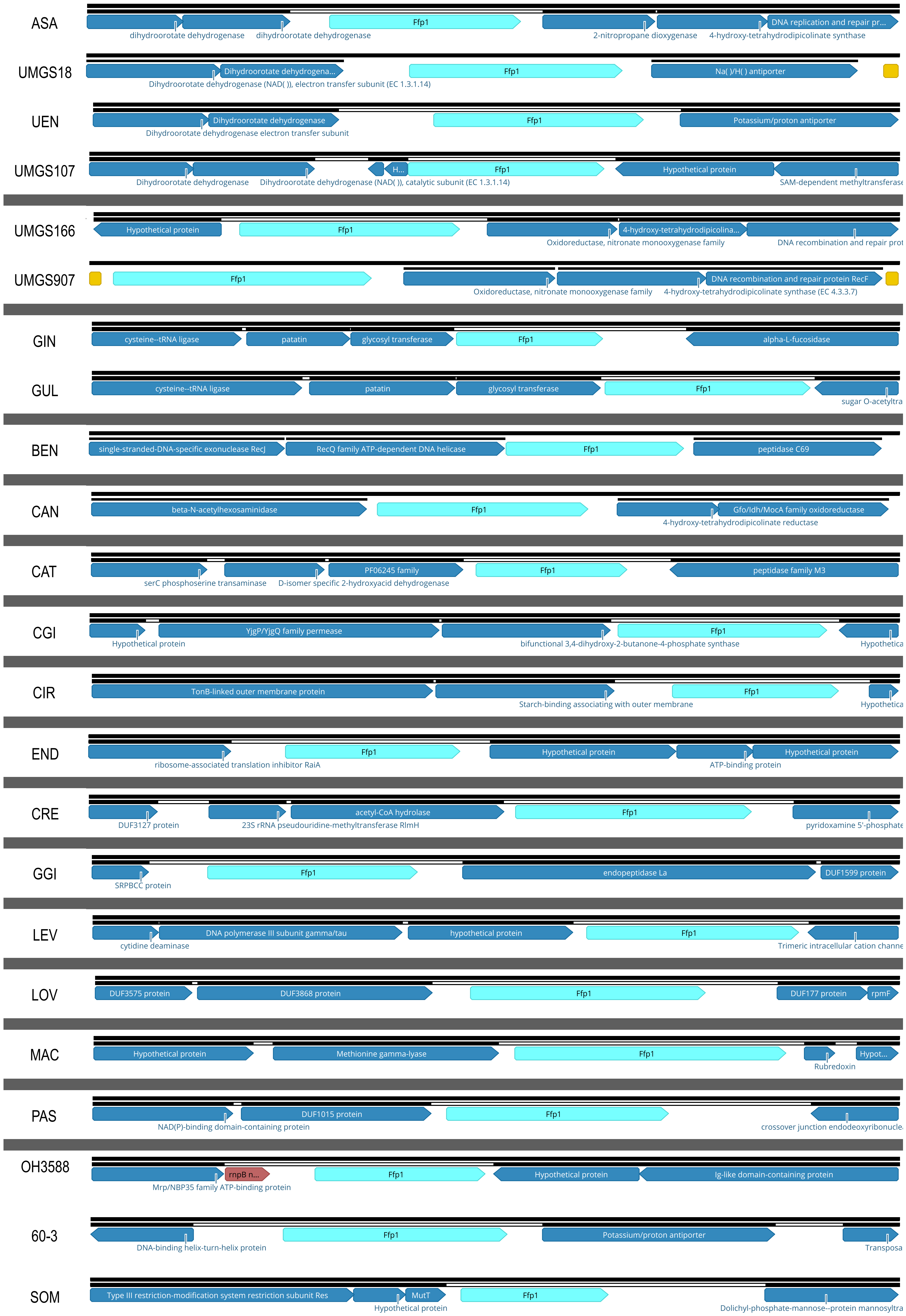
ASA
dihydroorotate dehydrogenase
dihydroorotate dehydrogenase
Ffp1
2-nitropropane dioxygenase
4-hydroxy-tetrahydrodipicolinate synthase
DNA replication and repair pr...
UMGS18
Dihydroorotate dehydrogena...
Dihydroorotate dehydrogenase (NAD( )), electron transfer subunit (EC 1.3.1.14)
Ffp1
Na( )/H( ) antiporter
UEN
Dihydroorotate dehydrogenase
Dihydroorotate dehydrogenase electron transfer subunit
Ffp1
Potassium/proton antiporter
UMGS107
Dihydroorotate dehydrogenase
Dihydroorotate dehydrogenase (NAD( )), catalytic subunit (EC 1.3.1.14)
H...
Ffp1
Hypothetical protein
SAM-dependent methyltransferase
UMGS166
Hypothetical protein
Ffp1
4-hydroxy-tetrahydrodipicolina...
Oxidoreductase, nitronate monooxygenase family
DNA recombination and repair prot
UMGS907
Ffp1
DNA recombination and repair protein RecF
Oxidoreductase, nitronate monooxygenase family
4-hydroxy-tetrahydrodipicolinate synthase (EC 4.3.3.7)
GIN
cysteine--tRNA ligase
patatin
glycosyl transferase
Ffp1
alpha-L-fucosidase
GUL
cysteine--tRNA ligase
patatin
glycosyl transferase
Ffp1
sugar O-acetyltra
BEN
single-stranded-DNA-specific exonuclease RecJ
RecQ family ATP-dependent DNA helicase
Ffp1
peptidase C69
CAN
beta-N-acetylhexosaminidase
Ffp1
Gfo/Idh/MocA family oxidoreductase
4-hydroxy-tetrahydrodipicolinate reductase
CAT
PF06245 family
Ffp1
peptidase family M3
serC phosphoserine transaminase
D-isomer specific 2-hydroxyacid dehydrogenase
CGI
YjgP/YjgQ family permease
Ffp1
Hypothetical protein
bifunctional 3,4-dihydroxy-2-butanone-4-phosphate synthase
Hypothetica
CIR
TonB-linked outer membrane protein
Ffp1
Starch-binding associating with outer membrane
Hypothetica
END
Ffp1
Hypothetical protein
Hypothetical protein
ribosome-associated translation inhibitor RaiA
ATP-binding protein
CRE
acetyl-CoA hydrolase
Ffp1
DUF3127 protein
23S rRNA pseudouridine-methyltransferase RlmH
pyridoxamine 5'-phosphate
GGI
Ffp1
endopeptidase La
DUF1599 protein
SRPBCC protein
LEV
DNA polymerase III subunit gamma/tau
hypothetical protein
Ffp1
cytidine deaminase
Trimeric intracellular cation channe
LOV
DUF3575 protein
DUF3868 protein
Ffp1
DUF177 protein
rpmF
MAC
Hypothetical protein
Methionine gamma-lyase
Ffp1
Hypot...
Rubredoxin
PAS
DUF1015 protein
Ffp1
NAD(P)-binding domain-containing protein
crossover junction endodeoxyribonucle
OH3588
rnpB n...
Ffp1
Hypothetical protein
Ig-like domain-containing protein
Mrp/NBP35 family ATP-binding protein
60-3
Ffp1
Potassium/proton antiporter
DNA-binding helix-turn-helix protein
Transposa
SOM
Type III restriction-modification system restriction subunit Res
MutT
Ffp1
Hypothetical protein
Dolichyl-phosphate-mannose--protein mannosyltra

| List of *Porphyromonas* (and related information) used in this study | | | | | | |
|---|---|---|---|---|---|---|
| **Species** | **Ref** | **Strain** | **Group acronym (this study)** | **Acronym (this study)** | **Genome Status (nb contigs)** | **Accession (WGS record)** |
| *asaccharolytica* | | PR42 6713P-I | ASA | ASA_PR426713P-I | Draft (58) | AENO00000000 |
| *asaccharolytica* | X | DSM 20707 | ASA | ASA_DSM 20707 | Complete (1) | NC_015501.1 |
| *sp.* | | MGYG-HGUT-04267 | ASA | ASA_MGYG-HGUT-04267 | Draft (74) | CABPCR010000001-07 |
| *bennonis* | X | DSM 23058 | BEN | BEN_DSM 23058 | Draft (87) | AQWR00000000 |
| *bennonis* | | JCM 16335 | BEN | BEN_JCM 16335 | Draft (242) | BAME00000000 |
| *canoris* | | COT-108 OH1224 | CAN | CAN_COT-108 OH1224 | Draft (21) | JQZX00000000 |
| *canoris* | X | COT-108 OH2762 | CAN | CAN_OH2762 | Draft (14) | JQZV00000000 |
| *sp.* | | COT-108_OH1349 | CAN | CAN_COT-108_OH1349 | Draft (43) | JRAH00000000 |
| *sp.* | | COT-108_OH2963 | CAN | CAN_COT-108_OH2963 | Draft (21) | JRAP00000000 |
| *catoniae* | | ATCC 51270 | CAT | CAT_ATCC 51270 | Draft (25) | JDFF00000000 |
| *catoniae* | X | F0037 | CAT | CAT_F0037 | Draft (18) | AMEQ00000000 |
| *cangingivalis* | | ATCC 700135 | CGI | CGI_ATCC 700135 | Draft (34) | FUWL00000000 |
| *cangingivalis* | | COT-109 OH1379 | CGI | CGI_OH1379 | Draft (21) | JQJF00000000 |
| *cangingivalis* | | COT-109 OH1386 | CGI | CGI_OH1386 | Draft (65) | JQJD00000000 |
| *cangingivalis* | | JCM 15983 | CGI | CGI_JCM 15983 | Draft (48) | BAKR00000000 |
| *cangingivalis* | X | NCTC 12856 | CGI | CGI_NCTC12856 | Complete (1) | LR134506 |
| *cangingivalis* | | NCTC 12857 | CGI | CGI_NCTC12857 | Draft (14) | UATO01000000 |
| *circumdentaria* | X | DSM 13022 | CIR | CIR_DSM_103022 | Draft (30) | JACIJQ010000001-07 |
| *circumdentaria* | | ATCC 51356 | CIR | CIR_ATCC 51356 | Draft (35) | FUXE00000000 |
| *crevioricanis* | | ATCC 55563 | CRE | CRE_ATCC 55563 | Draft (29) | FUXH00000000 |
| *crevioricanis* | | COT-253_OH1447 | CRE | CRE_OH1447 | Draft (30) | JQJC00000000 |
| *crevioricanis* | | COT-253_OH2125 | CRE | CRE_OH2125 | Draft (14) | JQJB00000000 |
| *crevioricani* | | JCM 13913 | CRE | CRE_JCM 13913 | Draft (89) | BAOV00000000 |
| *crevioricanis* | | JCM 15906 | CRE | CRE_JCM 15906 | Draft (118) | BAOU00000000 |
| *crevioricanis* | X | NCTC12858 | CRE | CRE_NCTC12858 | Complete (1) | LS483447 |
| *endodontalis* | | ATCC 35406 | END | END_ATCC 35406 | Draft (37) | ACNN00000000 |
| *endodontalis* | | NCTC13058 | END | END_NCTC13058 | Draft (2) | UGTE01000000 |
| *endodontalis* | X | FDAARGOS_1506 | END | END_FDAARGOS_1506 | Complete (1) | ASM2009737 |
| *gingivicanis* | X | COT-022 OH1391 | GGI | GGI_JCM 15907 | Draft (19) | JQZW00000000 |

List of *Porphyromonas* (and related information) used in this study

| Species | Ref | Strain | Group acronym (this study) | Acronym (this study) | Genome Status (nb contigs) | Accession (WGS record) |
|---|---|---|---|---|---|---|
| *gingivicanis* | | JCM 1597 | GGI | GGI_OH1391 | Draft (35) | BAKX00000000 |
| *gingivalis* | | 3_3 | GIN | GIN_3_3 | Draft (72) | FUFB00000000 |
| *gingivalis* | | 3A1 | GIN | GIN_3A1 | Draft (56) | FUFC00000000 |
| *gingivalis* | | 7BTORR | GIN | GIN_7BTORR | Draft (72) | FUFD00000000 |
| *gingivalis* | | 11A | GIN | GIN_11A | Draft (89) | FUFE00000000 |
| *gingivalis* | | 13_1 | GIN | GIN_13-1 | Draft (68) | FUGG00000000 |
| *gingivalis* | | 15_9 | GIN | GIN_15_9 | Draft (68) | FUGF00000000 |
| *gingivalis* | | 84_3 | GIN | GIN_84-3 | Draft (50) | FUFG00000000 |
| *gingivalis* | | 381 | GIN | GIN_381 | Complete (1) | CP012889 |
| *gingivalis* | | 381OKJP | GIN | GIN_381OKJP | Draft (127) | QPGS01000000 |
| *gingivalis* | | A7A1-28 | GIN | GIN_A7A1-28 | Complete (1) | CP013131 |
| *gingivalis* | | A7436 | GIN | GIN_A7436 | Complete (1) | CP011995 |
| *gingivalis* | | AFR5B1 | GIN | GIN_AFR5B1 | Draft (88) | FUFJ00000000 |
| *gingivalis* | | AJW4 | GIN | GIN_AJW4 | Complete (1) | CP011996 |
| *gingivalis* | | Ando | GIN | GIN_Ando | Draft (112) | BCBV01000000 |
| *gingivalis* | X | ATCC 33277 | GIN | GIN_ATCC 33277 | Complete (1) | CP025930/AP009380 |
| *gingivalis* | | ATCC 49417 | GIN | GIN_ATCC 49417 | Draft (77) | FUFH00000000 |
| *gingivalis* | | CP3 | GIN | GIN_CP3 | Draft (118) | SGBA01000000 |
| *gingivalis* | | F0185 | GIN | GIN_F0185 | Draft (113) | AWVC00000000 |
| *gingivalis* | | F0566 | GIN | GIN_F0566 | Draft (192) | AWVD00000000 |
| *gingivalis* | | F0568 | GIN | GIN_F0568 | Draft (154) | AWUU00000000 |
| *gingivalis* | | F0569 | GIN | GIN_F0569 | Draft (111) | AWUV00000000 |
| *gingivalis* | | F0570 | GIN | GIN_F0570 | Draft (117) | AWUW00000000 |
| *gingivalis* | | H3 | GIN | GIN_H3 | Draft (165) | SGAZ01000000 |
| *gingivalis* | | HG66 | GIN | GIN_HG66 | Complete (1) | CP007756.1 |
| *gingivalis* | | JCVI SC001 | GIN | GIN_JCVI SC001 | Draft (282 gaps) | APMB01000000 |
| *gingivalis* | | KCOM 2796 | GIN | GIN_KCOM 2796 | Complete (1) | CP024597 |
| *gingivalis* | | KCOM 2797 | GIN | GIN_KCOM 2797 | Draft (44) | NHRU00000000 |
| *gingivalis* | | KCOM 2798 | GIN | GIN_KCOM 2798 | Complete (1) | CP024598 |
| *gingivalis* | | KCOM 2799 | GIN | GIN_KCOM 2799 | Complete (1) | CP024601 |
| *gingivalis* | | KCOM 2800 | GIN | GIN_KCOM 2800 | Complete (1) | CP024599 |

| List of *Porphyromonas* (and related information) used in this study | | | | | | |
|---|---|---|---|---|---|---|
| **Species** | **Ref** | **Strain** | **Group acronym (this study)** | **Acronym (this study)** | **Genome Status (nb contigs)** | **Accession (WGS record)** |
| *gingivalis* | | KCOM 2801 | GIN | GIN_KCOM 2801 | Complete (1) | CP024600 |
| *gingivalis* | | KCOM 2802 | GIN | GIN_KCOM 2802 | Complete (1) | CP024591 |
| *gingivalis* | | KCOM 2803 | GIN | GIN_KCOM 2803 | Complete (1) | CP024592 |
| *gingivalis* | | KCOM 2804 | GIN | GIN_KCOM 2804 | Complete (1) | CP024593 |
| *gingivalis* | | KCOM 2805 | GIN | GIN_KCOM 2805 | Complete (1) | CP024594 |
| *gingivalis* | | KCOM 3001 | GIN | GIN_KCOM 3001 | Complete (1) | CP024595 |
| *gingivalis* | | KCOM 3131 | GIN | GIN_KCOM 3131 | Complete (1) | CP024596 |
| *gingivalis* | | MP4-504 | GIN | GIN_MP4-504 | Draft (92) | LOEL00000000 |
| *gingivalis* | | SJD2 | GIN | GIN_SJD2 | Draft (117) | ASYL00000000 |
| *gingivalis* | | SJD4 | GIN | GIN_SJD4 | Draft (147) | KZ248242-KZ248388 |
| *gingivalis* | | SJD5 | GIN | GIN_SJD5 | Draft (194) | ASYN00000000 |
| *gingivalis* | | SJD11 | GIN | GIN_SJD11 | Draft (156) | KZ248389-KZ248544 |
| *gingivalis* | | SJD12 | GIN | GIN_SJD12 | Draft (146) | KZ253969-KZ254114 |
| *gingivalis* | | SU60 | GIN | GIN_SU60 | Draft (53) | FUFI00000000 |
| *gingivalis* | | TDC60 | GIN | GIN_TDC60 | Complete (1) | CP025931/AP012203 |
| *gingivalis* | | W50 | GIN | GIN_W50 | Draft (104) | AJZS00000000 |
| *gingivalis* | | W83 | GIN | GIN_W83 | Complete (1) | CP025932/AE015924 |
| *gingivalis* | | W4087 | GIN | GIN_W4087 | Draft (114) | AWVE00000000 |
| *gingivalis* | | WW2096 | GIN | GIN_WW2096 | Draft (116) | NSLX01000000 |
| *gingivalis* | | WW2842 | GIN | GIN_WW2842 | Draft (95) | NSLW01000000 |
| *gingivalis* | | WW2866 | GIN | GIN_WW2866 | Draft (123) | NSLV01000000 |
| *gingivalis* | | WW2881 | GIN | GIN_WW2881 | Draft (484) | NSLU01000000 |
| *gingivalis* | | WW2885 | GIN | GIN_WW2885 | Draft (196) | NSLT01000000 |
| *gingivalis* | | WW2903 | GIN | GIN_WW2903 | Draft (115) | NSLS01000000 |
| *gingivalis* | | WW2931 | GIN | GIN_WW2931 | Draft (103) | NSLR01000000 |
| *gingivalis* | | WW2952 | GIN | GIN_WW2952 | Draft (162) | NSLQ01000000 |
| *gingivalis* | | WW3039 | GIN | GIN_WW3039 | Draft (132) | NSLN01000000 |
| *gingivalis* | | WW3102 | GIN | GIN_WW3102 | Draft (149) | NSLO01000000 |
| *gingivalis* | | WW5127 | GIN | GIN_WW5127 | Draft (119) | NSLL01000000 |
| *gulae* | | COT-052_OH1355 | GUL | GUL_OH1355 | Draft (40) | JRAG00000000 |
| *gulae* | | COT-052_OH1451 | GUL | GUL_OH1451 | Draft (89) | JRAI00000000 |

| List of *Porphyromonas* (and related information) used in this study | | | | | | |
|---|---|---|---|---|---|---|
| Species | Ref | Strain | Group acronym (this study) | Acronym (this study) | Genome Status (nb contigs) | Accession (WGS record) |
| *gulae* | | COT-052_OH2179 | GUL | GUL_OH2179 | Draft (26) | JRAJ00000000 |
| *gulae* | | COT-052_OH2199 | GUL | GUL_OH2199 | Draft (92) | JRAE01000000 |
| *gulae* | | COT-052_OH2857 | GUL | GUL_OH2857 | Draft (53) | JRFD00000000 |
| *gulae* | | COT-052_OH3439 | GUL | GUL_OH3439 | Draft (163) | JRAK00000000 |
| *gulae* | | COT-052_OH3471 | GUL | GUL_OH3471 | Draft (44) | JRAQ00000000 |
| *gulae* | | COT-052_OH3498 | GUL | GUL_OH3498 | Draft (71) | JRAF00000000 |
| *gulae* | X | COT-052_OH3856 | GUL | GUL_OH3856 | Draft (31) | JRAT00000000 |
| *gulae* | | COT-052_OH4119 | GUL | GUL_OH4119 | Draft (52) | JRAL00000000 |
| *gulae* | | DSM 15663 | GUL | GUL_DSM 15663 | Draft (63) | ARJN00000000 |
| *gulae* | | OH3161B | GUL | GUL_OH3161B | Draft (47) | JQJE00000000 |
| *sp.* | | UQD_349_COT-052_OH4946 | GUL | GUL_OH4946 | Draft (34) | JQZY00000000 |
| *levii* | | AF0918 | LEV | LEV_AF0918 | Draft (364) | SPNB01000000 |
| *levii* | | AF5678 | LEV | LEV_AF5678 | Draft (468) | SPNC01000000 |
| *levii* | X | DSM 23370 | LEV | LEV_DSM 23370 | Draft (124) | ARBX00000000 |
| *loveana* | X | DSM 28520 | LOV | LOV_DSM 28520 | Draft (39) | QEKY01000000 |
| *macacae* | | COT-192_OH2631 | MAC | MAC_OH2631 | Draft (48) | JRFB00000000 |
| *macacae* | | COT-192_OH2859 | MAC | MAC_OH2859 | Draft (32) | JRFA00000000 |
| *macacae* | X | DSM 20710 | MAC | MAC_DSM 20710 | Draft (31) | ARBY00000000 |
| *macacae* | | JCM 13914 | MAC | MAC_JCM 13914 | Draft (44) | BAKQ00000000 |
| *macacae* | | JCM 15984 | MAC | MAC_JCM 15984 | Draft (69) | BAKS00000000 |
| *macacae* | | NCTC11632 | MAC | MAC_NCTC11632 | Draft (6) | UGTF01000000 |
| *macacae* | | NCTC13100 | MAC | MAC_NCTC13100 | Draft (5) | UGTI01000000 |
| *pasteri* | X | JCM 30531 | PAS | PAS_JCM 30531 | Draft (9) | ASM1464775 |
| *somerae* | | DSM 23387 = JCM 13868 | SOM | SOM_DSM_23387 | Draft (95) | AQVC00000000 |
| *somerae* | X | CE91-St14 | SOM | SOM_St14 | Complete (1) | AP025559 |
| *sp.* | | UMGS1452 | UEN | UEN_UMGS1452 | Draft (337) | URXQ01000000 |
| *uenonis* | X | DSM23387 | UEN | UEN_DSM23387 | Draft (44) | AXVC00000000 |
| *uenonis* | | JCM13868 | UEN | UEN_JCM13868 | Draft (112) | BAJM00000000 |
| *sp.* | | COT-290_OH3588 | no assignation | PSP_OH3588 | Draft (48) | JRFC00000000 |

| List of *Porphyromonas* (and related information) used in this study | | | | | | |
|---|---|---|---|---|---|---|
| **Species** | **Ref** | **Strain** | **Group acronym (this study)** | **Acronym (this study)** | **Genome Status (nb contigs)** | **Accession (WGS record)** |
| *sp.* | | UMGS18 | no assignation | PSP_UMGS18 | Draft (60) | UWSO01000000 |
| *sp.* | | UMGS107 | no assignation | PSP_UMGS107 | Draft (65) | UQBE01000000 |
| *sp.* | | UMGS166 | no assignation | PSP_UMGS166 | Draft (83) | UQDF01000000 |
| *sp.* | | UMGS907 | no assignation | PSP_UMGS907 | Draft (109) | URDQ01000000 |
| *sp.* | | UMGS1085 | no assignation | PSP_UMGS1085 | Draft (286) | URKO01000000 |
| *sp.* | | 60-3 | **not *uenonis*** | PSP_60-3 | Draft (250) | ACLR00000000 |
| *sp.* | | 31_2 | ***Parabacteroides*** | not considered | Draft (13) | ACUD00000000 |
| *sp.* | | bin_26 | ***Genomes mixture*** | not considered | Draft (328 | RBKG01000000 |
| *somerae* | | KA00683 | ***Genomes mixture*** | not considered | Draft (62) | LSDK00000000 |
| *sp.* | | CAG:1061 | ***Genomes mixture*** | not considered | Draft (344) | CAXO000000000 |
| *sp.* | | oral taxon 279 str. F0450 | ***Genomes mixture*** | not considered | Draft (51) | ALKJ00000000 |
| *sp.* | | HMSC065F10 | ***Genomes mixture*** | not considered | Draft (103) | LTXH00000000 |
| *sp.* | | HMSC077F02 | ***Genomes mixture*** | not considered | Draft (85) | LTSX00000000 |
| *sp.* | | KLE 1280 | ***Genomes mixture*** | not considered | Draft (8) | JNOS00000000 |
| *sp.* | | UMGS2020 | ***Genomes mixture*** | not considered | Draft (280) | USRR01000000 |
| *sp.* | | COT-290_OH860 | ***Genomes mixture*** | not considered | Draft (82) | JRAR00000000 |
| *sp.* | | COT-239_OH1446 | ***Genomes mixture*** | not considered | Draft (37) | JRAO00000000 |
| *sp.* | | UMGS547 | ***Genomes mixture*** | not considered | Draft (46) | UQQP01000000 |
| *sp.* | | UMGS338 | ***Genomes mixture*** | not considered | Draft (33) | UQJH01000000 |
| *sp.* | | UMGS713 | ***Genomes mixture*** | not considered | Draft (163) | UQWO01000000 |
| *sp.* | | UMGS1769 | ***Genomes mixture*** | not considered | Draft (183) | USJF01000000 |
| *sp.* | | UMGS2040 | ***Genomes mixture*** | not considered | Draft (238) | USSQ01000000 |
| *sp.* | | oral taxon 278 str. W7784 | ***Genomes mixture*** | not considered | Draft (38) | AWUX00000000 |
| *sp.* | | MGYG-HGUT-04270 | ***Genomes mixture*** | not considered | Draft (197) | CABPCS010000001-07 |

| Acronym (this study) | HMM groups | Locus_tag | Old_locus_tag | Initiation codon reannotation | SignalP-6/LipoP prediction before curation | SignalP-6/LipoP prediction after curation | Size before biocuration (in aa) | Size after biocuration (in aa) | Lipoprotein signal peptide (Sec/SPII) | Cleavage site between pos. | Size after clivage (in aa) |
|---|---|---|---|---|---|---|---|---|---|---|---|
| ASA_PR426713P-I | Ffp1_A | HMPREF9294_RS07130 | HMPREF9294_0374 | VTLRL… (position +38aa, 5' truncation) | Cytoplasmic | SpII | 543 | 506 | VTLRLISLVVCMSVLLLGS | 19 and 20: LGS-CR | 487 |
| ASA_DSM 20707 | Ffp1_A | PORAS_RS06500 | Poras_1299 | VTPRL… (position +49aa, 5' truncation) | Cytoplasmic | SpII | 553 | 505 | VTPRLISLVVCMSVLLLGS | 19 and 20: LGS-CR | 486 |
| BEN_DSM 23058 | Ffp1_B | B088_RS06165 | - | - | SpII | - | 525 | - | MKRLKSTLLASIALLAVG | 18 and 19:AVG-CS | 507 |
| BEN_JCM 16335 | Ffp1_B | fig\|1236537.5.peg.6498 | - | - | SpII | - | 517 | - | MKRLKSTLLASIALLAVG | 18 and 19:AVG-CS | 499 |
| CAN_COT-108 OH1224 | Ffp1_B | HQ29_RS00605 | HQ29_00650 | - | SpII | - | 527 | - | MRKSHISALVFALALFSSG | 19 and 20:SSG-CK | 508 |
| CAN_OH2762 | Ffp1_B | HQ43_RS08510 | HQ43_09055 | - | SpII | - | 527 | - | MRKSHISALVFALALFSSG | 19 and 20:SSG-CK | 508 |
| CAN_OH1349 | Ffp1_B | JT26_RS08750 | JT26_09370 | - | SpII | - | 527 | - | MRKSHISALVLALALFSSG | 19 and 20:SSG-CK | 508 |
| CAN_OH2963 | Ffp1_B | HQ39_00925 | - | - | SpII | - | 527 | - | MRKSHISALVFALALFSSG | 19 and 20:SSG-CK | 508 |
| CAT_ATCC 51270 | Ffp1_A | HMPREF0636_1008 | - | MKPS….(position -8 aa, 5' elongation) | Cytoplasmic | SpII | 467 | 475 | MKPSLSLIMVLALAFSS | 17 and 18: FSS-CT | 458 |
| CAT_F0037 | Ffp1_A | HMPREF9134_01065 | - | MKPS….(position -8 aa, 5' elongation) | Cytoplasmic | SpII | 467 | 475 | MKPSLSLIMVLALAFSS | 17 and 18: FSS-CT | 458 |
| CGI_ATCC 700135 | Ffp1_B | B5D10_RS02040 | SAMN02745205_00414 | - | SpII | - | 501 | - | MKRPIKGLAILILLCVLVA | 19 and 20:LVA-CN | 482 |
| CGI_OH1379 | Ffp1_B | HQ34_RS02105 | HQ34_02210 | - | SpII | - | 501 | - | MKRPIKGLTILVLLCVLVA | 19 and 20:LVA-CN | 482 |
| CGI_OH1386 | Ffp1_B | HQ35_RS07210 | HQ35_07500 | - | SpII | - | 501 | - | MKRPIKGLTILVLLCVLVA | 19 and 20:LVA-CN | 482 |
| CGI_JCM 15983 | Ffp1_B | JCM15983_RS07445 | - | - | SpII | - | 501 | - | MKRPIKGLAILILLCVLVA | 19 and 20:LVA-CN | 482 |
| CGI_NCTC12856 | Ffp1_B | EL262_RS03910 | NCTC12856_00820 | - | SpII | - | 501 | - | MKRPIKGLAILILLCVLVA | 19 and 20:LVA-CN | 482 |
| CGI_NCTC12857 | Ffp1_B | NCTC12857_00769 | - | - | SpII | - | 501 | - | MKRPIKGLAILILLCVLVA | 19 and 20:LVA-CN | 482 |
| CIR_DSM_103022 | Ffp1_A | Ga0373368_1702 | - | LFFH …(position -55 aa, 5' elongation) | Cytoplasmic | SpII (best score) | 439 | 493 | LFFHIPLLSLFILSLLS | 17 and 18:LLS-CT | 476 |
| CIR_ATCC 51356 | Ffp1_A | SAMN02745171_01593 | - | LFFH …(position +4 aa, 5' truncation) | SpII | SpII (best score) | 497 | 493 | LFFHIPLLSLFILSLLS | 17 and 18:LLS-CT | 476 |
| CRE_ATCC 55563 | Ffp1_A | B5D48_RS07615 | SAMN02745203_01515 | LFRI …(position +4 aa, 5' truncation) | SpII | SpII (best score) | 485 | - | LFRIWLLPVVFVLAMG | 16 and 17:AMG-CS | 465 |
| CRE_OH1447 | Ffp1_A | HQ38_00070 | - | LFRI …(position +4 aa, 5' truncation) | SpII | SpII (best score) | 485 | - | LFRIWLLPVVFVLAMG | 16 and 17:AMG-CS | 465 |
| CRE_OH2125 | Ffp1_A | HQ45_07910 | - | LFRI …(position +4 aa, 5' truncation) | SpII | SpII (best score) | 485 | - | LFRIWLLPVVFVLAMG | 16 and 17:AMG-CS | 465 |
| CRE_JCM 13913 | Ffp1_A | PORCAN_38 | - | LFRI …(position +4 aa, 5' truncation) | SpII | SpII (best score) | 485 | - | LFRIWLLPVVFVLAMG | 16 and 17:AMG-CS | 465 |
| CRE_JCM 15906 | Ffp1_A | TX94_RS03855 | PORCRE_924 | LFRI …(position +4 aa, 5' truncation) | SpII | SpII (best score) | 485 | - | LFRIWLLPVVFVLAMG | 16 and 17:AMG-CS | 465 |
| CRE_NCTC12858 | Ffp1_A | DQN70_RS01465 | NCTC12858_00290 | LFRI …(position +4 aa, 5' truncation) | SpII | SpII (best score) | 485 | - | LFRIWLLPVVFVLAMG | 16 and 17:AMG-CS | 465 |
| END_ATCC 35406 | Ffp1_A | POREN0001_RS08405 | POREN0001_0653 | - | SpII | - | 490 | - | MKSTAKILFGGLFMGVLLLA | 20 and 21:LLA-CH | 470 |
| END_NCTC13058 | Ffp1_A | NCTC13058_00879 | - | - | SpII | - | 490 | - | MKSTAKILFGGLFMGVLLLA | 20 and 21:LLA-CH | 470 |
| END_FDAARGOS_1506 | Ffp1_A | LA319_RS03360 | LA319_03360 | - | SpII | - | 490 | - | MKSTAKILFGGLFMGVLLLA | 20 and 21:LLA-CH | 470 |
| GGI_JCM 15907 | Ffp1_A | JCM15907_RS02095 | - | LAYR…(position +4 aa, 5' truncation) | SpII | - | 495 | 491 | LAYRFISILLFISFLGS | 17 and 18:LGS-CK | 474 |
| GGI_OH1391 | Ffp1_A | HQ36_RS03135 | HQ36_03345 | LAYR…(position +4 aa, 5' truncation) | SpII | - | 495 | 491 | LAYRFISILLFISFLGS | 17 and 18:LGS-CK | 474 |
| GIN_3_3 | Ffp1_A | PGIN_3-3_01611 | - | - | SpII | - | 481 | - | MLTKLKTLLLGCSLACIGFS | 20 and 21: GFS-CS | 461 |
| GIN_3A1 | Ffp1_A | B0496_RS06505 | PGIN_3A1_01536 | - | SpII | - | 481 | - | MLTKLKTLLLGCSLACIGFS | 20 and 21: GFS-CS | 461 |
| GIN_7BTORR | Ffp1_A | PGIN_7BTORR_01660 | - | - | SpII | - | 481 | - | MLTKLKTLLLGCSLACIGFS | 20 and 21: GFS-CS | 461 |
| GIN_11A | Ffp1_A | PGIN_11A_01261 | - | - | SpII | - | 481 | - | MLTKLKTLLLGCSLACIGFS | 20 and 21: GFS-CS | 461 |
| GIN_13-1 | Ffp1_A | PGIN_13-1_00894 | - | - | SpII | - | 481 | - | MLTKLKTLLLGCSLACIGFS | 20 and 21: GFS-CS | 461 |
| GIN_15_9 | Ffp1_A | CEP61_RS05480 | PGIN_15-9_01143 | - | SpII | - | 481 | - | MLTKLKTLLLGCSLACIGFS | 20 and 21: GFS-CS | 461 |
| GIN_84-3 | Ffp1_A | PGIN_84-3_01869 | - | - | SpII | - | 481 | - | MLTKLKTLLLGCSLACIGFS | 20 and 21: GFS-CS | 461 |
| GIN_381 | Ffp1_A | PGF_RS08670 | PGF_00017830 | - | SpII | - | 481 | - | MLTKLKTLLLGCSLACIGFS | 20 and 21: GFS-CS | 461 |
| GIN_381OKJP | Ffp1_A | DOE52_RS07275 | DOE52_07370 | - | SpII | - | 481 | - | MLTKLKTLLLGCSLACIGFS | 20 and 21: GFS-CS | 461 |
| GIN_A7A1-28 | Ffp1_A | PGS_00016750 | - | - | SpII | - | 481 | - | MLTKLKTLLLGCSLACIGFS | 20 and 21: GFS-CS | 461 |
| GIN_A7436 | Ffp1_A | PGA7_RS08340 | PGA7_00017440 | - | SpII | - | 481 | - | MLTKLKTLLLGCSLACIGFS | 20 and 21: GFS-CS | 461 |
| GIN_AFR5B1 | Ffp1_A | PGIN_AFR-5B1_00930 | - | - | SpII | - | 481 | - | MLTKLKTLLLGCSLACIGFS | 20 and 21: GFS-CS | 461 |
| GIN_AJW4 | Ffp1_A | PGJ_RS08140 | PGJ_00016930 | - | SpII | - | 481 | - | MLTKLKTLLLGCSLACIGFS | 20 and 21: GFS-CS | 461 |
| GIN_Ando | Ffp1_A | PGANDO_1295 | - | - | SpII | - | 481 | - | MLTKLKTLLLGCSLACIGFS | 20 and 21: GFS-CS | 461 |
| GIN_ATCC 33277 | Ffp1_A | PGN_RS08575 | PGN_1808 | - | SpII | - | 481 | - | MLTKLKTLLLGCSLACIGFS | 20 and 21: GFS-CS | 461 |
| GIN_ATCC 49417 | Ffp1_A | PGIN_ATCC49417_00365 | - | - | SpII | - | 481 | - | MLTKLKTLLLGCSLACIGFS | 20 and 21: GFS-CS | 461 |
| GIN_CP3 | Ffp1_A | EW639_RS04185 | EW639_04180 | - | SpII | - | 481 | - | MLTKLKTLLLGCSLACIGFS | 20 and 21: GFS-CS | 461 |
| GIN_F0185 | Ffp1_A | HMPREF1988_00474 | - | - | SpII | - | 481 | - | MLTKLKTLLLGCSLACIGFS | 20 and 21: GFS-CS | 461 |
| GIN_F0566 | Ffp1_A | HMPREF1989_01686 | - | - | SpII | - | 481 | - | MLTKLKTLLLGCSLACIGFS | 20 and 21: GFS-CS | 461 |
| GIN_F0568 | Ffp1_A | HMPREF1553_02200 | - | - | SpII | - | 481 | - | MLTKLKTLLLGCSLACIGFS | 20 and 21: GFS-CS | 461 |
| GIN_F0569 | Ffp1_A | HMPREF1554_01931 | - | - | SpII | - | 481 | - | MLTKLKTLLLGCSLACIGFS | 20 and 21: GFS-CS | 461 |
| GIN_F0570 | Ffp1_A | HMPREF1555_00703 | - | - | SpII | - | 481 | - | MLTKLKTLLLGCSLACIGFS | 20 and 21: GFS-CS | 461 |
| GIN_H3 | Ffp1_A | EW638_RS01795 | EW638_01795 | - | SpII | - | 481 | - | MLTKLKTLLLGCSLACIGFS | 20 and 21: GFS-CS | 461 |
| GIN_HG66 | Ffp1_A | EG14_RS00370 | EG14_00390 | - | SpII | - | 481 | - | MLTKLKTLLLGCSLACIGFS | 20 and 21: GFS-CS | 461 |
| GIN_JCVI SC001 | Ffp1_A | A343_0931 | - | - | SpII | - | 481 | - | MLTKLKTLLLGCSLACIGFS | 20 and 21: GFS-CS | 461 |
| GIN_KCOM 2796 | Ffp1_A | CS059_RS02825 | CS059_02815 | - | SpII | - | 481 | - | MLTKLKTLLLGCSLACIGFS | 20 and 21: GFS-CS | 461 |
| GIN_KCOM 2797 | Ffp1_A | CBG53_RS04765 | CBG53_04755 | - | SpII | - | 481 | - | MLTKLKTLLLGCSLACIGFS | 20 and 21: GFS-CS | 461 |
| GIN_KCOM 2798 | Ffp1_A | CS374_RS06220 | CS374_06175 | - | SpII | - | 481 | - | MLTKLKTLLLGCSLACIGFS | 20 and 21: GFS-CS | 461 |
| GIN_KCOM 2799 | Ffp1_A | CS387_RS05960 | CS387_05925 | - | SpII | - | 481 | - | MLTKLKTLLLGCSLACIGFS | 20 and 21: GFS-CS | 461 |
| GIN_KCOM 2800 | Ffp1_A | CS388_RS06295 | CS388_06250 | - | SpII | - | 481 | - | MLTKLKTLLLGCSLACIGFS | 20 and 21: GFS-CS | 461 |
| GIN_KCOM 2801 | Ffp1_A | CS543_RS02035 | CS543_02040 | - | SpII | - | 481 | - | MLTKLKTLLLGCSLACIGFS | 20 and 21: GFS-CS | 461 |
| GIN_KCOM 2802 | Ffp1_A | CS544_RS06860 | CS544_06820 | - | SpII | - | 481 | - | MLTKLKTLLLGCSLACIGFS | 20 and 21: GFS-CS | 461 |
| GIN_KCOM 2803 | Ffp1_A | CS545_RS01100 | CS545_01090 | - | SpII | - | 481 | - | MLTKLKTLLLGCSLACIGFS | 20 and 21: GFS-CS | 461 |
| GIN_KCOM 2804 | Ffp1_A | CS546_RS04285 | CS546_04255 | - | SpII | - | 481 | - | MLTKLKTLLLGCSLACIGFS | 20 and 21: GFS-CS | 461 |
| GIN_KCOM 2805 | Ffp1_A | CS548_RS00115 | CS548_00110 | - | SpII | - | 481 | - | MLTKLKTLLLGCSLACIGFS | 20 and 21: GFS-CS | 461 |
| GIN_KCOM 3001 | Ffp1_A | CS550_RS00280 | CS550_00280 | - | SpII | - | 481 | - | MLTKLKTLLLGCSLACIGFS | 20 and 21: GFS-CS | 461 |
| GIN_KCOM 3131 | Ffp1_A | CS549_RS03615 | CS549_03580 | - | SpII | - | 481 | - | MLTKLKTLLLGCSLACIGFS | 20 and 21: GFS-CS | 461 |
| GIN_MP4-504 | Ffp1_A | AT291_01865 | - | - | SpII | - | 481 | - | MLTKLKTLLLGCSLACIGFS | 20 and 21: GFS-CS | 461 |
| GIN_SJD2 | Ffp1_A | SJDPG2_RS03990 | SJDPG2_06325 | - | SpII | - | 481 | - | MLTKLKTLLLGCSLACIGFS | 20 and 21: GFS-CS | 461 |
| GIN_SJD4 | Ffp1_A | SJDPG4_RS07350 | SJDPG4_07395 | - | SpII | - | 481 | - | MLTKLKTLLLGCSLACIGFS | 20 and 21: GFS-CS | 461 |
| GIN_SJD5 | Ffp1_A | SJDPG5_RS08005 | SJDPG5_07225 | - | SpII | - | 481 | - | MLTKLKTLLLGCSLACIGFS | 20 and 21: GFS-CS | 461 |

| Acronym (this study) | HMM groups | Locus_tag | Old_locus_tag | Initiation codon reannotation | SignalP-6/LipoP prediction before curation | SignalP-6/LipoP prediction after curation | Size before biocuration (in aa) | Size after biocuration (in aa) | Lipoprotein signal peptide (Sec/SPII) | Cleavage site between pos. | Size after clivage (in aa) |
|---|---|---|---|---|---|---|---|---|---|---|---|
| GIN_SJD11 | Ffp1_A | SJDPG11_RS03850 | SJDPG11_03950 | - | SpII | - | 481 | - | MLTKLKTLLLGCSLACIGFS | 20 and 21: GFS-CS | 461 |
| GIN_SJD12 | Ffp1_A | SJDPG12_RS04055 | SJDPG12_04100 | - | SpII | - | 481 | - | MLTKLKTLLLGCSLACIGFS | 20 and 21: GFS-CS | 461 |
| GIN_SU60 | Ffp1_A | PGIN_YH522_00088 | - | - | SpII | - | 481 | - | MLTKLKTLLLGCSLACIGFS | 20 and 21: GFS-CS | 461 |
| GIN_TDC60 | Ffp1_A | PGTDC60_RS00615 | PGTDC60_0141 | - | SpII | - | 481 | - | MLTKLKTLLLGCSLACIGFS | 20 and 21: GFS-CS | 461 |
| GIN_W50 | Ffp1_A | HMPREF1322_RS07675 | HMPREF1322_0808 | - | SpII | - | 481 | - | MLTKLKTLLLGCSLACIGFS | 20 and 21: GFS-CS | 461 |
| GIN_W83 | Ffp1_A | CF003_RS19550 | CF003_1881 | - | SpII | - | 481 | - | MLTKLKTLLLGCSLACIGFS | 20 and 21: GFS-CS | 461 |
| GIN_W4087 | Ffp1_A | HMPREF1990_01936 | - | - | SpII | - | 481 | - | MLTKLKTLLLGCSLACIGFS | 20 and 21: GFS-CS | 461 |
| GIN_WW2096 | Ffp1_A | CLI72_04935 | - | - | SpII | - | 481 | - | MLTKLKTLLLGCSLACIGFS | 20 and 21: GFS-CS | 461 |
| GIN_WW2842 | Ffp1_A | CLI73_05985 | - | - | SpII | - | 481 | - | MLTKLKTLLLGCSLACIGFS | 20 and 21: GFS-CS | 461 |
| GIN_WW2866 | Ffp1_A | CLI74_08035 | - | - | SpII | - | 481 | - | MLTKLKTLLLGCSLACIGFS | 20 and 21: GFS-CS | 461 |
| GIN_WW2881 | Ffp1_A | CLI75_RS00850 | CLI75_00850 | - | SpII | - | 481 | - | MLTKLKTLLLGCSLACIGFS | 20 and 21: GFS-CS | 461 |
| GIN_WW2885 | Ffp1_A | CLI76_07255 | - | - | SpII | - | 481 | - | MLTKLKTLLLGCSLACIGFS | 20 and 21: GFS-CS | 461 |
| GIN_WW2903 | Ffp1_A | CLI77_RS05155 | CLI77_05155 | - | SpII | - | 481 | - | MLTKLKTLLLGCSLACIGFS | 20 and 21: GFS-CS | 461 |
| GIN_WW2931 | Ffp1_A | CLI78_RS08380 | CLI78_08375 | - | SpII | - | 481 | - | MLTKLKTLLLGCSLACIGFS | 20 and 21: GFS-CS | 461 |
| GIN_WW2952 | Ffp1_A | CLI79_07275 | - | - | SpII | - | 481 | - | MLTKLKTLLLGCSLACIGFS | 20 and 21: GFS-CS | 461 |
| GIN_WW3039 | Ffp1_A | CLI82_RS08915 | CLI82_08920 | - | SpII | - | 481 | - | MLTKLKTLLLGCSLACIGFS | 20 and 21: GFS-CS | 461 |
| GIN_WW3102 | Ffp1_A | CLI81_06020 | - | - | SpII | - | 481 | - | MLTKLKTLLLGCSLACIGFS | 20 and 21: GFS-CS | 461 |
| GIN_WW5127 | Ffp1_A | CLI84_RS06820 | CLI84_06820 | - | SpII | - | 481 | - | MLTKLKTLLLGCSLACIGFS | 20 and 21: GFS-CS | 461 |
| GUL_OH1355 | Ffp1_A | HQ42_RS07930 | HQ42_08260 | - | SpII | - | 481 | - | MLTKLKSLLLGCSFACVGFS | 20 and 21: GFS-CS | 461 |
| GUL_OH1451 | Ffp1_A | HR08_RS10055 | HR08_10540 | - | SpII | - | 481 | - | MLMKLKSLLLGCSLACIGFS | 20 and 21: GFS-CS | 461 |
| GUL_OH2179 | Ffp1_A | HR09_01945 | - | - | SpII | - | 481 | - | MLTKLKSLLLGCSFACVGFS | 20 and 21: GFS-CS | 461 |
| GUL_OH2199 | Ffp1_A | HR10_RS02290 | HR10_02360 | - | SpII | - | 481 | - | MLTKLKSLLLGCSFACVGFS | 20 and 21: GFS-CS | 461 |
| GUL_OH2857 | Ffp1_A | HQ46_RS03870 | HQ46_04080 | - | SpII | - | 481 | - | MLTKLKSLLLGCSLACIGFS | 20 and 21: GFS-CS | 461 |
| GUL_OH3439 | Ffp1_A | HR15_RS03480 | HR15_03625 | - | SpII | - | 481 | - | MLMKLKSLLLGCSLACIGFS | 20 and 21: GFS-CS | 461 |
| GUL_OH3471 | Ffp1_A | HQ40_RS09880 | HQ40_10345 | - | SpII | - | 481 | - | MLMKLKSLLLGCSLACIGFS | 20 and 21: GFS-CS | 461 |
| GUL_OH3498 | Ffp1_A | HR16_RS01285 | HR16_01335 | - | SpII | - | 481 | - | MLTKLKSLLLGCSLACIGFS | 20 and 21: GFS-CS | 461 |
| GUL_OH3856 | Ffp1_A | HQ49_05395 | - | - | SpII | - | 481 | - | MLMKLKSLLLGCSLACIGFS | 20 and 21: GFS-CS | 461 |
| GUL_OH4119 | Ffp1_A | HR17_RS03350 | HR17_03500 | - | SpII | - | 481 | - | MLTKLKSLLLGCSLACIGFS | 20 and 21: GFS-CS | 461 |
| GUL_DSM 15663 | Ffp1_A | F452_RS0101570 | - | - | SpII | - | 481 | - | MLTKLKSLLLGCSFACVGFS | 20 and 21: GFS-CS | 461 |
| GUL_OH3161B | Ffp1_A | HR13_RS00620 | HR13_00630 | - | SpII | - | 481 | - | MLTKLKSLLLGCSFACVGFS | 20 and 21: GFS-CS | 461 |
| GUL_OH4946 | Ffp1_A | HQ50_02910 | - | - | SpII | - | 481 | - | MLTKLKSLLLGCSLACIGFS | 20 and 21: GFS-CS | 461 |
| LEV_AF0918 | Ffp1_B | E4P48_RS01530 | E4P48_01535 | - | SpII | - | 514 | - | MKTHLLCAIALLFLLLG | 17 and 18:LLG-CN | 497 |
| LEV_AF5678 | Ffp1_B | E4P47_RS07065 | E4P47_07070 | - | SpII | - | 514 | - | MKTHLLCAIALLFLLLG | 17 and 18:LLG-CN | 497 |
| LEV_DSM 23370 | Ffp1_B | A3GG_RS0105690 | - | - | SpII | - | 514 | - | MKTHLLCAIALLFLLLG | 17 and 18:LLG-CN | 497 |
| LOV_DSM 28520 | Ffp1_A | C7382_RS00405 | C7382_10187 | - | SpII | - | 481 | - | MKLKLLFASLALMCLSIA | 18 and 19:SIA-CS | 463 |
| MAC_OH2631 | Ffp1_A | HR11_RS03275 | HR11_03565 | - | SpII | - | 479 | - | MKKHLLFVFTVFLIAISS | 18 and 19:ISS-CN | 461 |
| MAC_OH2859 | Ffp1_A | HQ47_RS07445 | HQ47_07890 | - | SpII | - | 479 | - | MKKHLLFVFTVFLIAISS | 18 and 19:ISS-CN | 461 |
| MAC_DSM 20710 | Ffp1_A | fig\|1122974.3.peg.681 | - | - | SpII | - | 479 | - | MKKHLLFVFALFLIAISS | 18 and 19:ISS-CN | 461 |
| MAC_JCM 13914 | Ffp1_A | fig\|1236519.3.peg.1431 | - | - | SpII | - | 479 | - | MKKHLLFVFALFLIAISS | 18 and 19:ISS-CN | 461 |
| MAC_JCM 15984 | Ffp1_A | JCM15984DRAFT_02252 | - | note : this ORF is 3' truncated (end of contig) | SpII | - | 464 | - | …AISS (contig extremity) | na | 460 |
| MAC_NCTC11632 | Ffp1_A | NCTC11632_00113 | - | - | SpII | - | 479 | - | MKKHLLFVFTVFLIAISS | 18 and 19:ISS-CN | 461 |
| MAC_NCTC13100 | Ffp1_A | DXZ22_RS03080 | NCTC13100_00629 | - | SpII | - | 479 | - | MKKHLLFVFTVFLIAISS | 18 and 19:ISS-CN | 461 |
| PAS_JCM 30531 | Ffp1_A | IEX77_RS07640 | GCM10007088_14930 | MRRA…(position +10aa, 5' truncation) | SpII | SpII (best score) | 486 | 476 | MRRALSLSLSLLLLAG | 16 and 17:LACG-CS | 460 |
| SOM_DSM_23387 | Ffp1_B | A3GK_RS10475 | - | note : this ORF is 3' truncated (end of contig) | SpII | - | 483 | - | MRRLLSIILLALIVSS | 16 and 17:VSS-CN | 467… |
| SOM_St14 | Ffp1_B | CE91St14_06390 | - | MRRL…(-20aa, 5' elongation) | Cytoplasmic | SpII | 484 | 505 | MRRLLSIILLALIVSS | 16 and 17:VSS-CN | 489 |
| PSP_60-3 | Ffp1_A | PORUE0001_RS06855 | PORUE0001_1094 | LATSK…(position +23, 5' truncation) | SpII | SpII (best score) | 528 | 506 | LATSKLISLILCMSVLLLGA | 20 and 21:LGA-CR | 486 |
| PSP_OH3588 | Ffp1_A | HQ48_RS05340 | HQ48_05560 | LTKR…(position +7, 5' truncated) | SpII | - | 481 | 474 | LTKRMVLAGMSIVVLAFA | 18 and 19:AFA-CK | 456 |
| PSP_UMGS18 | Ffp1_A | fig\|159274.3.peg.1457 | - | LISL…(position +28, 5' truncation) | SpII | SpII (best score) | 529 | 501 | LISLVLCMSVLLLGA | 15 and 16:LGA-CR | 486 |
| PSP_UMGS107 | Ffp1_A | fig\|159274.4.peg.1387 | - | LIPEL…(position -14 aa, ORF 5' elongation) | Other | SpII | 490 | 504 | LIPELSFALCISILMLGA | 18 and 19:LGA-CR | 486 |
| PSP_UMGS166 | Ffp1_A | fig\|159274.5.peg.636 | - | LKHAF…(position +31, 5' truncation) | Other | SpII | 540 | 509 | LKHAFTATPKLLSLVLCMSVLLLGA | 25 and 26:LGA-CR | 484 |
| PSP_UMGS907 | Ffp1_A | fig\|159274.9.peg.1443 | - | LKLTC…(position +35, 5' truncation) | Other | SpII | 545 | 511 | LKLTCTATARLIPLVLCTSILLLGA | 25 and 26:LGA-CR | 486 |
| UEN_UMGS1452 | Ffp1_A | fig\|159274.11.peg.121 | - | MPRLI…(position 24) | SpII | SpII (best score) | 493 | 503 | MPRLISLVLCMSVLLLGA | 18 and 19:LGA-CR | 485 |
| UEN_DSM23387 | Ffp1_A | L215_RS09265 | - | MPRLI…(position -10 aa, ORF 5' elongation) | Other | SpII | 493 | 503 | MPRLISLVLCMSVLLLGA | 18 and 19:LGA-CR | 485 |
| UEN_JCM13868 | Ffp1_A | JCM13868_RS05230 | - | MPRLI…(position -10 aa, ORF 5' elongation) | Other | SpII | 527 | 503 | MPRLISLVLCMSVLLLGA | 18 and 19:LGA-CR | 485 |
